# De novo Folding Mechanisms of Lasso Peptides

**DOI:** 10.64898/2026.03.30.715466

**Authors:** Song Yin, Xuenan Mi, Susanna E. Barrett, Douglas A. Mitchell, Diwakar Shukla

## Abstract

Lasso peptides adopt a distinctive [1]rotaxane conformation, yet the principles governing the folding of this kinetically trapped structure have remained elusive. Here, we integrated extensive molecular dynamics simulations and deep learning to elucidate the de novo folding mechanism of 20 lasso peptides lacking secondary post-translational modifications. We constructed Multi-Ensemble Markov Models for each lasso peptide and uncovered a universal uphill folding landscape with spontaneous folding probabilities consistently below 0.8%. Loop stability strongly correlated with folding propensity, and targeted experiments further validated that enhancing loop β-hairpin formation promotes folding of microcin J25, the well-studied lasso peptide extensively characterized as an in vitro model. Additionally, the substantial entropy cost opposed lasso peptide folding. Simulations mimicking enzymatic spatial confinement reduced this penalty and stabilize folding. Leveraging Variational AutoEncoder-based pathway clustering, we resolved distinct pathway channels and representative folding pathways. Together, these findings establish representative folding models and fundamental thermodynamic and kinetic principles for rational engineering of lasso peptides.

## Introduction

Lasso peptides represent one molecular class of ribosomally synthesized and post-translationally modified peptide (RiPP) natural products.^1^ Compared to the other ∼50 described classes of RiPPs, lasso peptides are distinguished by their [1]rotaxane shape, which confers exceptional stability against proteases and harsh environmental conditions.^2,3^ To form the knot, a linear peptide must first adopt a lasso-like “pre-folded” conformation (**Figure 1A**) where the peptide first bends to create a loop which positions the N-terminus for subsequent isopeptide bond formation with an adenylated acidic acceptor residue (Asp or Glu, typically located at position 7, 8, or 9 in the lasso peptide core sequence). The mature lasso peptide conformation is stabilized primarily by bulky plug residues that hinder unthreading through steric entrapment.^4^ The distinctive fold and globular shapes of lasso peptides give rise to their diverse biological activities. For example, the lasso peptides microcin J25 and capistruin exhibit antimicrobial activity by inhibiting bacterial RNA polymerase by blocking nucleoside triphosphate (NTP) entry into the active site.^5^ Lariocidin inhibits the ribosome and disrupts protein synthesis, thus suppressing bacterial growth.^6^ In addition to their antibacterial activities, other lasso peptides show activity as glucagon receptor antagonists (BI-32169)^7^ and endothelin receptor B antagonists (RES-701).^8^ They have also been shown to display antiviral (siamycin),^9^ and anticancer (chaxapeptin) properties.^10^

**Figure 1.**
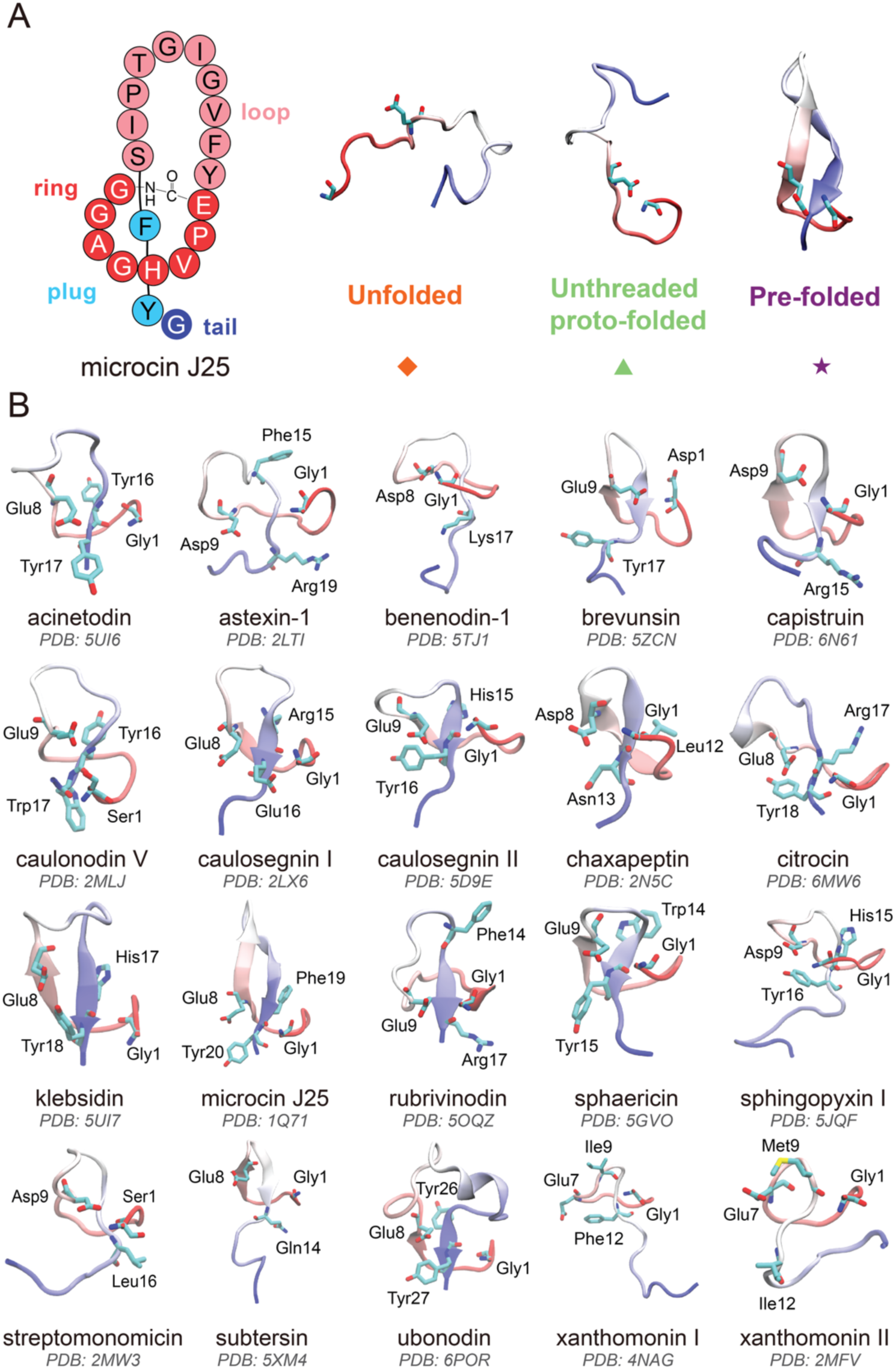
Structural overview and pre-folded conformations of lasso peptides. (A) Structural overview of lasso peptide microcin J25, and schematic representations of the three distinct folding conformations: unfolded (orange diamond), unthreaded proto-folded (green triangle), and pre-folded (purple star). These symbols are used in numerous additional figures. (B) Pre-folded structures of the 20 studied lasso peptides, shown as cartoon representations. The N-terminal region is colored red while the C-terminal region is colored blue. Plug residues and residues forming the isopeptide bond are shown in stick representations.

Despite these broad biological functions, harnessing the therapeutic potential of lasso peptides through rational design is limited by an incomplete understanding of their intrinsic folding propensities. Lasso peptide biosynthesis begins with a the ribosomal production of a precursor peptide, with genomic analysis revealing conserved N-terminal leader and diverse C-terminal core regions.^1,11^ During processing, the leader region is bound by the RiPP recognition element (RRE), which in turn activates the leader peptidase to reveal a new N-terminus at position one of the core peptide.^3,12^ The linear core peptide is then delivered to the cyclase, which adenylates and pre-folds the structure.^3,13^ Before cyclization, the core peptide can form three distinct conformations: unfolded, unthreaded proto-folded, and pre-folded (**Figure 1A**).^14^ Productive lasso peptide formation requires the core peptide to adopt a pre-folded conformation before macrolactam formation.^13^ Nevertheless, it is unclear how much the linear core peptide’s inherent solution-phase flexibility or ability to adopt a pre-folded conformation contributes to substrate selectivity and lasso peptide processing. Ferguson *et al.* showed that the core region of the microcin J25 precursor peptide (i.e., McjA_core_) modified with azidoacetic acid in position one and propargylglycine in place of the acceptor residue could not form microcin J25 (i.e., the mature lasso peptide) through copper-catalyzed azide-alkyne cycloaddition (CuAAC).^15^ Instead, an unthreaded “branched-cyclic” product was observed. Short molecular dynamics (MD) simulations (95 ns) further supported that McjA_core_ did not spend an appreciable amount of time in a pre-folded state.

Recently, da Hora *et al.* explored the *de novo* folding process of McjA_core_ through ∼7.2 µs unbiased MD simulations and ∼3.9 µs replica exchange MD (REMD) simulations.^14^ The results suggested that *de novo* folding process of microcin J25 is rare (probability ∼0.8%). The pre-folded structure was shown to be metastable before ring formation, but it readily transitions to entropically favored unfolded and unthreaded proto-folded structures.^14^ Additionally, all the obtained pre-folded structures were right-handed, consistent with those observed in nature.^14^ Simulations started with chemical modifications via an auxiliary with a thiol attached to the nitrogen atom of Gly1 and thioester functionalities at position 8 show a marked increase in the stability of pre-folded structures (probability ∼3.4%).^14^ Both studies underscored that *de novo* folding of microcin J25 is thermodynamically unfavorable. However, previous studies have focused on a single lasso peptide, which is likely insufficient to fully capture the mechanism of lasso peptide folding. More than 40 lasso peptides without secondary post-translational modifications have been structurally characterized, and their sequences and native folded structures vary substantially from microcin J25.^3,16^ Therefore, it is imperative to investigate the folding processes of multiple distinct lasso peptides to capture a representative folding landscape and elucidate universal principles of lasso peptide folding.^17^

To accurately characterize the folding landscape across a diverse library of lasso peptides, standard MD simulations face a significant hurdle: the disparity between accessible simulation timescales and the slow biological rates of protein or peptide folding^17^. Classical MD simulations must integrate Newton’s equations of motion with femtosecond timesteps (10^-15^ s), while protein or peptide folding occurs over microseconds to seconds. This requires approximately 10^9^-10^12^ integration steps, making direct simulation of complete folding events computationally prohibitive for most systems.^17,18^ Consequently, although individual MD trajectories can capture local conformational fluctuations, global folding events often exceed the simulation trajectory length. To bridge this gap, the Markov State Model (MSM) provides a robust statistical framework that connects short, independent trajectories to reconstruct global conformational changes, thereby enabling the estimation of long-timescale kinetics and thermodynamics from MD simulations.^19–21^ Fundamentally, MSM relies on counting transitions between discrete states to estimate a transition probability matrix governing the system’s dynamics. This method assumes sufficient local sampling of reversible transitions between connected states to satisfy the principle of detailed balance, which dictates that the probability flux between states must be equal in both directions to estimate global equilibrium properties accurately.^22^ However, for systems characterized by high energetic barriers and rare transition events, such as lasso peptide folding, unbiased simulations alone often fail to sufficiently sample rare states to ensure detailed balance for accurate MSM construction.^23,24^

In this study, we combined extensive unbiased and biased MD simulations to systematically investigate the *de novo* folding mechanisms of 20 distinct lasso peptides lacking secondary post-translational modifications, with experimentally determined three-dimensional structures (**Figure 1B**). Unbiased simulations (∼200 µs per lasso peptide) revealed limited sampling of pre-folded states due to a strong kinetic asymmetry between folding and unfolding of the lasso peptides, leading to incorrect MSM construction and overestimation of unfolded states and folding barriers. To address this, we employed the Transition-based Reweighting Analysis Method (TRAM),^24^ a statistically optimal framework that integrates biased simulations, such as umbrella sampling, to enhance sampling of rarely visited states and accurately estimate thermodynamics and kinetics. TRAM leverages the local-equilibrium approximation in MSM and biased simulations to enforce local equilibrium in interstate transitions. It estimates Multi-Ensemble Markov Models (MEMMs) with full thermodynamic and kinetic information across all simulation ensembles (**Figure 2A**). Previous studies have shown that TRAM-predicted kinetics align more closely with experimental results than those from MSM approaches, making TRAM well-suited to characterizing lasso peptide folding.^23,24^ Using TRAM to estimate MEMMs, we achieved robust thermodynamic and kinetic characterization for each of the 20 lasso peptides. Our results demonstrate a universal uphill folding free-energy profile. The pre-folded conformation is sparsely populated (< 0.8%) and thermodynamically unfavorable for all lasso peptides examined. Further analyses indicated two critical determinants of folding efficiency, loop stability and entropic cost. Loop stability strongly correlated with the probability of pre-folded conformations. Experiments on microcin J25 confirmed a correlation between β-hairpin propensity in the loop region and lasso peptide production, validating that enhanced loop stability facilitates lasso peptide folding. A substantial entropy cost was observed for the folding of all lasso peptides. Confinement simulations demonstrated that mimicking the cyclase pocket’s spatial confinement stabilizes folding by restricting extended conformations and reducing the entropic cost, thereby promoting the pre-folded form. Microcin J25 presented the highest propensity for *de novo* folding (∼0.75%) among all studied lasso peptides, owing to the stable loop and low entropic cost. To further decipher the folding mechanisms, we applied transition path theory (TPT) and a Variational AutoEncoder (VAE)-based Latent-space Path Clustering (LPC) method^25^ to find the distinct folding pathway channels and representative kinetic folding pathway, confirming the common hallmarks of *de novo* lasso peptide folding, such as β-hairpin formation in the loop region and its role in directing folding. Collectively, our integrative framework provides a representative view of the thermodynamic and kinetic principles underlying the *de novo* folding of lasso peptides, thereby establishing guidelines for the rational engineering of designed lasso peptides.

**Figure 2.**
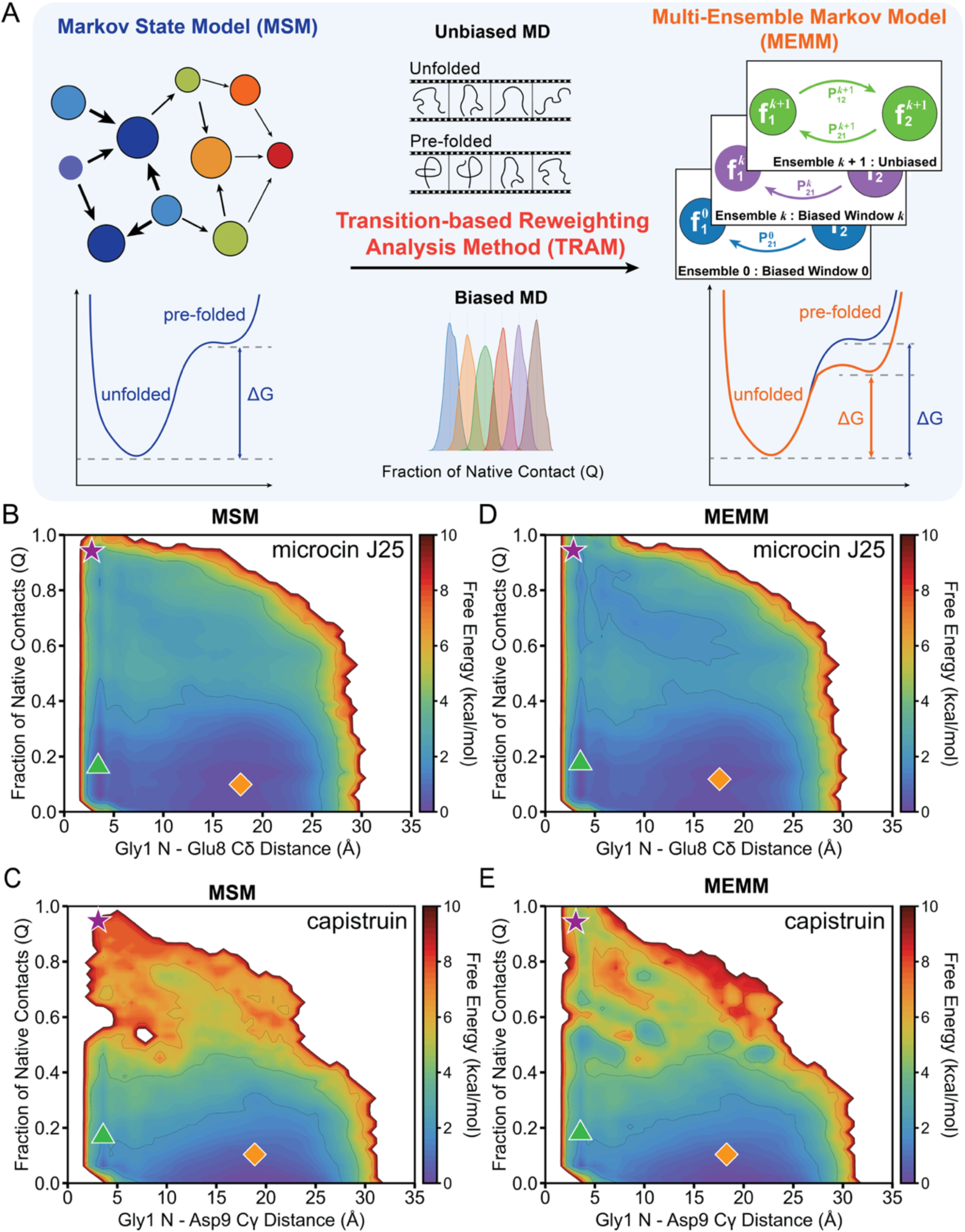
Lasso peptide folding free energy landscapes resolved by MSM and MEMM. (A) Schematic comparison of MSM and MEMM. (B–C) Two-dimensional free energy landscapes obtained from MSM for microcin J25 (B) and capistruin (C). (D–E) Two-dimensional free energy landscapes obtained from MEMM for microcin J25 (D) and capistruin (E). Representative conformations are marked on the free energy landscapes using the same symbols and color scheme as in Figure 1A. **Universal Uphill Thermodynamics Underlying Rare *De novo* Lasso Peptide Folding**

## Results

### Multi-Ensemble Markov Model Resolves Lasso Peptide Folding Landscapes

Through extensive MD simulations, we first systematically assessed the ability of each lasso peptide to adopt and maintain its threaded conformation in the absence of the lasso cyclase enzyme (pH-neutral water, 300 K), mapping their folding landscapes to elucidate the *de novo* folding mechanism. To quantitatively characterize the conformational landscape of lasso peptide folding, we employed two complementary metrics: the fraction of native contacts (Q)^26^ and the ring-closure distance. Q quantifies the overall similarity to the pre-folded structure by measuring the proportion of residue-residue interactions relative to those in the native structure.^26^ Ranging from 0 (unfolded) to 1 (pre-folded), Q has proven effective as a folding coordinate for atomistic simulations of proteins and peptides. ^26,27^ In this study, we established a threshold of Q ≥ 0.8 to indicate the pre-folded structure across all lasso peptides. Barrett *et al.* performed MD simulations of the adenylated, pre-folded fusilassin intermediate bound within the cyclase FusC (WP_104612995.1) catalytic pocket. The highest probability density peak around Q = 0.8 indicates a strong likelihood that adenylated FusA retains the pre-folded shape, supporting Q ≥ 0.8 as a reasonable threshold for defining a pre-folded conformation.^27^ However, Q alone is insufficient to distinguish between the unthreaded proto-folded and unfolded structures, since Q would be low in both cases. Therefore, ring-closure distance, i.e., the distance between the N-terminus and the side chain carboxylate carbon of acceptor residue (Cδ of Glu or Cγ of Asp), is applied as a complementary metric and direct indicator of ring closure. The ring-closure distances of the pre-folded conformations for each lasso peptide were calculated and summarized in **Table S1**. Based on prior observations for microcin J25, we applied a ring-closure distance threshold of 6 Å, beyond which the macrolactam ring is insufficiently constrained to enforce the threaded conformation.^14^ da Hora *et al.* developed a tool to distinguish pre-folded from unthreaded proto-folded conformations for lasso peptides.^14^ We applied this tool to microcin J25 and found that >97% of conformations with Q *≥* 0.8 and a ring-closure distance *≤* 6 Å were classified as pre-folded, suggesting that joint Q and ring-closure distance are effective metrics for identifying pre-folded lasso structures.

For each lasso peptide, we initially performed ∼200 µs of unbiased MD simulations starting from both pre-folded and unfolded structures. We constructed MSMs for each lasso peptide (**Figure 2A**) and further projected the data onto two-dimensional free energy landscapes (**Figure 2B-C, Figure S4**) using the two previously described metrics. For microcin J25, we observed a connected free energy landscape encompassing all three conformational states. However, capistruin, like most of the lasso peptides studied, exhibited a disconnected free energy landscape with rare pre-folded states and a much higher folding free energy barrier. This discrepancy arises from kinetic asymmetry, in which the unfolding process proceeds much more rapidly than the corresponding folding process. This leads unbiased simulations to predominantly sample unfolded states and undersample pre-folded states. The rapid unfolding process of microcin J25 was previously observed in unbiased simulations starting from pre-folded structures.^14^ Within 500 ns, microcin J25 started unfolding and fluctuated between the pre-folded and unthreaded proto-folded states.^14^ Similarly, we observed this unfolding process in trajectories initiated from the pre-folded structure, but no folding in trajectories initiated from the extended structure for microcin J25 (**Figure S5**). For capistruin, this unfolding process occurred even more rapidly within 10 ns. This kinetic asymmetry poses a fundamental challenge to the reliable construction of MSMs. Detailed balance, the guiding principle of MSM, dictates that the probability flux between connected states must be equal in both directions at equilibrium, implying that accurate free energy can be computed only if the simulation captures these reciprocal transitions.^22^ However, the sampling deficiency in lasso peptide unbiased simulations leads MSM to overestimate the thermodynamic stability of unfolded states relative to pre-folded states, resulting in artificially high free-energy folding barriers and ultimately inaccurate thermodynamic and kinetic estimates.^23,24^

To overcome these limitations, we implemented TRAM,^23^ a statistically optimal framework that leverages biased simulations to populate rarely visited states and estimates thermodynamics and kinetics using reweighting estimators (**Figure 2A**). Biased simulations, implemented here via umbrella sampling, facilitated exploration of high free energy pre-folded and transition states that are rarely visited in unbiased trajectories, effectively capturing critical folding transitions across distinct ensembles.^23^ TRAM subsequently integrated biased and unbiased data, enforcing local equilibrium within each ensemble to estimate a MEMM that reconstructs the global equilibrium thermodynamics and kinetics.^23^ Umbrella sampling was performed along *Q* as the reaction coordinate for each lasso peptide. All frames from unbiased simulations were distributed into 50 equally spaced umbrella sampling windows spanning *Q* from 0 to 1. We further combined the unbiased and biased data to construct MEMMs for each lasso peptide. For microcin J25, the MEMM-weighted landscape (**Figure 2D**) closely resembles the MSM-weighted landscape, yielding a well-connected free energy surface spanning all three states. Compared with other lasso peptides studied, microcin J25 has a more stable pre-folded structure; therefore, it allows unbiased simulations alone to adequately sample the conformational space and satisfy MSM detailed balance. This agreement demonstrates that when sampling is sufficient, both MSM and MEMM yield consistent thermodynamic predictions. In contrast, capistruin exemplifies the advantages of MEMM. For a more flexible lasso peptide, unbiased sampling is insufficient for MSM construction, leading to disconnected states and unreliable transition statistics. MEMM overcomes these limitations and produces a properly connected free energy landscape with all three possible conformational states observed (**Figure 2E**), enabling more reliable estimation of folding free energy.

Comparison between MEMM-weighted free energy landscapes (**Figure 3A**) and the corresponding MSM-weighted landscapes (**Figure S4**) reveals that MEMM consistently yields a more robust thermodynamic characterization, particularly for lasso peptides in which conformational sampling of the pre-folded state is challenging in unbiased simulations. Notably, all studied lasso peptides exhibit a universal uphill folding free energy profile. In the two-dimensional free energy landscape, the unfolded ensemble resides at the global minimum, and folding proceeds via a series of progressively higher-energy intermediates toward the pre-folded states. This directly reflects the kinetic asymmetry inherent to lasso peptide folding dynamics: unfolding proceeds as a thermodynamically favorable, kinetically facile process, whereas folding requires overcoming an energetically uphill barrier, resulting in intrinsically slower kinetics. We further elucidated the ability of each lasso peptide to adopt and maintain the pre-folded conformation, as well as its folding free energy. The probability of the pre-folded conformation for all 20 lasso peptides was quantified by the sum of MEMM weights for pre-folded frames with the threshold of Q *≥* 0.8 and ring-closure distance *≤* 6 Å (**Figure 3B**). Microcin J25 emerged as a notable outlier, with the highest probability (∼0.75%) for pre-folded conformations. This result is consistent with the previous study which reported that the probability of reaching the pre-folded state for microcin J25 is ∼0.8%.^14^ Besides, rubrivinodin showed a slightly lower probability of ∼0.63%; however, the other 18 lasso peptides exhibited probabilities of pre-folded states below 0.2%. This demonstrates the inherent thermodynamic challenge of stabilizing the pre-folded conformation purely through intramolecular interactions and without the scaffolding provided by the lasso peptide cyclase during biosynthesis. The thermodynamic basis for these differences becomes apparent on examining the folding free energy (ΔG_f_, **Figure 3C**). Consistent with their higher pre-folded populations, microcin J25 and rubrivinodin display comparatively low folding free energies (ΔG_f_ < 3 kcal·mol⁻¹), in contrast to the substantially higher ΔG_f_ observed for all other peptides. ΔG_f_ demonstrates a strong linear correlation with the probability of the pre-folded conformation with a Pearson correlation coefficient *r* of -0.81 and a p-value of 1.6×10^-5^. This outstanding negative correlation arises directly from the universal uphill folding landscapes observed for all lasso peptides. Because the unfolded ensemble occupies the global minimum, the probability of forming the pre-folded state depends strongly on the height of the pre-folded basin along the uphill free energy surface. Lasso peptides with smaller ΔG_f_ therefore correspond to shallower uphill landscapes and exhibit higher, though still rare, *de novo* pre-folded populations.

**Figure 3.**
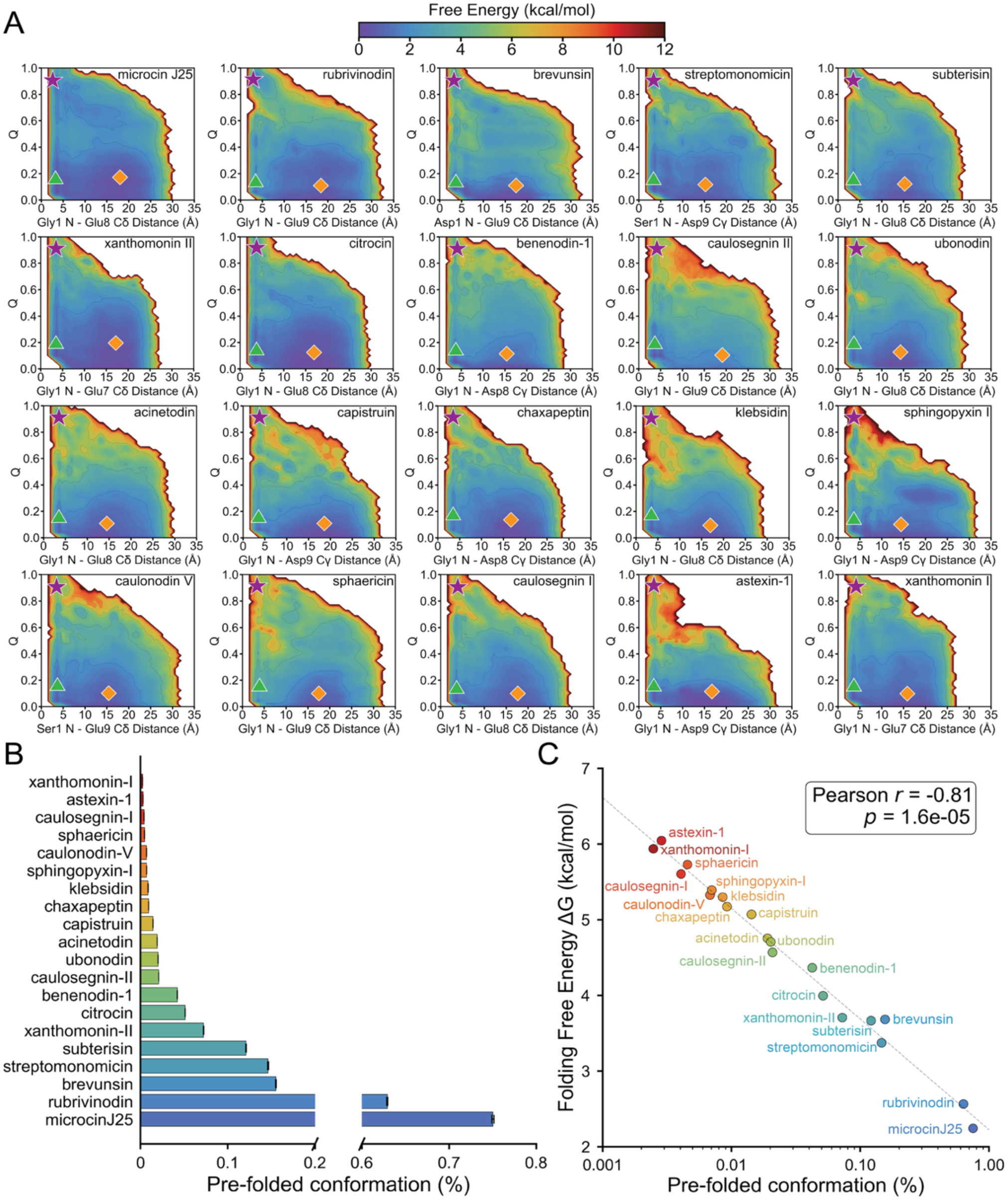
Universal uphill folding landscapes underlying the rare *de novo* folding of lasso peptides. (A) Two-dimensional free energy landscapes obtained from MEMM for all 20 lasso peptides analyzed in this study. Representative conformations are marked on the free energy landscapes using the same symbols and color scheme as in Figure 1A. Panels are arranged by decreasing probability of the pre-folded conformation, as quantified in panel B. (B) Probability of the pre-folded conformation for each lasso peptide. Error bars denote the standard deviation estimated from bootstrapping. Colors encode the normalized probability, ranging from low (red) to high (blue). (C) Correlations between folding free energy ΔG_f_ and the probability of pre-folded conformation. Each lasso peptide label is colored consistently with panel B.

### Loop Stability Facilitates *De Novo* Lasso Peptide Folding

The loop region plays a crucial role in the efficiency of lasso peptide formation. The sequence composition, length, and secondary structure of the loop region dictate the peptide’s capacity to establish stable intramolecular contacts to maintain threadedness. To quantitatively assess loop dynamics and stability, we calculated the relaxation time of the loop fraction of native contact, Q_loop_. The time-dependent expectation value of the Q_loop_ was computed from the transition probability matrix of MEMM, and the relaxation time was defined as the point at which the Q_loop_ reaches a stable plateau (changes less than 0.1%). Lasso peptides with extended Q_loop_ relaxation times, indicating greater loop stability, exhibited higher probabilities of the pre-folded conformation (Pearson *r* = 0.66, **Figure 4A**). This is also supported by a recent study that demonstrated limited tolerance for sequence variation in the fusilassin loop region, underscoring its critical role in interactions with the lasso cyclase.^27^

**Figure 4.**
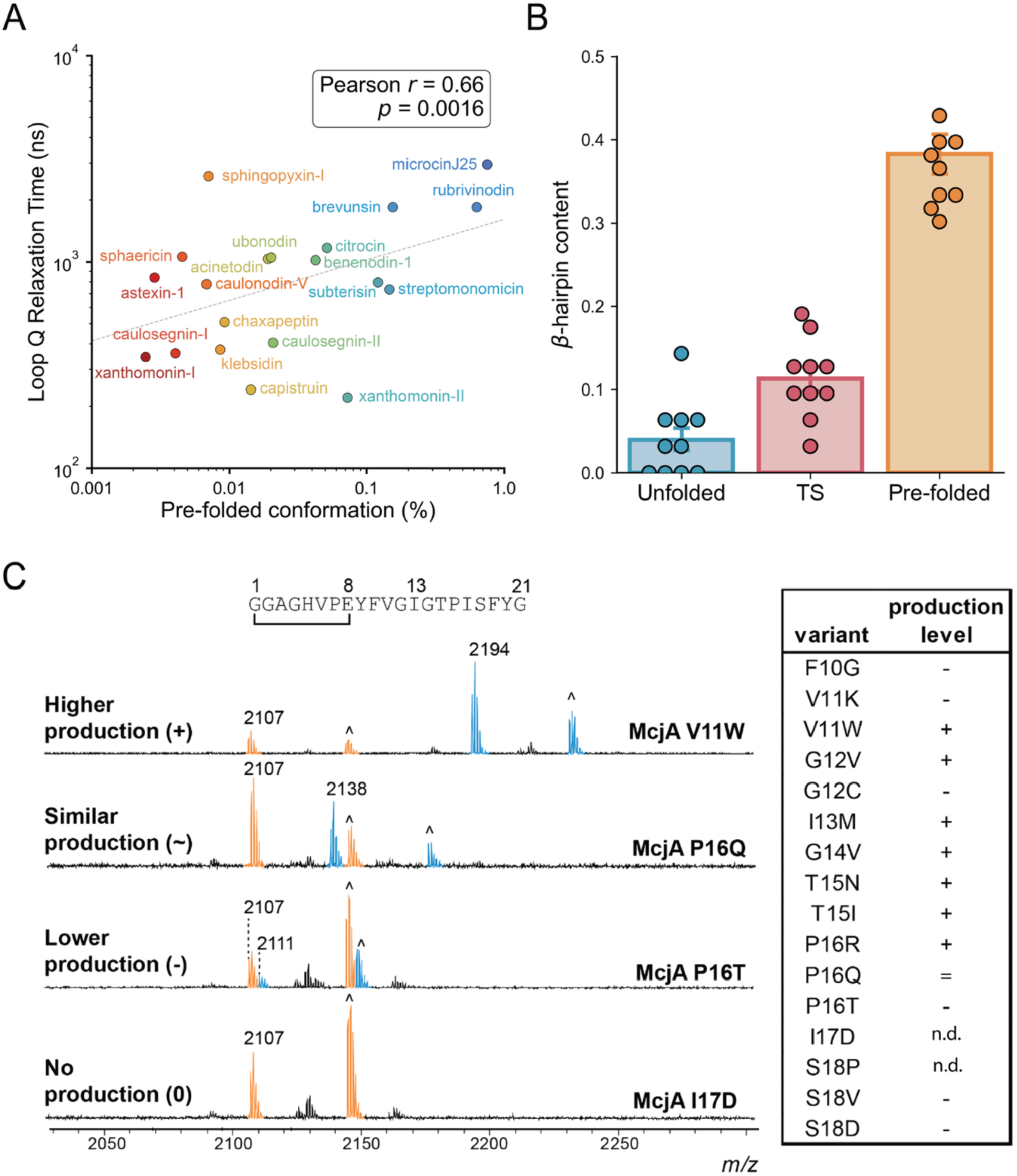
Computational and experimental analysis of loop stability in *de novo* lasso peptide folding. (A) Correlations between Q_loop_ relaxation time and the probability of pre-folded conformation. Each peptide label is colored consistently with Figure 3B. (B) The β-hairpin content of the unfolded states, the transition states (TS), and the pre-folded states in microcin J25 simulations. (C) Representative MALDI-TOF-MS spectra for variants with (+) high production relative to wild-type, (=) similar production, (−) lower production, and (n.d.) no detectable. The wild-type [M+H]^+^ and [M+K]^+^ are highlighted in orange while the variant [M+H]^+^ and [M+K]^+^ are blue. The number indicates the [M+H]^+^ mass for the McjA wild-type and variant peaks. The ^ indicates the [M+K]^+^ peaks. The table shows the variants and their production level.

Loop stability in lasso peptides is closely associated with the emergence of secondary structure, particularly β-hairpin. For instance, microcin J25, with the highest probability of maintaining the pre-folded state, possesses a relatively long loop that forms the most stable region by adopting a well-defined β-hairpin structure. Klebsidin shares with microcin J25 the same 8-residue ring size, identical ring-closure residues (Gly and Glu), a comparable loop length (8 and 10 residues for klebsidin and microcin J25, respectively), and a one-residue tail (excluding plug). Despite these similarities, klebsidin displays reduced β-hairpin content in its loop region, resulting in weaker loop stabilization. This structural difference is manifested as a shorter Q_loop_ relaxation time and a reduced propensity to populate pre-folded states for klebsidin. In addition, lasso peptides such as xanthomonin-I and chaxapeptin possess extremely short loops of only one to three residues, which are inherently incapable of supporting secondary structure formation. This absence of stabilizing interactions results in minimal loop persistence, as reflected in relatively short Q_loop_ relaxation times and correspondingly low probabilities of adopting pre-folded conformations.

To further clarify the importance of β-hairpin loop stability in microcin J25 folding, we conducted a quantitative comparison of the β-hairpin content in different states. In addition to the unfolded and pre-folded states that are distinct in the free energy landscape, identifying the transition states between them is crucial for understanding conformational changes during folding. Transition states typically comprise a set of structures located at a region of the free energy map that separates the reaction and product free energy basins. To identify these states, we employed a deep learning network named TS-DAR (Transition State identification via Dispersion and vAriational principle Regularized neural networks), which is designed to detect out-of-distribution (OOD) data and capture the transition states.^28^ By jointly optimizing the VAMP-2 score and dispersion loss, TS-DAR learned a regularized hyperspherical latent representation for each conformation along the MD trajectories. Within this latent space, both the unfolded and pre-folded metastable states were compactly arranged and uniformly separated. The transition states between these two metastable states will have equal angular distances to the metastable state centers. The β-hairpin content observed in the transition states lies between that of the unfolded and pre-folded states, indicating the initiation of β-hairpin formation during the transition states (**Figure 4B**).

Based on this mechanistic insight, we hypothesized that variants that disrupt the β-hairpin content of the core peptide would decrease or prevent lasso peptide folding. We predicted the β-hairpin content of all possible single-site variants of microcin J25 using S4PRED.^29^ We selected 16 variants, conducted 1 µs MD simulations for all variants, and calculated β-hairpin content using the Define Secondary Structure of Proteins (DSSP) algorithm implemented in MDTraj.^30^ The β-hairpin content calculated from the MD simulations is summarized in **Table S5**. To experimentally validate these findings, we selected three variants predicted to increase β-hairpin content and three additional variants predicted to decrease it. Repeated attempts using circular dichroism (CD) spectroscopy were unsuccessful, likely due to the general challenge of observing less intense features such as β-hairpin by CD, which is further exacerbated by the low percentage of β-hairpin containing peptides in the population.^31,32^ Complementary NMR analysis was also attempted to identify long-range Nuclear Overhauser Effect (NOE) cross-peaks indicative of lasso folding, but did not yield conclusive results.

To answer whether predicted β-hairpin content correlated with lasso variant production, we used cell-free biosynthesis (CFB) to rapidly evaluate lasso peptide formation in vitro. First, a CFB system capable of robustly producing wild-type microcin J25 was optimized. This system relied on *E. coli* lysate containing microcin J25 biosynthetic enzymes (leader peptidase McjB, lasso cyclase McjC, and ABC transporter McjD). The enzyme-containing lysate was then used to express the McjA precursor peptide fused to maltose-binding protein (MBP) (**Figure S7, S8**). After an overnight reaction, Matrix-Assisted Laser Desorption Ionization Time of Flight Mass Spectrometry (MALDI-TOF-MS) was used to determine whether the biosynthetic enzymes could cyclize the variants. The production observed in CFB was consistent with previously reported heterologous production data.^33^ The only exception was McjA S18V. We observed production in CFB but not in heterologous expression, suggesting that the polar S18 side chain is important for export but not cyclization.^34^

To test whether McjA variants predicted to enhance or disrupt β-hairpin content affected the production of the lasso variant, we used a competition experiment. Here, equal concentrations of the DNA encoding the wild-type precursor and the variant precursor were added to the CFB reaction. MALDI-TOF-MS peak heights for the wild-type microcin J25 and variant peptides were then compared and qualitatively ranked (**Figures 4C, S9**). Efforts to quantify the variant’s production relative to the wild-type product using Ultra High Performance Liquid Chromatography (UHPLC) were unsuccessful due to peak overlap. Of the sixteen variants, the production of ten correlated well with the β-hairpin content predicted by the MD simulations (∼63%). The production levels for the variants with the most extreme difference in predicted β-hairpin content from the wild type (decrease: S18P and V11K, increase: G14V, T15I, and V11W) aligned well with the predicted β-hairpin content values. However, the correlation was less clear for variants with β-hairpin content prediction values closer to the wild-type value. This suggests that there are more variables to lasso peptide processing than just the core peptide’s propensity to adopt a β-hairpin or lasso-like conformation. Indeed, these cell-free competition experiments include the cyclase and β-hairpin formation may be less critical in the enzyme’s active site than in solution.

### Entropic Cost Limits *De Novo* Lasso Peptide Folding

During lasso peptide biosynthesis, a flexible linear peptide is folded into a conformationally restricted, lower-entropy structure. To quantitatively evaluate the contribution of entropy to folding energetics, we employed the PARENT program to calculate conformational entropy for both pre-folded and unfolded ensembles.^35^ Across 20 lasso peptides, folding entropic cost displayed a moderate correlation (Pearson *r* = -0.55) with the probability of pre-folded conformation (**Figure 5A**). To consider sequence-length influence, we normalized the entropic cost by peptide length and reported the average folding entropic cost per residue (**Figure S10**). A slightly better correlation (Pearson *r* = -0.58) between average entropic cost per residue and pre-folded conformation probability was observed.

**Figure 5.**
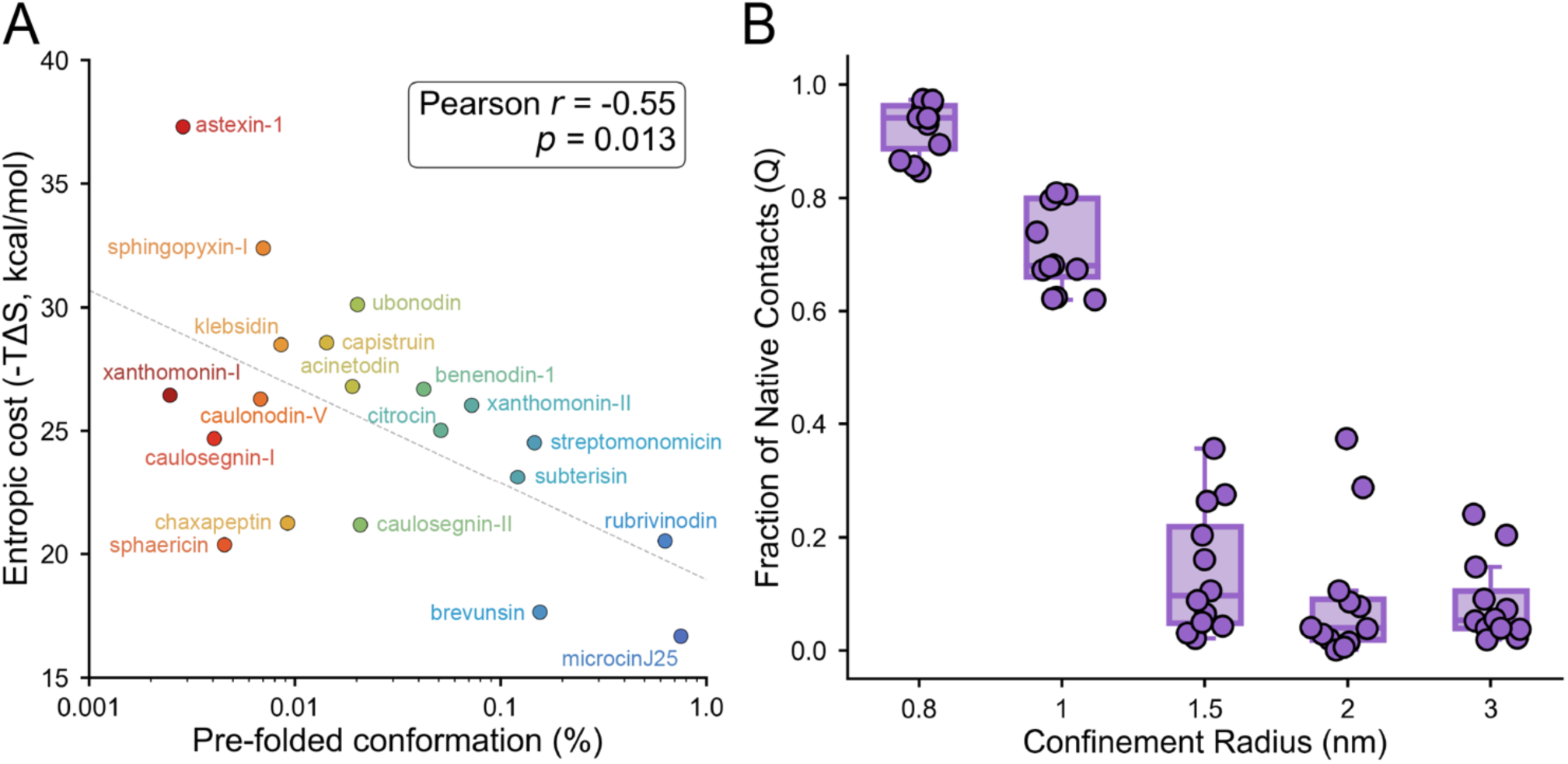
Spatial confinement reduces the entropic cost of *de novo* lasso peptide folding. (A) Correlations between entropy cost and the probability of pre-folded conformation. Each peptide label is colored consistently with Figure 3B. (B) Average Fraction of Native Contacts throughout MD simulations of capistruin under sphere confinement with varying radius.

In general, the transition from a flexible, unfolded structure to a compact, pre-folded conformation requires a substantial reduction in conformational freedom, thereby imposing a large entropic penalty. According to the Gibbs equation, Δ*G* = Δ*H* − *T*Δ*S*, the substantial entropic penalty (−*T*Δ*S*) of lasso peptide folding can be partially offset by stabilizing interactions. Such stabilization can arise from enthalpic interactions, including intramolecular hydrogen bonds and electrostatic interactions, as well as hydrophobic packing, which provides favorable solvent entropy through desolvation. Consequently, variations in loop secondary structure determine how effectively stabilizing interactions can partially compensate for the entropic penalty of folding, leading to pronounced differences in folding efficiency among lasso peptides. Lasso peptides similar to microcin J25 with well-structured loop regions rich in β-hairpins fold efficiently by maximizing enthalpic gains while minimizing entropic penalties. In contrast, peptides with long but unstructured loops lack stabilizing secondary interactions; for example, astexin-1 and klebsidin incur high entropic costs with little compensatory enthalpy, resulting in lower folding efficiency. However, these differences reflect only the *de novo* folding propensities of lasso peptides. Even peptides with the highest intrinsic propensity to adopt pre-folded conformations do not fold efficiently on their own and require a cyclase for productive lasso peptide formation. Given the dominant entropic penalty associated with adopting the threaded conformation, we hypothesized that the cyclase enzyme facilitates folding in part by spatial confinement that restricts conformational freedom and effectively reduces the entropic cost of forming and maintaining the pre-folded state. To validate our hypothesis, we conducted a confinement simulation for capistruin, a representative lasso peptide with a high folding free energy and entropic cost. Specifically, we constructed spherical confinement potentials of varying radii to simulate the restricted environment of the cyclase enzyme pocket. The most intense confinement with the smallest radius (r = 0.8 nm) maintained a sustained Q around 1.0 for capistruin throughout the simulations (**Figure 5B**, **S11**), indicating its capacity to consistently preserve the pre-folded conformation. However, when the confinement radius was increased to 1.0 nm, capistruin exhibited a gradual loss of Q in the initial stages of the simulation, though the decay occurred at a relatively slow rate, suggesting a partial retention of the pre-folded structure. However, a further increase in the sphere radius resulted in a rapid, pronounced loss of the pre-folded structure within 10 ns of the simulation. These results show that spatial confinement of the cyclase pocket effectively reduces the entropic cost, thereby stabilizing lasso peptide folding intermediates and preserving the pre-folded conformation.

Taken together, lasso peptide *de novo* folding is dominated by unfavorable entropic penalties with comparatively limited enthalpic stabilization, rendering spontaneous folding in solution intrinsically unlikely. Efficient folding, therefore, requires minimizing entropic costs through loop pre-organization and secondary structure formation, while simultaneously maximizing stabilizing enthalpic interactions. In addition, other factors not examined in this study, such as N- or C-terminal flexibility and plug residue identity, may also modulate folding efficiency. For example, Gly is strongly overrepresented at position 1 of lasso peptide cores because its small size facilitates backbone turning by minimizing steric hindrance and lowering the entropic cost of N-terminal fixation^27,33,36^.

During lasso peptide processing, cyclase enzymes provide a critical chaperone environment that helps overcome these intrinsic thermodynamic challenges. By confining the peptide substrate within a structured binding pocket, the cyclase reduces the entropic penalty of folding by limiting conformational freedom and stabilizing intermediates. At the same time, the cyclase establishes specific enzyme–substrate interactions that contribute to favorable enthalpic stabilization and facilitate productive threading. While cyclases are ATP-dependent enzymes and adenylation is required to generate the electrophilic intermediate for ring closure, it remains unclear to what extent the energy released during ATP hydrolysis directly offsets the entropic penalties. Thus, cyclase-mediated chaperoning likely plays a multifaceted role in enabling the successful formation of lasso peptides that would otherwise be unable to fold spontaneously in solution.

### *De novo* Lasso Peptides Folding Pathways

Due to the diversity of 20 lasso peptides studied here, it is reasonable to expect that each peptide may adopt distinct folding pathways. Mapping these pathways enables us to elucidate details of lasso folding, identify kinetic bottlenecks, and ultimately derive insights into lasso peptide engineering. By constructing MEMM, the entire conformational space is discretized into multiple microstates (**Figure 6A**). We applied Transition Path Theory^37^ (TPT) to identify an ensemble of 10,000 transition pathways and obtained the most representative folding pathway with the highest flux on the two-dimensional free energy landscape (**Figure 6A**). However, it is challenging to comprehend the molecular mechanism of folding from thousands of individual kinetic pathways, each representing a small fraction of the total folding flux. Moreover, at a coarser level, different pathways may share substantial structural similarity; such pathways can be defined as the same higher-level “route” or “transition tube”.^38^ We applied a Variational Autoencoder (VAE) based algorithm, known as Latent space Path Clustering^25^ (LPC), to effectively discern diverse folding routes. The resulting two-dimensional latent space representation (**Figure 6B**) provided a compact visualization of the folding pathway diversity of microcin J25. Each point in the latent space represents a single pathway, and the size is determined by its corresponding normalized flux value. Subsequently, K-means clustering was applied to group pathways in the VAE latent space, and the optimal number of clusters was determined through silhouette analysis.^39^ Two distinct and separated metastable path channels were identified: one channel comprising 74% of the total folding flux and another channel accounting for 26% of the total flux. We next presented structural details of the most representative pathway in channel 1, which carries the maximum folding flux, and calculated the folding transition time, the Mean First-Passage Time (MFPT, **Figure 6C**). For microcin J25, folding begins with *β-hairpin* formation at approximately 9.91 µs, with partial completion after an additional 28.1 µs. This β-hairpin formation stabilizes the loop region, which is a prerequisite for threading and ring closure. During this process, the loop and tail have largely formed, but the N-terminal region remains mobile. After 13.6 µs, N-terminal turning begins, and the β-hairpin content continues to increase, fully forming after another 6.23 µs. Finally, the N-terminal segment crosses the C-terminal loop and tail to achieve a pre-folded conformation within an additional 3.03 µs. The β-hairpin formation time for microcin J25 occurred on a similar timescale to the simulated folding times of the WW domain (21 μs), NTL9 (29 μs), and Protein G (65 μs), each of which forms β-sheet structure during folding^17^. This folding pathway demonstrates the dual roles of β-hairpins in lasso formation: providing structural stability while constraining the loop geometry to enable further threading and ring closure.

**Figure 6.**
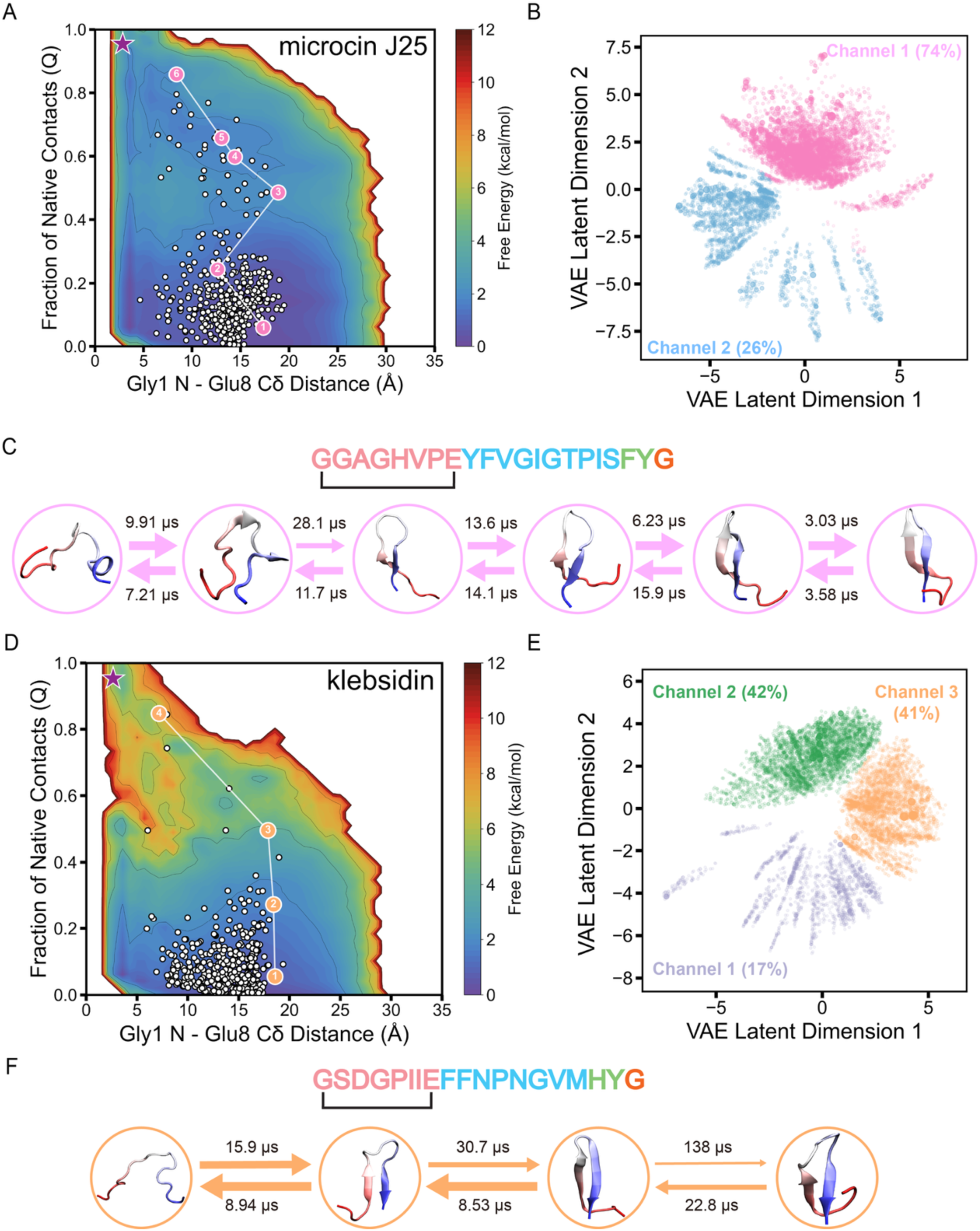
Folding pathways and channels of microcin J25 and klebsidin. (A) Distribution of microstate centers on the MEMM-weighted free energy landscape of microcin J25 and the most representative folding pathway highlighted with the color of its corresponding pathway channel. (B) Kinetic pathways and pathway channels of microcin J25 in the latent space captured by the VAE-based LPC algorithm. Each point in the latent space represents a single pathway, and its size was determined by the corresponding normalized flux value. Each pathway channel was color-labeled with its relative percentage of the total flux. (C) Structural details of the most representative folding pathway with MFPT calculated between every two adjacent states for microcin J25. The same analyses are presented for klebsidin (D–F) for comparison.

Compared with microcin J25, the MEMM-weighted free energy landscape for klebsidin (**Figure 6D**) exhibits unfavorable thermodynamics and a much lower folding probability. Applying LPC to 10,000 TPT pathways for klebsidin folding, we identified three distinct folding pathway channels (**Figure 6E**). The most representative pathway in channel 3 (**Figure 6F**) revealed that klebsidin initiates β-hairpin formation in the loop region at 15.9 µs, which completes after an additional 30.7 µs, slightly slower than microcin J25. However, the most significant difference lies in the final threading step: klebsidin requires 138 µs to form the pre-folded structure. During this prolonged interval, the N-terminus initiates turning, but this process is destabilized by partial loss of secondary structure in the loop region. Specifically, the β-hairpin undergoes local unfolding, resulting in a more flexible loop. This loss of pre-organized secondary structure constitutes a key mechanistic bottleneck in the folding of klebsidin. Thus, microcin J25 represents a highly optimized case, with fast and stable β-hairpin formation that promotes efficient threading. In contrast, klebsidin exemplifies how destabilized secondary structure can prolong folding times and reduce the probability of pre-folding.

We extended this folding pathway analysis to all remaining lasso peptides (**Figures S13–S30**). A recurring theme emerged: for 12 of the 20 studied lasso peptides, β-hairpin formation in the loop region consistently serves as the initiating event in folding, and its successful formation contributes to lasso loop stability and the eventual formation of pre-folded structure. Despite sharing highly similar sequences and structures, some lasso peptides demonstrate pronounced divergence in their folding kinetics. Caulosegnin I requires substantially longer (1.57 ms, **Figure S19**) to form the loop β-hairpin than caulosegnin II (67.4 µs, **Figure S20**). Although both peptides contain a single proline within the loop, caulosegnin II folds more rapidly because its loop region contains a Pro-Gly motif immediately upstream of the β-hairpin initiation site at His15, which stabilizes a tight β-turn, thereby accelerating downstream β-hairpin nucleation. In addition, the additional prolines in the ring and tail regions, together with the intrinsic β-carbon methyl groups of threonine residues, further rigidify the surrounding scaffold in caulosegnin II, thereby reducing conformational flexibility. In addition, xanthomonins I and II are distinguished from the rest of the dataset by having the smallest ring regions (7 residues), identical ring sequences, short loops (5 residues), and long tails (8 residues). These lasso peptides, however, exhibit distinct kinetics for forming the final pre-folded structure: xanthomonin I folds 3-fold slower (306 µs, **Figure S29**) than xanthomonin II (102 µs, **Figure S30**). Because neither lasso peptide forms a stabilizing loop β-hairpin, the final transition is instead governed by the conformational behavior of the tail. The presence of two C-terminal Glu residues increases solvation and electrostatic repulsion, therefore xanthomonin I displayed a more extended tail along the folding pathway (**Figure S29**). This tail extension increases conformational entropy, reducing the probability of threaded conformations and slowing the folding process.

## Discussion

In this study, we present a representative computational and experimental investigation of the *de novo* folding mechanisms of lasso peptides lacking secondary post-translational modifications. By integrating extensive unbiased and biased molecular dynamics simulations with Multi-Ensemble Markov models, we elucidated shared thermodynamic and kinetic principles governing lasso peptide folding. We observed a universal uphill folding free energy landscape, in which unfolded conformations are thermodynamically favored, and pre-folded conformations are intrinsically rare. Across all 20 lasso peptides examined here, the probability of pre-folded conformations remains below 0.8%, underscoring that spontaneous lasso peptide folding is disfavored in solution and generally requires enzymatic assistance for efficient formation.

Loop stability plays a central role in shaping folding efficiency. Lasso peptides with pre-organized loops, often stabilized by β-hairpin structures, exhibit longer loop relaxation times and higher probabilities of pre-folded states. Among all the lasso peptides studied, microcin J25, which displays exceptional loop stability, shows the highest folding probability. Consistent with these observations, experiments revealed a strong correlation between β-hairpin propensity in the loop region and successful production of microcin J25. Variants predicted to enhance β-hairpin formation generally exhibited higher production yields, supporting the critical role of loop stability in efficient folding. Comparative kinetic pathway analysis between microcin J25 and klebsidin further demonstrates that destabilization of loop β-hairpin structures constitutes a major kinetic bottleneck that hinders productive folding.

In addition, folding is strongly limited by entropic cost. Formation of the threaded, pre-folded structure requires a substantial reduction in conformational freedom, imposing a dominant entropic penalty that disfavors the de novo folding. Confinement simulations demonstrated that mimicking the spatial restriction of the cyclase pocket markedly stabilizes the threaded conformation by limiting accessible conformations and reducing the folding entropic cost. These results support a model in which cyclases facilitate lasso peptide formation not only by catalyzing macrolactam formation but also by acting as folding chaperones. By confining the peptide substrate within a structured binding pocket, the cyclase simultaneously reduces the entropic penalty of folding. It provides favorable enzyme–substrate interactions that stabilize intermediates and promote productive threading.

Collectively, our results establish a mechanistic framework for lasso peptide folding. These insights provide guiding principles for the rational design and engineering of lasso peptides with improved folding efficiency and stability, and for expanding the functional repertoire of this promising class of natural products.

## Methods

### Fraction of native contacts (Q)

Q is defined as the fraction of heavy atoms in the native structure that are in close spatial proximity at some instant.^26^ The detailed definition of Q is as follows:

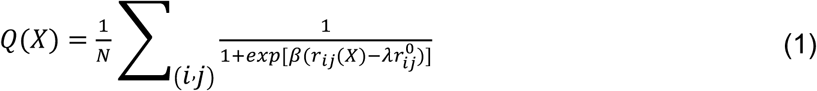

where N is the total number of pairs of heavy atoms (*i*, *j*). All pairs of heavy atoms *i* and *j* belonging to residues *θ_i_* and *θ_j_* are in contact with a residue separation |*θ_i_* − *θ_j_*| > 3, and the distance between *i* and *j* is less than 4.5 Å. *r*_ij_(*X*) is the distance between *i* and *j* in conformation X, *r_ij_*^0^ is the distance between *i* and *j* in the native state. *β* is a smoothing parameter, and the factor *λ* accounts for fluctuation when the contact is formed. *β* = 5 Å ^−1^ and *λ* = 1.8 are set for all-atom models. In this work, Q was calculated using MDTraj.^40^

### Unbiased Molecular Dynamics (MD) simulations

Unbiased MD simulations were performed starting from the pre-folded and unfolded structure of all 20 lasso peptides. Core sequence, ring closure distance, Protein Data Bank (PDB) ID, and structure determination method are provided in **Table S1**. All 20 lasso peptides were solvated in a TIP3P water box and neutralized with Na^+^ and Cl^−^ ions. Atomic interactions were characterized using the Amber ff14SB force field implemented in the Amber18 software.^41,42^ All systems were first minimized for 5000 steps using the steepest descent method, and further minimized for 45000 steps using the conjugate gradient method. The minimized systems were then heated from 0 K to 300 K in the NVT ensemble for 3 ns using a Langevin thermostat with a collision frequency of 2 ps^−1^.^43^ The heated systems were pressurized for 2 ns under an NPT ensemble with a constant pressure of 1 bar using a Monte Carlo barostat.^44^ The peptide backbone atoms were harmonically restrained with a force constant of 5 kcal·mol^−1^·Å^−2^ during heating and pressurization steps. Then the restraints were removed, and the systems were further equilibrated for 50 ns in the NPT ensemble (300 K and 1 bar). The SHAKE algorithm was applied to constrain hydrogen-bond lengths.^45^ A cutoff distance of 10 Å was used for non-bonded force calculation. The Particle Mesh Ewald method was used to compute contributions of long-range electrostatic interactions.^46^ Periodic boundary conditions were maintained throughout the simulation. Hydrogen mass repartitioning was used to increase the simulation time step to 4 fs by redistributing mass between hydrogen atoms and heavy atoms of the peptide.^47^ The production runs were performed in the NPT ensemble using OpenMM^47^ software on the distributed computing platform folding@home.^48^

### Markov State Model (MSM)

MSM provides a framework for analyzing MD trajectories by discretizing the conformational space into kinetically distinct states and modeling transitions between them as a Markov process. The fundamental principle underlying MSM is the assumption of local equilibrium, where transitions between states satisfy the detailed balance:

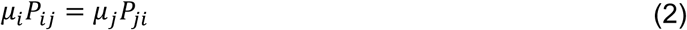

where *μ_i_* is the stationary density of the Markovian state *i* and *P_ij_* represents the probability of transition from the *i* to *j* state at the chosen lag time. To estimate the transition probabilities, MSM employs a maximum likelihood approach where the likelihood function is:

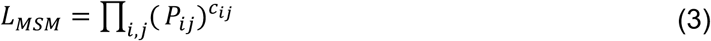

where *c_ij_* represents the observed transition counts between states. This yields the optimal transition probability matrix that best describes the observed dynamics, enabling the extraction of both thermodynamic and kinetic information. The following steps were conducted to build an MSM:

1. Featurization: All pairwise residue-residue distances were calculated to characterize peptide conformational changes during MD simulations.
2. Dimensionality reduction: Time-lagged independent component analysis (tICA) was employed to identify time-independent components (tICs) representing the slowest timescales.^49^ Each tIC is a linear combination of features.
3. Clustering: The K-means algorithm^50^ was applied to discretize MD simulation data into microstates. The MSM lag time, *τ*, was estimated as the minimum value at which the implied timescales converged in the implied timescale plot. (**Figure S1**). An appropriate lag time *τ* is chosen to maintain Markovianity, which means transition probabilities depend only on the current state and not on the prior history of the system.^20^
4. Hyperparameter optimization: The tIC dimension ranged from 2 to 10, and the number of microstates ranged from 100 to 700. These two hyperparameters were optimized by maximizing the VAMP-2 score (**Figure S2**). VAMP-2 score, calculated as the sum of squared eigenvalues of the transition probability matrix, maximizes the kinetic variance captured by the features.^51^ A 10-fold cross-validation was conducted to obtain the average score using pyEMMA.^52^ The selected MSM lag time, the tIC dimension, and the number of microstates for each lasso peptide were summarized in **Table S2**. Comparison of MSM-weighted and raw populations were shown in **Figure S3**. Stronger linear correlation indicates that MSM reweighting preserves the overall population distribution.

### Biased MD simulations: umbrella sampling

We performed umbrella sampling for biased MD simulations, using Q as the reaction coordinate. Initial configurations for umbrella sampling windows were selected from unbiased MD simulations. All frames from unbiased MD simulations were discretized into 50 evenly distributed bins, with Q ranging from 0 (unfolded) to 1 (pre-folded), and each bin served as an independent umbrella sampling window. We applied a harmonic potential for each window based on the root-mean-square deviation (RMSD) of heavy atoms relative to their reference structure. The restraint potential was defined as:

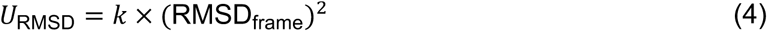

where *k* is the force constant set to 20 kcal·mol^−1^·Å^−2^, and RMSD_frame_ represents the instant RMSD of the current configuration compared to the reference structure. This restraint potential maintained each window near its designated region of conformational space while allowing sufficient fluctuations to achieve adequate sampling. The choice of force constant ensured sufficient overlap in sampling between adjacent windows for subsequent reconstruction of the free energy. The amount of biased umbrella sampling data for each lasso peptide is summarized in **Table S3**.

### Transition-based Reweighting Analysis Method (TRAM)

TRAM estimates a Multi-Ensemble Markov model (MEMM) across all simulation ensembles combining both biased and unbiased data, leverages the local equilibrium approximation of MSM, and benefits from biased simulations to enforce local equilibrium in interstate transitions that are difficult to attain.^23,24^ In TRAM, the conformational space is discretized into non-overlapping states, which are then classified into distinct ensembles. Each ensemble represents simulations performed under identical potential energy. Unbiased MD simulations were designated as a single ensemble, while each umbrella sampling window was treated as an independent ensemble. The interstate transitions follow:

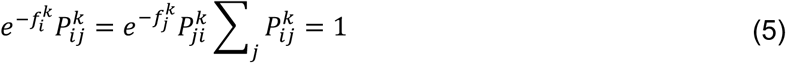

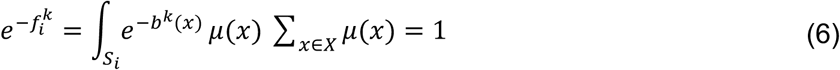

Where *f_i_^k^* is the local free energy of the *i* state and the *k* ensemble, *μ*(*x*) is the stationary density of the state *S_i_* in the ensemble *k*. *μ*(*x*) of each conformation *x* in the state *S_i_* in the ensemble *k* is weighted with the negative exponential of bias energy *b^k^*(*x*).

TRMA also employs the maximum likelihood approach, which combines the likelihood function of MSM and local equilibrium:

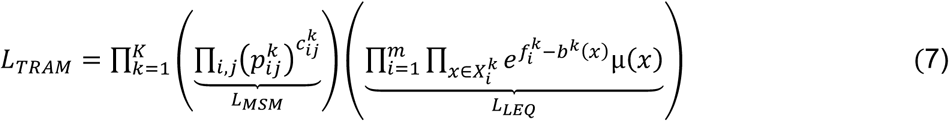

Wu *et al.* transformed this problem into a tractable system of nonlinear algebraic equations and solved it using an iterative algorithm.^24^ The implementation of TRAM consisted of the following steps:

1. Bias energy calculation: For each frame in both biased and unbiased trajectories, we calculated the bias potential energy. This resulted in a bias energy matrix with dimensions [*N*_frames_ × (*N*_windows_ + 1)] for each trajectory, where the additional column represents the unbiased ensemble with zero bias potential.
2. Thermodynamic state assignment: each frame was assigned to its corresponding thermodynamic state by constructing thermodynamic state trajectories (ttrajs). This assignment created a mapping between simulation frames and thermodynamic ensembles, accounting for a total of *N*_windows_ + 1 thermodynamic states.
3. Conformational state discretization: tICA was applied separately to both biased and unbiased trajectories with a lag time of *τ*. The biased tICs were used to transform unbiased data and vice versa, creating a combined feature space that captured the slowest process from both sampling regimes. Biased and unbiased tICs were concatenated and subsequently clustered using K-means clustering to construct discretized conformational states (dtrajs). The TRAM lag time was also estimated from the implied timescale plot (**Figure S6**). The selected lag time, the tIC dimension, and the number of microstates for each lasso peptide were summarized in **Table S3**.
4. TRAM implementation: the TRAM estimator from pyEMMA^52^ was applied to integrate the ttrajs, dtrajs, and biased energies to construct a MEMM.

### MD trajectory analysis

MD trajectory files were processed and analyzed using the CPPTRAJ in AMBERTools.^53^ Feature calculation was performed using MDTraj (version 1.9.0).^40^ Visual Molecular Dynamics (VMD) 1.9.3 was used to visualize the MD trajectories and generate snapshots.^54^ Secondary structure content was calculated using the Define Secondary Structure of Proteins (DSSP) algorithm^30^ implemented in MDTraj.^40^ Two-dimensional free energy plots were generated using the Python packages Matplotlib^55^ and Seaborn^56^.

### Error analysis for the percentage of pre-folded conformation

The error bar for the percentage of pre-folded conformation was calculated using bootstrapping with 100 iterations. In each iteration, 80% of the total number of trajectories were randomly selected for the equilibrium probability calculations.

### Folding free energy calculation

The folding free energy was calculated from the MEMM. From the converged MEMM, we extracted the full stationary distribution across all microstates. The pre-folded and unfolded states were defined by the *Q* and ring-closure distances (pre-folded state: Q *≥* 0.8 and ring-closure distance *≤* 6 Å, and unfolded state: Q *≤* 0.1 and ring closure distance *≥* 6 Å). The folding free energy Δ*G_f_* was computed using the reweighted equilibrium probabilities:

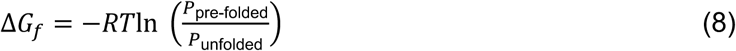

where *R* is the universal gas constant, *T* is the temperature, and the equilibrium probabilities are the sum of the weights for all frames belonging to that state. To estimate the statistical uncertainty of the calculated folding free energy, we performed bootstrap resampling with 100 iterations. In each iteration, 80% of the MD frames were randomly selected from the dataset for the equilibrium probability calculations.

### **Q**_loop_ relaxation time analysis

To quantify the stability of the loop region in each lasso peptide, we performed relaxation time analyses using the MEMM. The relaxation time characterizes the rate at which the loop equilibrates from its initial distribution toward the stationary state. The fraction of native contacts within the loop region, Q_loop_, was selected as the observable to monitor relaxation dynamics. The time evolution of the Q_loop_ expectation value was computed as follows:

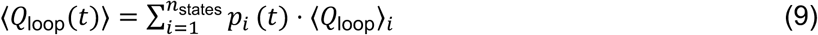

where *p*(*t*) is the probability of being in the state *i* at time *t*, obtained by:

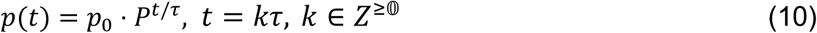

Here, *P* is the transition probability matrix of the MEMM, *τ* is the lag time of the MEMM, and *k* is a non-negative integer step index. The relaxation curve was computed for a sufficient time to observe convergence to the equilibrium. Q_loop_ relaxation time was defined as the time at which the observable reaches a plateau, detected automatically with a tolerance of 0.1% of the total signal change.

### Entropic cost calculation

To quantitatively assess the role of entropy in the folding dynamics of lasso peptides, we employed the PARENT program to calculate the conformational entropy for the pre-folded and unfolded structures.^35^ The PARENT program used the maximum information spanning tree (MIST) algorithm to compute conformational entropy.^57^ We randomly select 10000 frames from pre-folded (Q *≥* 0.8 and ring closure distance *≤* 6 Å) and unfolded (Q *≤* 0.1 and ring closure distance *≥* 6 Å) states for entropy calculation.

### Confinement simulations on capistruin

Using the pre-folded structure of capistruin, the solution system was first constructed and then prepared using the same procedures as previous unbiased MD simulations. During the production runs, a dummy spherical container was created to simulate the pocket confinement. An external force with a force constant of 5 kcal·mol^−1^·Å^−2^ was applied outside the sphere, while no force was applied within the sphere. We investigated radii of the dummy sphere ranging from 0.8 to 3 nm. For each system, we conducted 100 ns production runs with three replicates to estimate error. The average Q within 20 ns intervals was computed, after which capistruin reached structural stability across all replicates. The variation of Q for capistruin over 100 ns MD simulations under spherical confinement at different radii is shown in **Figure S11**.

### Transition Path Theory (TPT)

TPT is used to extract the transition pathways from unfolded to pre-folded states.^37,58^ After determining the unfolded (U) and pre-folded (F) states, the forward committor probability *q_i_*^+^ can be calculated from the transition probability matrix. The forward committor probability *q_i_*^+^ is the probability that, when the system is in the intermediate state *i*, it will next reach the pre-folded state rather than the unfolded state. Therefore, *q_i_*^+^ = 0 if *i* ∈ U, while *q_i_*^+^ = 1 if *q_i_*^+^ is calculated as follows:

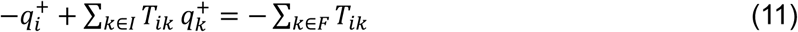

where *I* is a set of the intermediate states, *F* is a set of the pre-folded states, and *T* is the transition probability matrix.

Additionally, the backward committor probability *q_i_*^−^ is defined as the probability that, when the system is in the intermediate state *i*, it will next reach the unfolded state rather than the pre-folded state. Under the detailed balance condition of MSM and MEMM, the backward committor probability *q_i_*^−^ = 1 − *q_i_*^+^. The effective flux *f_ij_* is defined as the probability of proceeding through the intermediate states *i* and *j* during the transition from the unfolded to the pre-folded states. *f_ij_* is computed as follows:

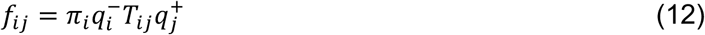

where *π_i_* is the stationary probability of the state *i*.

### Variational AutoEncoder (VAE)-based Latent space Path Clustering (LPC)

Each transition pathway consists of a sequence of microstates connecting the unfolded and pre-folded microstates. To dissect potential differences in folding mechanisms of different lasso peptides, the VAE-based LPC algorithm was applied to cluster kinetic pathways into distinct pathway channels.^25^ The LPC algorithm considers each pathway as a flow in the lower-dimensional subspaces and clusters the pathways based on their spatial distribution. Using microcin J25 as an example, the procedure for identifying pathway channels is described as follows:

1. Identify pathways from TPT: the average Q of each microstate was used to define the unfolded and pre-folded microstates for the TPT analysis. Microstates with an average Q *≥* 0.8 were defined as pre-folded, whereas those with an average Q ≤ 0.1 were defined as unfolded (when the maximum average Q was slightly below 0.8, the microstate with the highest average Q was assigned as pre-folded). TPT was employed to obtain an ensemble of 10,000 folding pathways for microcin J25. The cumulative flux as a function of the number of kinetic pathways is shown in **Figure S12A**.
2. Embed each pathway in a low-dimensional subspace: three pairs of tICs (tIC1-tIC2, tIC1-tIC3, and tIC2-tIC3), each representing different kinetics, were used as collective variables to construct three two-dimensional subspaces. Each subspace was discretized into 50 bins along each dimension, and every microstate was projected onto these subspaces. For each pathway, the projected subspaces of all microstates along that pathway were summed. The resulting three subspaces were concatenated into a single matrix, and the matrix was then flattened into a vector of length 7,500.
3. Train VAE: a VAE was trained on the vectors of 10,000 pathways with a split ratio of 0.25, batch size of 250, and learning rate of 8 × 10^−5^. After 100 training epochs, the model converged (**Figure S12B**) and generated a two-dimensional latent space.
4. Cluster pathways on the latent space: K-means clustering^50^ was applied to the latent space to classify the kinetic pathways into distinct pathway channels. The optimal number of clusters was determined by silhouette analysis.^39^

For the other lasso peptides, the cumulative flux and VAE training curves are summarized in **Figure S12C and S12D** for klebsidin, and in **Figures S13–S30** for the remaining 18 lasso peptides.

### General experimental materials and methods

The primers used to construct plasmids for microcin J25 variants were made by Integrated DNA Technologies. Primer sequences are listed in **Table S4**. The pLAM58 plasmid, which contains the microcin J25 peptidase, cyclase, and transporter, was provided by Lassogen. Q5 DNA polymerase, Dpn1, and NEBuilder HiFi DNA Assembly Master Mix were purchased from New England Biolabs. Chemical reagents were from Sigma Aldrich and used without further purification. Sanger sequencing was performed by the Core DNA Sequencing Facility at the University of Illinois at Urbana-Champaign. Matrix-assisted laser desorption/ionization time-of-flight mass spectrometry (MALDI-TOF-MS) data were acquired in reflector positive mode on a Bruker UltrafleXtreme instrument (Bruker Daltonics) at the University of Illinois School of Chemical Sciences Mass Spectrometry Laboratory. MALDI-TOF-MS data analysis was carried out using the Bruker FlexAnalysis software.

### Molecular biology techniques

Gibson assembly was used to construct the pET28-MBP-McjA plasmid. To make single-site variants, QuikChange site-directed mutagenesis or overlap extension PCR was used.^30^

### Preparation of *E. coli* lysate containing McjB, McjC, and McjD

pLAM58 was transformed into BL21 *E. coli* cells. A single colony was grown up in 5 mL of LB media containing 50 µg/µL kanamycin overnight at 37 °C with shaking. This culture was used to inoculate 500 mL of 2X YPTG media (10 g/L yeast extract, 16 g/L tryptone, 3 g/L KH_2_PO_4_, 7 g/L K_2_HPO_4_, 5 g/L NaCl, 18 g/L) containing 50 µg/µL kanamycin. The culture was allowed to grow at 37 °C while shaking. When the OD_600_ reached 0.5, a final concentration of 1 mM Isopropyl β-D-1-thiogalactopyranosid (IPTG) was added, and the culture was allowed to shake at 37 °C for an additional 2.5 h. The cells were spun down at 3,800 × g for 15 min at 4 °C. They were then washed two times in 200 mL cold S30A buffer [10 mM Tris, pH = 8.2, 14 mM magnesium acetate, 60 mM potassium glutamate, 2 mM 1,4-dithiothreitol (DTT) freshly added]. After washing, the cell pellet was resuspended in 30 mL S30A buffer and decanted into a pre-weighed 50 mL conical tube. The cells were centrifuged, the supernatant was removed, and the cell pellet was weighed, flash frozen in liquid nitrogen, and stored at −80 °C overnight. The next day, the cell pellet was resuspended in cold S30A buffer with freshly added DTT at 1 mL per g of cell pellet. The cells were lysed with a french press (single pass, 10,000 psi) and centrifuged at 24,000 × g for 45 min at 4 °C. The supernatant was removed and aliquoted before storing at −80 °C.

### Energy buffer preparation

Energy mix was prepared as described in Si. et al^59^.

### Expression and purification of GamS

GamS was prepared as described in Si. et al^59^.

### DNA template preparation for cell-free biosynthesis

Linear DNA templates were made from circular plasmids or overlap extension products using polymerase chain reaction and primers CFB-long-F and CFB-long-R (**Table S4**). The PCR products were examined on a gel and then purified using the Qiagen PCR clean-up kit per the manufacturer’s instructions. The concentration of the linear products was determined using the Nanodrop OneC instrument (Thermo Scientific), measuring absorbance at 260 nm.

### Production of variants in cell-free biosynthesis reactions

Reactions were carried out at a 10 µL volume. The reaction contained the following: 0.31 µL DTT (100 mM stock), 0.1 µL of IPTG (50 mM stock), 4.5 µL of energy buffer, 0.25 µL of GamS (125 µM stock), 3.5 µL of lysate, and 200 ng of template DNA. For competition experiments, 100 ng of wild-type DNA and 100 ng of variant DNA were added in place of the 200 ng template DNA. The reactions were allowed to sit at room temperature overnight. They were quenched with 60% acetonitrile, and then the debris was spun down at 8,000 × g for 8 min. 0.8 µL of the sample was spotted with 0.8 µL of a saturated solution of sinapic acid for analysis with MALDI-TOF-MS. To rank the variants by production level compared to the wild-type, the peak heights for the [M+H]^+^, [M+Na]^+^, and [M+K]^+^ were added together for the variant and for the wild-type. The sum of the variant peak heights was then divided by the sum of wild-type peak heights. Values greater than 1.2 were considered +, values between 0.8 and 1.19 were considered =, and values less than 0.79 were considered −. Variants with no production were marked with n.d. (not detected).

## Acknowledgement

This work was supported, in part, by grants from the National Institutes of Health (R35GM142745 and R21AI167693 to D.S. and R35GM158411 to D.A.M.). The authors thank the volunteers of folding@home for donating computing time. The authors also thank Dr. Soumajit Dutta, Dr. Prateek Bansal, and Dr. Austin Weigle for their help and discussion.

## Author Contributions

D.S. and D.A.M. conceived of the project and oversaw research progress. X.M performed the unbiased MD simulations. S.Y. performed the biased MD simulations. S.Y. and X.M performed computational data analysis. S.E.B. performed experiments and the corresponding data analysis. S.Y. and X.M. wrote the paper with help from S.E.B., and editing guidance from D.A.M. and D.S.

## Additional information

Supplementary Information and Source Data is available for this paper. Correspondence and requests for materials should be addressed to Douglas A. Mitchell or Diwakar Shukla.

