## Supplementary Information for "De novo Folding Mechanisms of Lasso Peptides"

**Table S1. Examined lasso peptides and their key attributes.** Core sequence, pre-folded ring-closure distance, Protein Data Bank (PDB) ID, and structure determination method of lasso peptides studied in molecular dynamics simulations. Ring residues are highlighted in red, loop residues are highlighted in pink, plug residues are highlighted in cyan, and tail residues are highlighted in blue. Pre-folded ring-closure distance denotes the distance between the N-terminus and the side chain carboxylate carbon of acceptor residue (Cδ of Glu or Cγ of Asp) of the pre-folded conformation. NMR represents solution Nuclear Magnetic Resonance spectroscopy, and XRD represents X-ray Diffraction.

| Lasso peptide | Core sequence | Pre-folded ring-closure distance (Å) | PDB ID | Structure determination method |
|---|---|---|---|---|
| acinetodin | GGKGPIFETWVTEGNYYG | 2.7 | 5UI6[1] | NMR |
| astexin-1 | GLSQGVEPDIGQTYFEESRINQD | 2.8 | 2LTI[2] | NMR |
| benenodin-1 | GVGFGRPDSILTQEQAKPMGLDRD | 2.8 | 5TJ1[3] | NMR |
| brevunsin | DGMGEEFIEGIVRDSLYPPAG | 2.7 | 5ZCN[4] | NMR |
| capistruin | GTPGFQTPDARVISRFGFN | 3.0 | 6N61[5] | XRD |
| caulonodin V | SIGDSGLRESMSSQTYWP | 3.8 | 2MLJ[6] | NMR |
| caulosegnin I | GAFVGQPEAVNPLGREIQG | 3.5 | 2LX6[7] | NMR |
| caulosegnin II | GTLTPGLPEDFLPGHYMPG | 3.1 | 5D9E[8] | XRD |
| chaxapeptin | GFGSKPLDSFGLNFF | 3.0 | 2N5C[9] | NMR |
| citrocin | GGVGKIIEYFIGGGVGRYG | 2.7 | 6MW6[10] | NMR |
| klebsidin | GSDGPIIEFFNPNGVMHYG | 2.6 | 5UI7[1] | NMR |
| microcin J25 | GGAGHVPEYFVGIGTPISFYG | 2.8 | 1Q71[11] | NMR |
| rubrivinodin | GAPSLINSEDNPAFPQRV | 2.9 | 5OQZ[12] | XRD |
| sphaericin | GLPIGWWIERPSGWYFPI | 2.7 | 5GVO[13] | NMR |
| sphingopyxin I | GIEPLGPVDEDQGEHYLFAGG | 2.8 | 5JQF[14] | XRD |
| streptomonomicin | SLGSSPYNDILGYPALIVIYP | 3.0 | 2MW3[15] | NMR |
| subterisin | GPPGDRIEFGVLAQLPGLDRD | 3.0 | 5XM4[16] | NMR |
| ubonodin | GGDGSIAEYFNRPMHIHDWQIMDSGYYG | 2.8 | 6POR[17] | NMR |
| xanthomonin I | GGPLAGEEIGGFNVPGISEE | 3.0 | 4NAG[18] | XRD |
| xanthomonin II | GGPLAGEEMGGITTLGISQD | 2.9 | 2MFV[18] | NMR |

**Table S2. Optimized hyperparameters for building Markov State Models (MSMs).** Optimized lag time, time-structure Independent Components (tIC) dimensions, and number of microstates for building MSMs.

| Lasso peptide | Lag time (ns) | tIC dimensions | Number of microstates |
|---|---|---|---|
| acinetodin | 50 | 4 | 400 |
| astexin-1 | 80 | 8 | 100 |
| benenodin-1 | 125 | 8 | 100 |
| brevunsin | 40 | 6 | 400 |
| capistruin | 60 | 10 | 100 |
| caulonodin V | 60 | 8 | 200 |
| caulosegnin I | 55 | 8 | 500 |
| caulosegnin II | 80 | 10 | 100 |
| chaxapeptin | 75 | 8 | 400 |
| citrocin | 60 | 8 | 200 |
| klebsidin | 100 | 10 | 400 |
| microcin J25 | 30 | 10 | 400 |
| rubrivinodin | 80 | 10 | 300 |
| sphaericin | 40 | 8 | 400 |
| sphingopyxin I | 65 | 6 | 700 |
| streptomonomicin | 60 | 10 | 200 |
| subterisin | 40 | 6 | 100 |
| ubonodin | 55 | 6 | 200 |
| xanthomonin I | 70 | 10 | 300 |
| xanthomonin II | 50 | 10 | 400 |

**Table S3. Optimized hyperparameters for building MultiEnsemble Markov Models (MEMMs).** Umbrella Sampling data, lag time, tIC dimensions, and number of microstates for building MEMMs using Transition-based Reweighting Analysis Method.

| Lasso peptide | Umbrella Sampling data (μs) | Lag time (ns) | tIC dimensions | Number of microstates |
|---|---|---|---|---|
| acinetodin | 7.60 | 15 | 4 | 400 |
| astexin-1 | 7.60 | 15 | 8 | 100 |
| benenodin-1 | 7.60 | 15 | 8 | 100 |
| brevunsin | 7.60 | 15 | 6 | 100 |
| capistruin | 10.7 | 10 | 10 | 100 |
| caulonodin V | 7.60 | 15 | 8 | 200 |
| caulosegnin I | 7.60 | 15 | 8 | 500 |
| caulosegnin II | 7.60 | 15 | 10 | 100 |
| chaxapeptin | 10.6 | 15 | 8 | 300 |
| citrocin | 7.60 | 15 | 8 | 200 |
| klebsidin | 7.60 | 15 | 10 | 400 |
| microcin J25 | 7.60 | 15 | 10 | 400 |
| rubrivinodin | 7.60 | 15 | 10 | 300 |
| sphaericin | 14.3 | 10 | 8 | 200 |
| sphingopyxin I | 15.7 | 15 | 6 | 500 |
| streptomonomicin | 7.60 | 15 | 10 | 200 |
| subterisin | 7.60 | 15 | 6 | 100 |
| ubonodin | 10.8 | 15 | 6 | 200 |
| xanthomonin I | 7.60 | 15 | 10 | 300 |
| xanthomonin II | 7.60 | 10 | 10 | 400 |

**Table S4. Oligonucleotide primers used in this study.** Sequences are listed 5' to 3'. F refers to a forward primer while R refers to a reverse primer. Mutations to the wild-type sequence are capitalized within the primer sequence.

| | CFB DNA template primers | Primer Sequence |
|---|---|---|
| 1 | CFB-long-F | gctatcatgccataccgcgaaaggttttgcgccattcg |
| 2 | CFB-long-R | aaccgtctatcagggcgatggcccactacgtgaaccatc |
| | **Cloning McjA primers** | |
| 3 | McjA-gib-F | gaacctgtacttccaatccggatccatgattaagcattttcattttaataaactg |
| 4 | McjA-gib-R | ctttgttagcagccggatctcagttcagccatagaaagatataggtg |
| 5 | pET28-gib-F | actgagatccggctgctaac |
| 6 | pET28-gib-R | ggatccggattggaagtacag |
| | **Overlap Extension Primers** | |
| 7 | McjA-10-12-OE-R | atactcaggcacatgtcctgc |
| 8 | McjA-F10G-F | gcaggacatgtgcctgagtatGGTgtggggattggtacacctatatc |
| 9 | McjA-V11W-F | caggacatgtgcctgagtattttTGGgggattggtacacctatatctttc |
| 10 | McjA-G12C-F | caggacatgtgcctgagtattttgtgTGCattggtacacctatatctttctatggctg |
| 11 | McjA-G12V-F | caggacatgtgcctgagtattttgtgGTTattggtacacctatatctttctatggctg |
| 12 | McjA-13-15-OE-R | ccccacaaaatactcaggcac |
| 13 | McjA-I13M-F | catgtgcctgagtattttgtggggatGggtacacctatatctttctatggctg |
| 14 | McjA-G14V-F | catgtgcctgagtattttgtggggattGTTacacctatatctttctatggctgaace |
| 15 | McjA-T15N-F | catgtgcctgagtattttgtggggattggtAACcctatatctttctatggctgaactgag |
| 16 | McjA-16-18-OE-R | tgtaccaatccccacaaaatactcag |
| 17 | McjA-P16R-F | ctgagtattttgtggggattggtacaCGTatatctttctatggctgaactgagatc |
| 18 | McjA-P16Q-F | ctgagtattttgtggggattggtacaCAGatatctttctatggctgaactgagatc |
| 19 | McjA-S18D-F | gtattttgtggggattggtacacctataGATttctatggctgaactgagatccg |
| 20 | McjA-21-OE-R | atagaaagatataggtgtaccaatccccac |
| 21 | McjA-G21F-F | ggggattggtacacctatatctttctatTTTgaactgagatccggctgctaac |
| | **Site-directed mutagenesis primers** | |
| 22 | McjA-P16T-F | ggattggtacaACCatatctttctatggctgaactgagatc |
| 23 | McjA-P16T-R | tagaaagatatGGTtgtaccaatccccacaaaatactcagg |
| 24 | McjA-S18V-F | gtacacctataGTTttctatggctgaactgagatccggctg |
| 25 | McjA-S18V-R | cagccatagaaAACtataggtgtaccaatccccacaaaata |
| 26 | McjA-S18P-F | gtacacctataCCGttctatggctgaactgagatccggctg |
| 27 | McjA-S18P-R | cagccatagaaCGGtataggtgtaccaatccccacaaaata |
| 28 | McjA-I17D-F | ttggtacacctGATtctttctatggctgaactgagatccgg |
| 29 | McjA-I17D-R | ccatagaaagaATCaggtgtaccaatccccacaaaatactc |
| 30 | McjA-T15I-F | gtggggattggtATTcctatatctttctatggctgaactgag |
| 31 | McjA-T15I-R | gaaagatataggAATaccaatccccacaaaatactcaggcac |
| 32 | McjA-V11K-F | ctgagtattttAAGgggattggtacacctatatctttctat |
| 33 | McjA-V11K-R | gtaccaatcccCTTaaaatactcaggcacatgtcctgcacc |

**Table S5. Table of selected microcin J25 loop variants ranked by β-strand content.** The varied residues in the sequence are shown in bold. Production levels were determined using the competition CFB experiments and are relative to wild-type. Specifically, the production level was determined by dividing the sum of the peak intensities for the cyclized variant [M+H]+, [M+Na]+, and [M+K]+ adducts by the sum of the peak intensities for the wild-type [M+H]+, [M+Na]+, and [M+K]+ adducts. Production values greater than 1.2 are denoted +, values between 0.8 and 1.19 are =, and values less than 0.79 are −. Variants that were not detected are n.d.

| Variant | Sequence | CFB Production level relative to WT | S4PRED[19] predicted β-strand content | S4PRED[19] prediction | MD predicted β-strand content (mean ± SD) | MD prediction |
|---|---|---|---|---|---|---|
| WT | GGAGHVPEYFVGIGTPISFYG | | 1.837 | | 7.32 ± 0.01 | |
| S18P | GGAGHVPEYFVGIGTPI**P**FYG | n.d. | 0.947 | break | 1.99 ± 0.01 | break |
| V11K | GGAGHVPEYF**K**GIGTPISFYG | - | 0.457 | break | 2.15 ± 0.02 | break |
| G12V | GGAGHVPEYFV**V**IGTPISFYG | + | 5.104 | enhance | 3.50 ± 0.02 | break |
| G12C | GGAGHVPEYFV**C**IGTPISFYG | - | 4.427 | enhance | 3.63± 0.02 | break |
| P16R | GGAGHVPEYFVGIGT**R**ISFYG | + | 4.377 | enhance | 4.56 ± 0.02 | break |
| I17D | GGAGHVPEYFVGIGTP**D**SFYG | n.d. | 0.791 | break | 5.18 ± 0.02 | break |
| F10G | GGAGHVPEY**G**VGIGTPISFYG | - | 1.145 | break | 5.21 ± 0.02 | break |
| I13M | GGAGHVPEYFVG**M**GTPISFYG | + | 1.087 | break | 8.87 ± 0.01 | enhance |
| P16Q | GGAGHVPEYFVGIGT**Q**ISFYG | = | 4.496 | enhance | 9.30 ± 0.01 | enhance |
| T15N | GGAGHVPEYFVGIG**N**PISFYG | + | 1.121 | break | 9.42 ± 0.01 | enhance |
| P16T | GGAGHVPEYFVGIGT**T**ISFYG | - | 4.761 | enhance | 10.01 ± 0.01 | enhance |
| S18V | GGAGHVPEYFVGIGTPI**V**FYG | - | 3.636 | enhance | 10.01 ± 0.01 | enhance |
| S18D | GGAGHVPEYFVGIGTPI**D**FYG | - | 1.186 | break | 10.02 ± 0.01 | enhance |
| V11W | GGAGHVPEYF**W**GIGTPISFYG | + | 1.128 | break | 10.03 ± 0.01 | enhance |
| T15I | GGAGHVPEYFVGIG**I**PISFYG | + | 3.846 | enhance | 10.04 ± 0.01 | enhance |
| G14V | GGAGHVPEYFVGI**V**TPISFYG | + | 3.594 | enhance | 10.06 ± 0.01 | enhance |

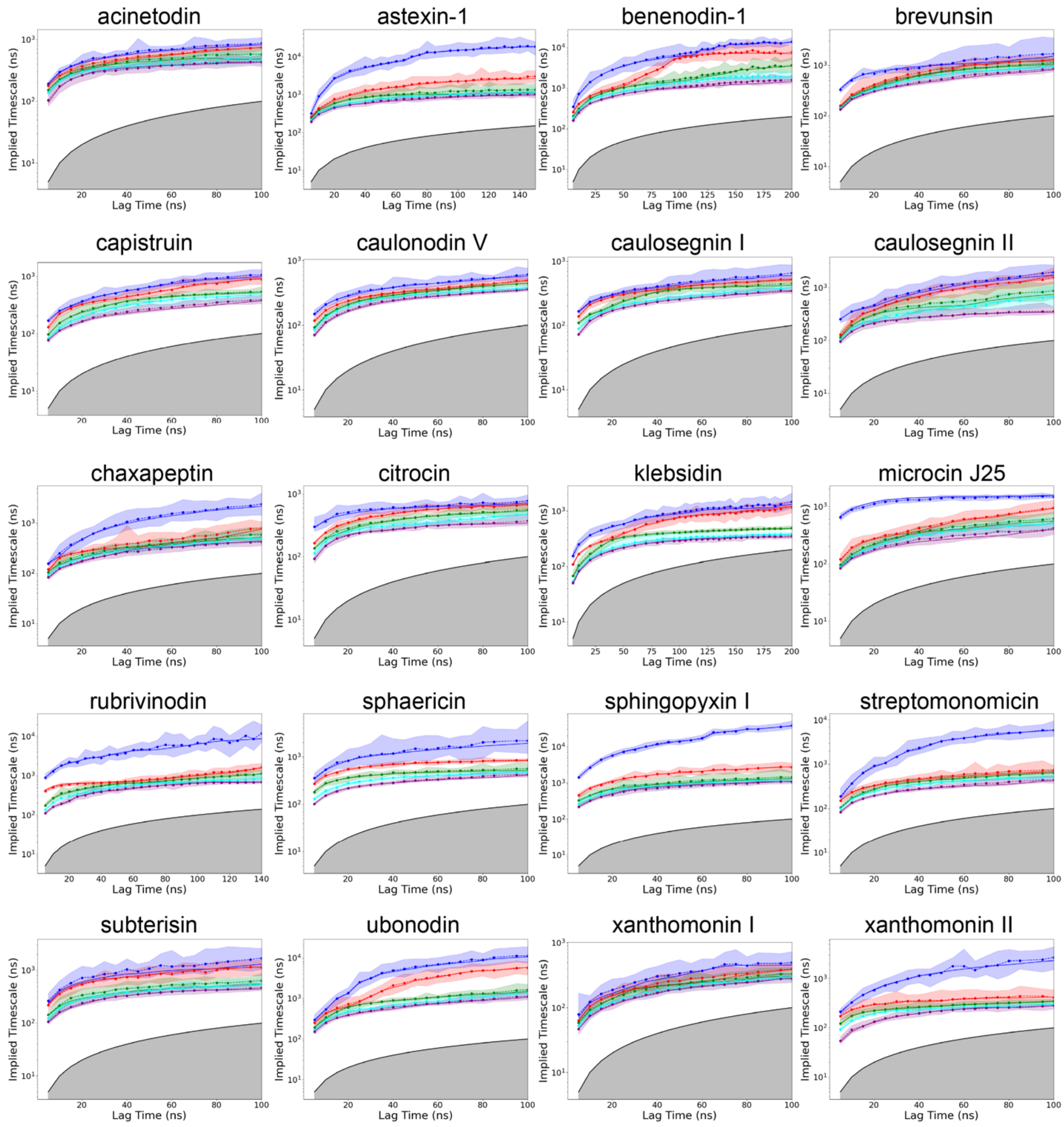

**Figure S1. Implied timescale analysis for MSM lag time selection.** The five slowest implied timescales are plotted as a function of lag time for each system, with colors ranking the processes from slowest to fastest (blue, red, green, cyan and purple). Convergence of the implied timescales indicates the chosen lag time for MSM construction in Table S2. The shaded regions around each line represent 95% confidence intervals estimated via Bayesian sampling of the transition matrix. The grey-shaded area below the black line, where the implied timescale equals the lag time, corresponds to processes faster than the lag time and are not resolved by the MSM.

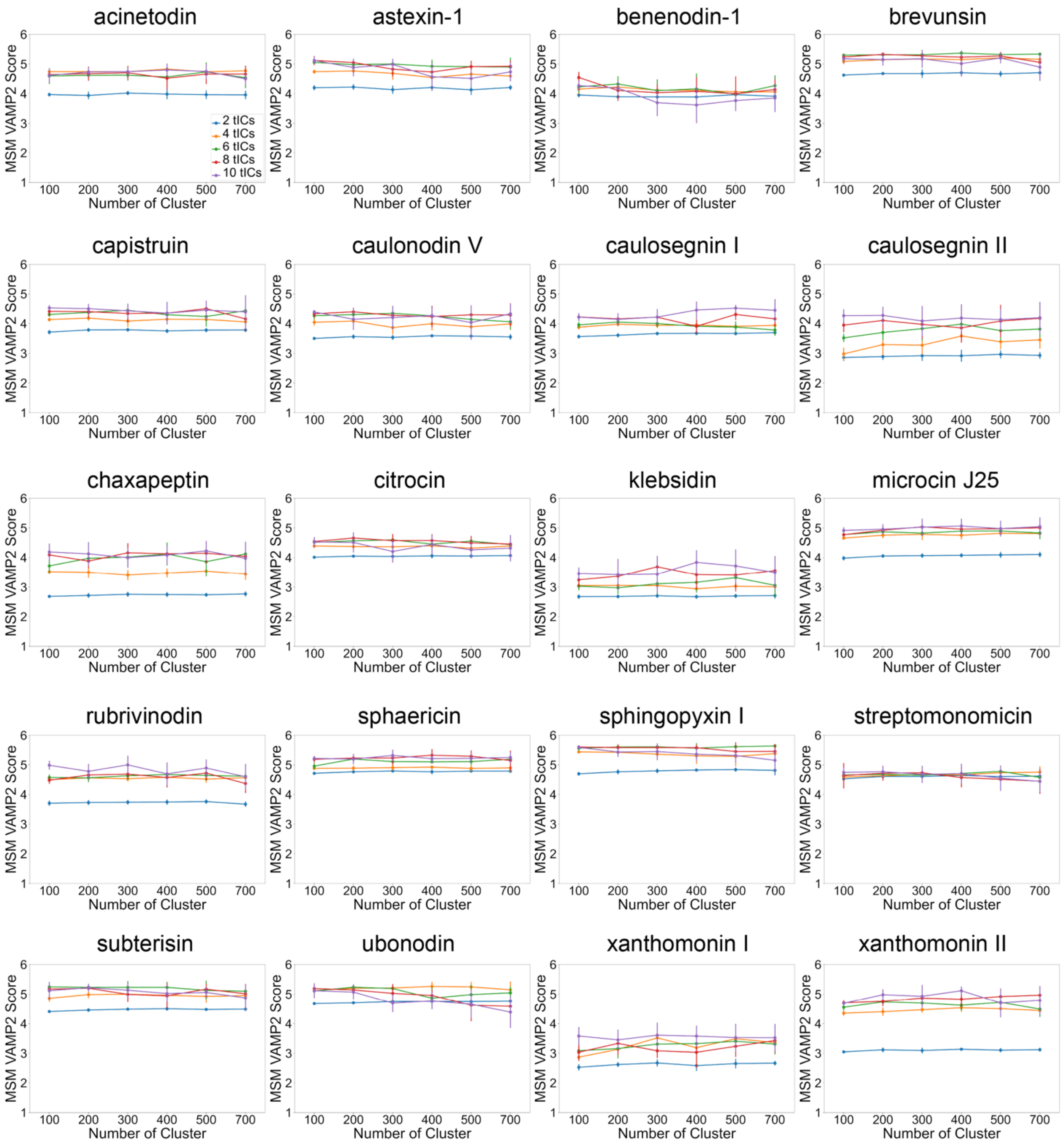

**Figure S2. MSM hyperparameter optimization.** Variational Approach for Markov Processes (VAMP-2) scores are shown as a function of the number of clusters and tIC dimensions for each system. Higher VAMP-2 scores indicate better resolution of slow kinetic processes. The optimal number of clusters and tIC dimension were selected by maximizing the VAMP-2 score and are reported in Table S2.

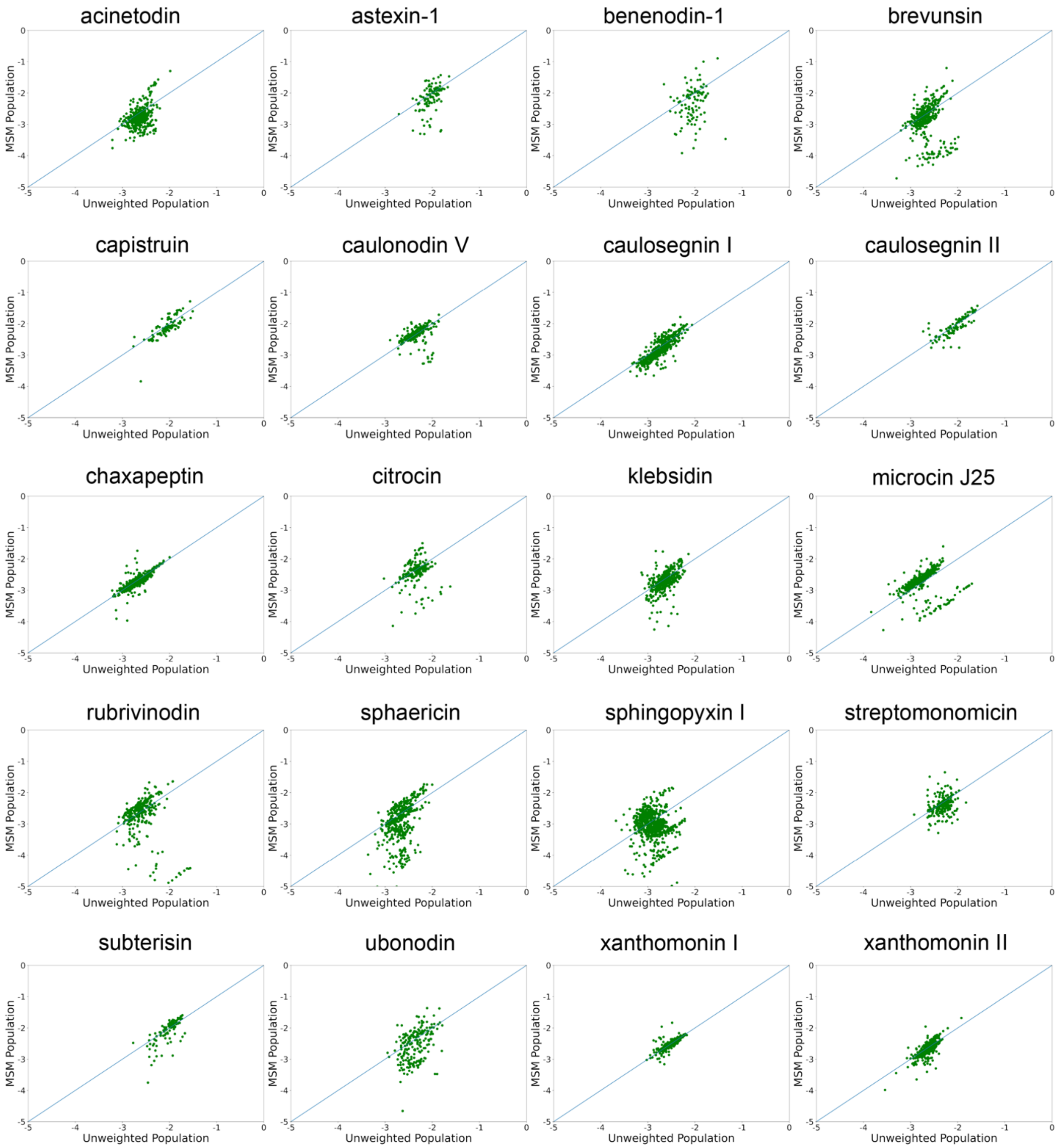

**Figure S3. Comparison of MSM-weighted and raw populations for each microstate.** Stronger linear correlation indicates that MSM reweighting preserves the overall population distribution.

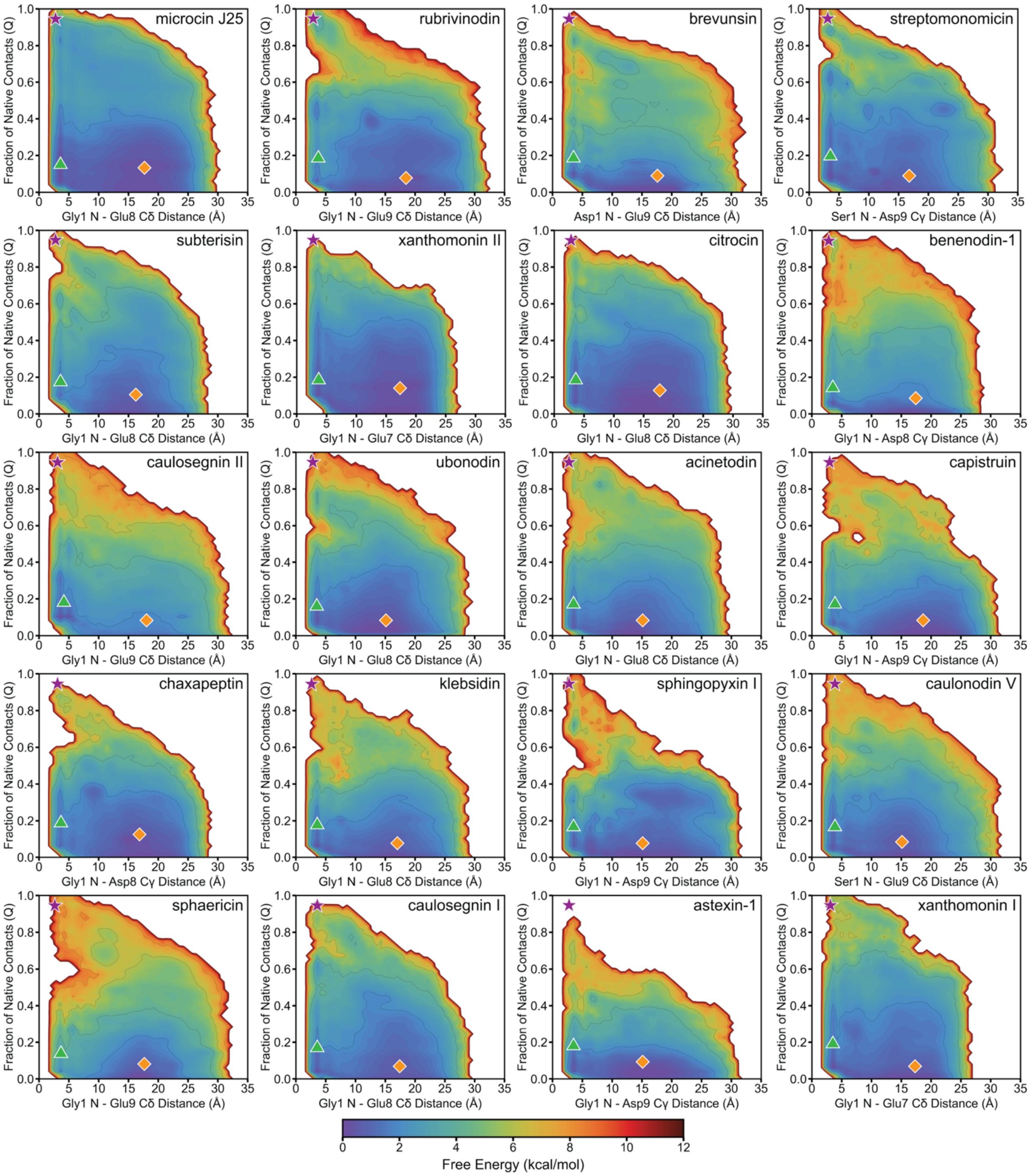

**Figure S4. MSM-weighted two-dimensional free energy landscapes.** The following representative conformations are marked: pre-folded (purple star), non-threaded proto-folded (green triangle), and unfolded (orange diamond). Panels are arranged in the same order as the MEMM-weighted two-dimensional free energy landscapes (Figure 3).

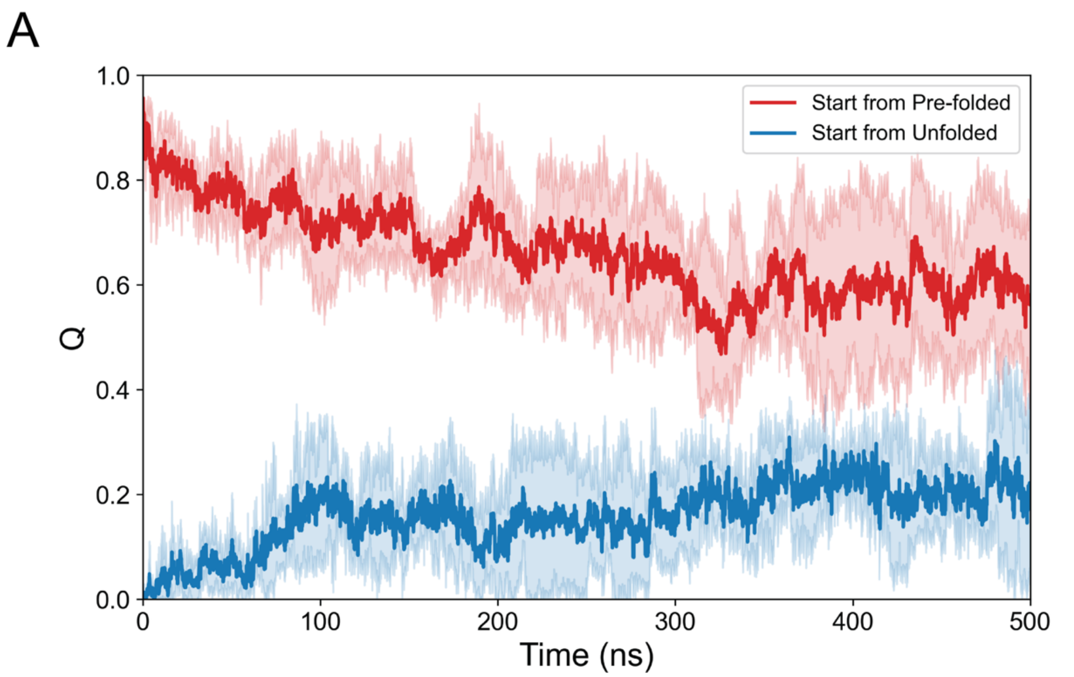

**Figure S5. Fraction of native contacts (Q) variation during unbiased MD simulations of microcin J25 and capistruin. 500 ns** trajectories initiated from the pre-folded structure are shown in red, whereas those initiated from the unfolded structure are shown in blue. Solid lines represent the mean Q values, and shaded regions denote the standard deviation across five independent replicates.

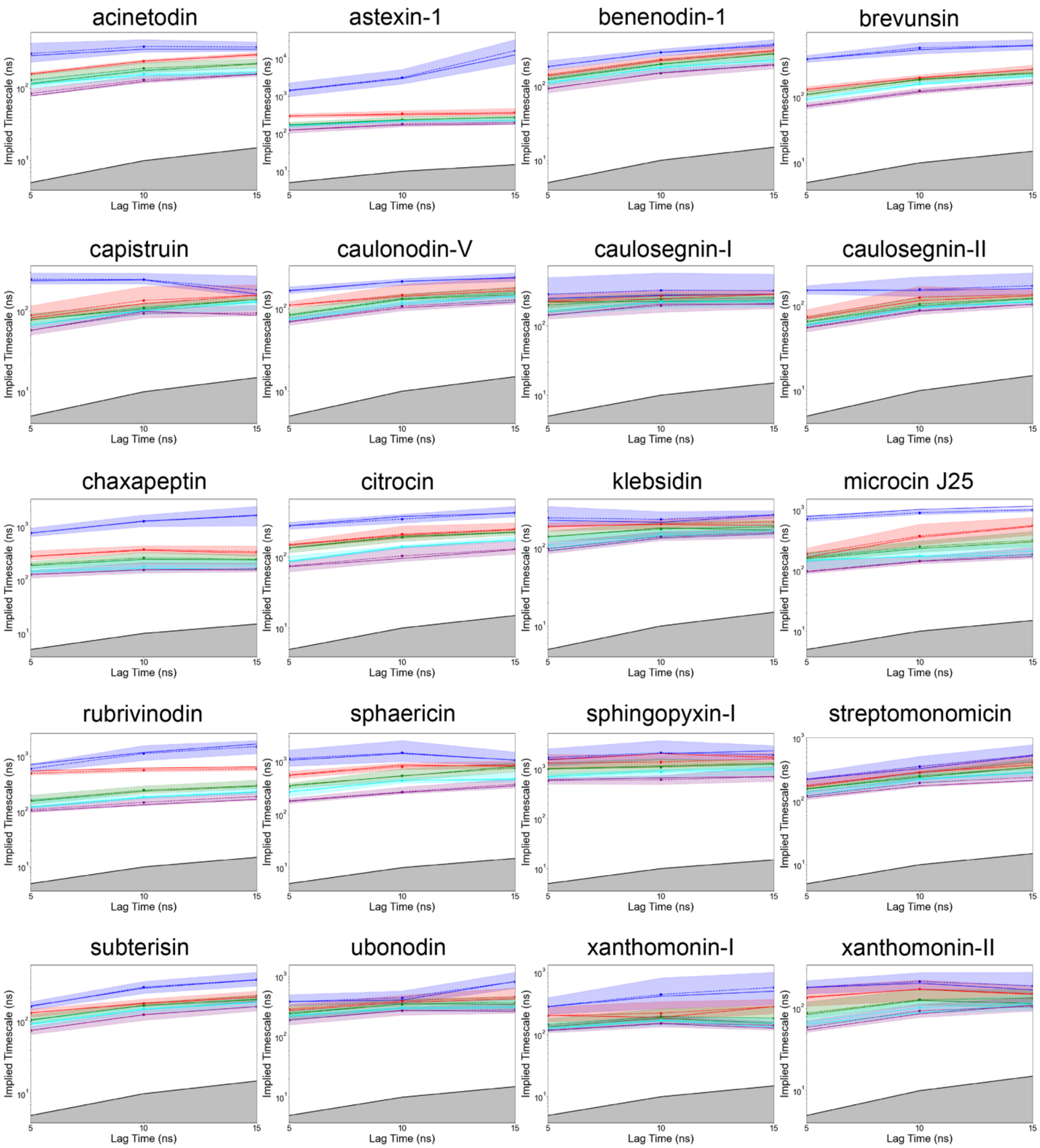

**Figure S6. Implied timescale analysis for MEMM lag time selection.** The five slowest implied timescales are plotted as a function of lag time for each system, with colors ranking the processes from slowest to fastest (blue, red, green, cyan and purple). Convergence of the implied timescales indicates the chosen lag time for MEMM construction in Table S3. The shaded regions around each line represent 95% confidence intervals estimated via Bayesian sampling of the transition matrix. The grey-shaded area below the black line, where the implied timescale equals the lag time, corresponds to processes faster than the lag time and are not resolved by the MEMM.

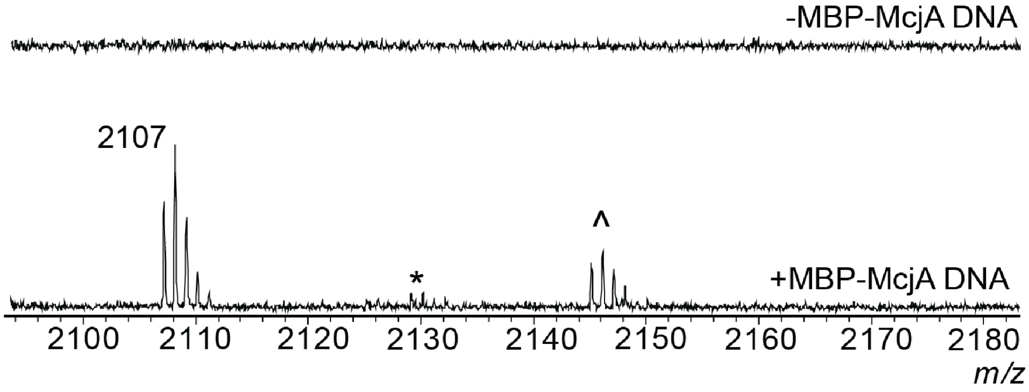

**Figure S7. Microcin J25 production in lysate pre-expressing McjBCD.** The mass indicates the $[M+H]^+$ peak, * indicates the $[M+Na]^+$ peak, and ˆ indicates the $[M+K]^+$ peak.

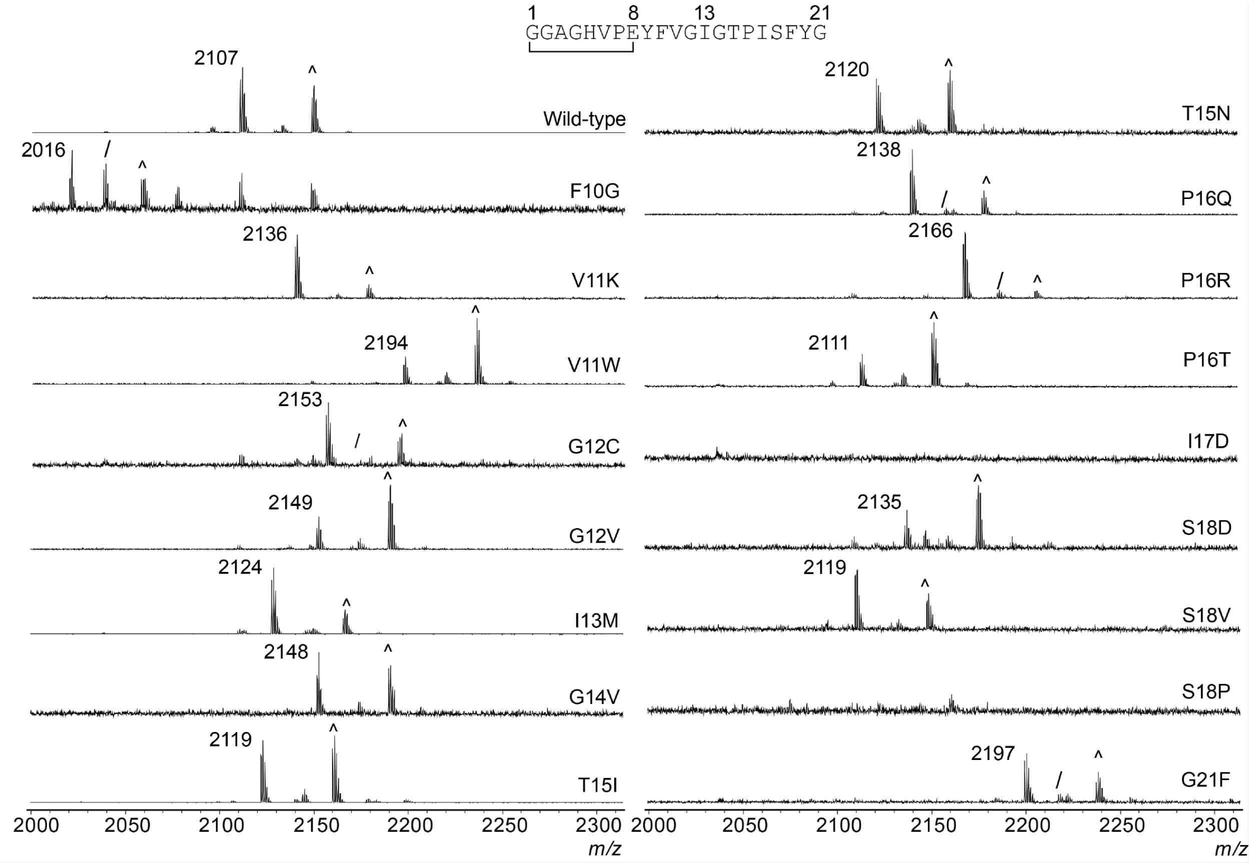

**Figure S8. Microcin J25 variant production in lysate pre-expressed McjBCD.** Variant identity is indicated at the bottom right of each spectrum. The cyclized $[M+H]^+$ peak is indicated by the mass. The ^ indicates the $[M+K]^+$ peak, and / indicates the linear $[M+H]^+$ peak.

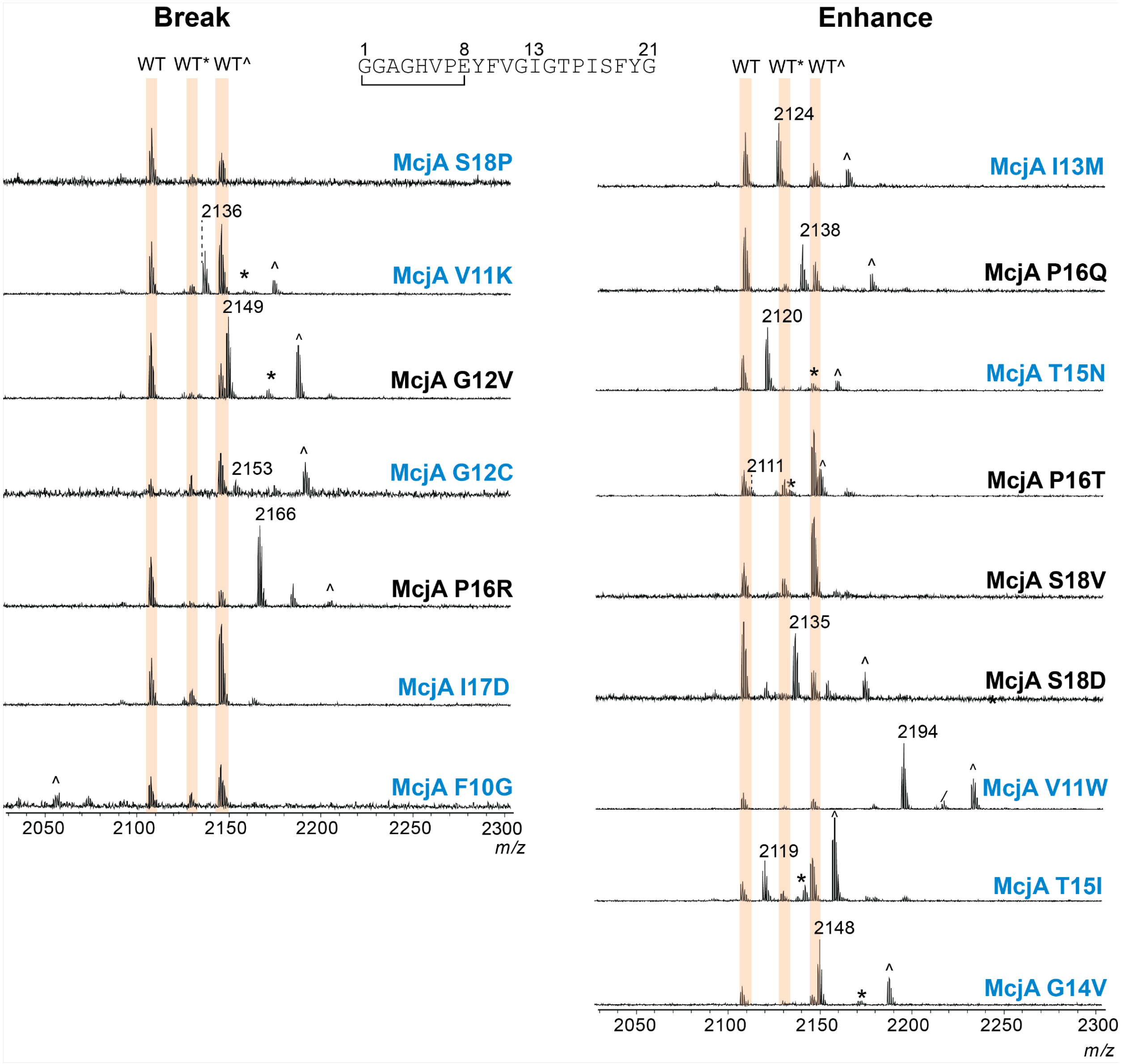

**Figure S9. Cell-free biosynthetic assessment of microcin J25 variants.** Competition assays between McjA wild-type and variants were assessed by MALDI mass spectrometry. Peaks corresponding to the wild-type microcin J25 adducts ($[M+H]^+$, $[M+Na]^+$, and $[M+K]^+$) are highlighted in orange. The cyclized $[M+H]^+$ peak for the variant peptide is indicated by the mass. The * indicates the cyclized $[M+Na]^+$ peak, the ^ indicates the cyclized $[M+K]^+$ peak, and the / indicates the linear $[M+H]^+$ peak. The spectra are split into break (left) and enhance (right) β-hairpin content based on MD simulation data. A blue label indicates that the data aligns with the MD-predicted β-hairpin content.

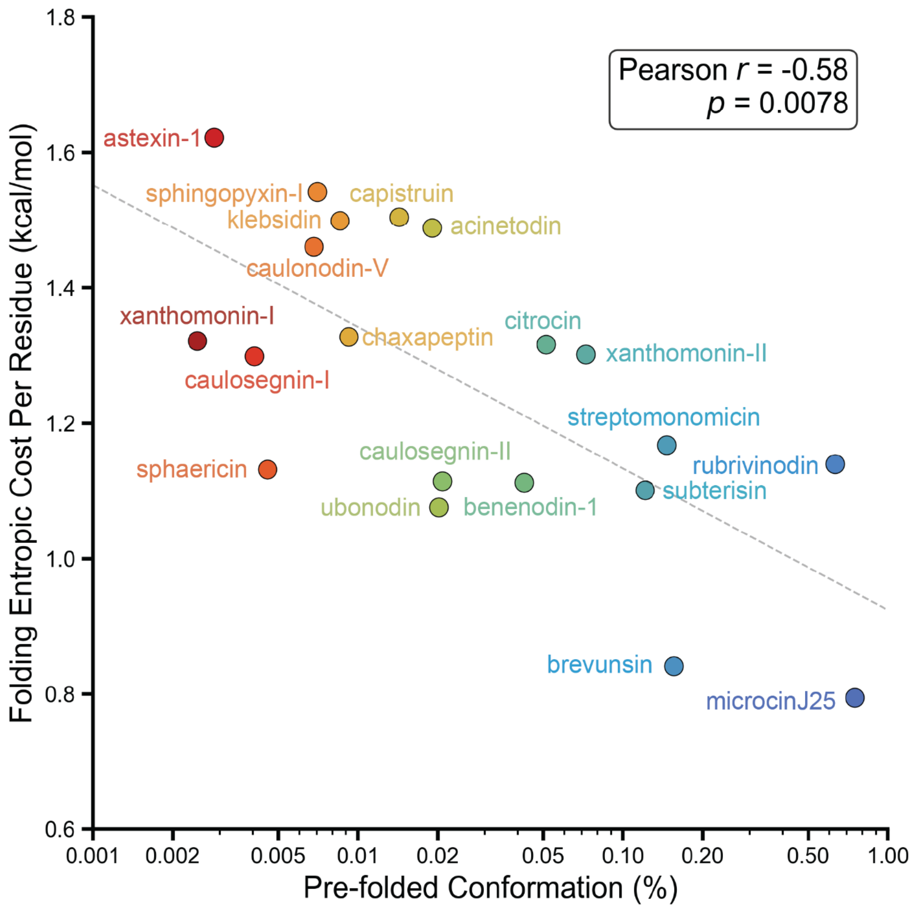

**Figure S10. Correlations between entropy cost per residue and the probability of pre-folded conformation.** The entropy cost per residue is defined as the total entropic cost normalized by the length of the lasso peptide.

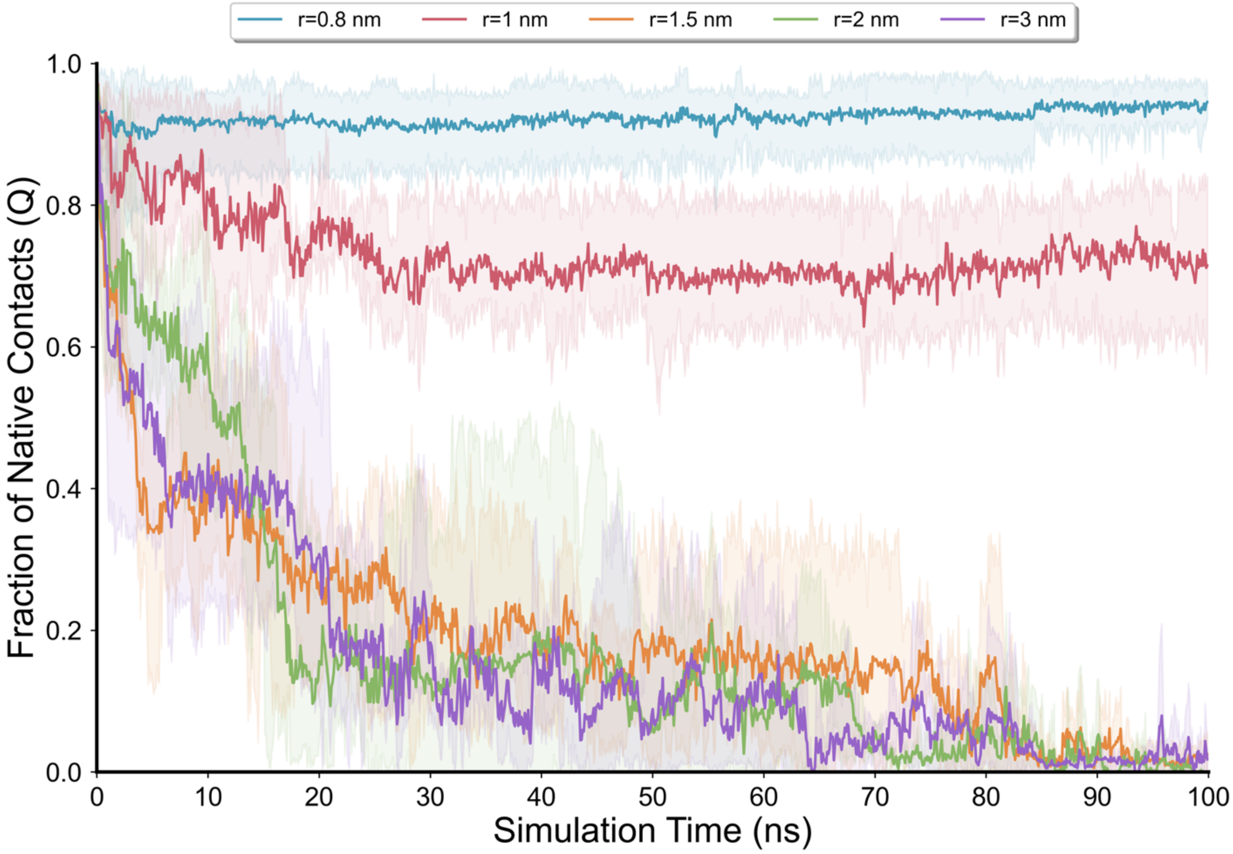

**Figure S11. Q variation during the confinement MD simulations of capistruin.** A 100 ns MD simulation of capistruin was conducted, with spherical confinement assessed at five different radii (0.8-3.0 nm). Solid lines represent mean values, and shaded regions indicate the min-max range across three independent replicates.

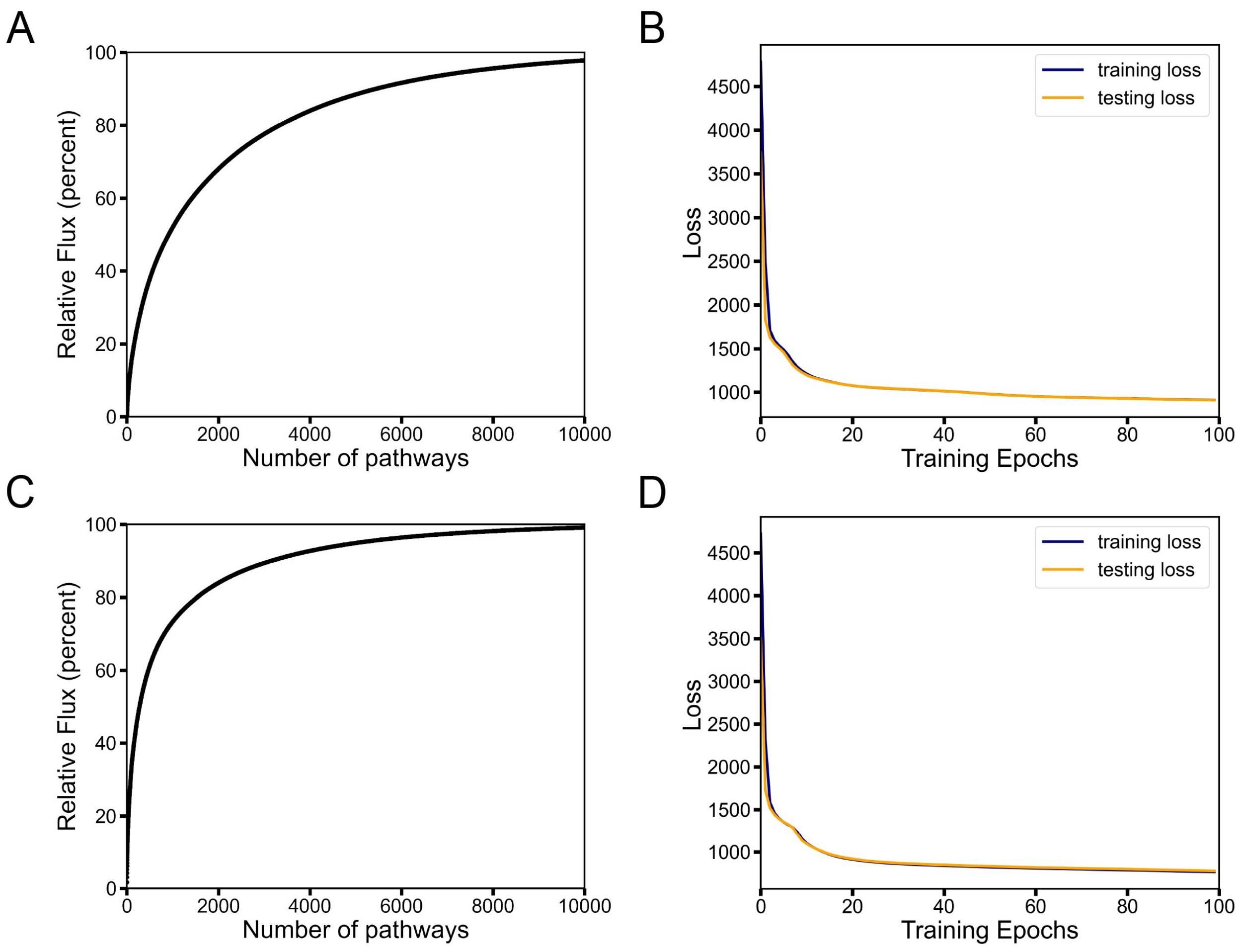

**Figure S12. The cumulative flux and Variational AutoEncoder (VAE) training curves for microcin J25 and klebsidin.** (A) Cumulative flux as a function of the number of kinetic pathways for microcin J25. (B) Loss function vs training epochs of VAE models trained on kinetic folding pathways of microcin J25. (C) Cumulative flux as a function of the number of kinetic pathways for klebsidin. (D) Loss function vs training epochs of VAE models trained on kinetic folding pathways of klebsidin.

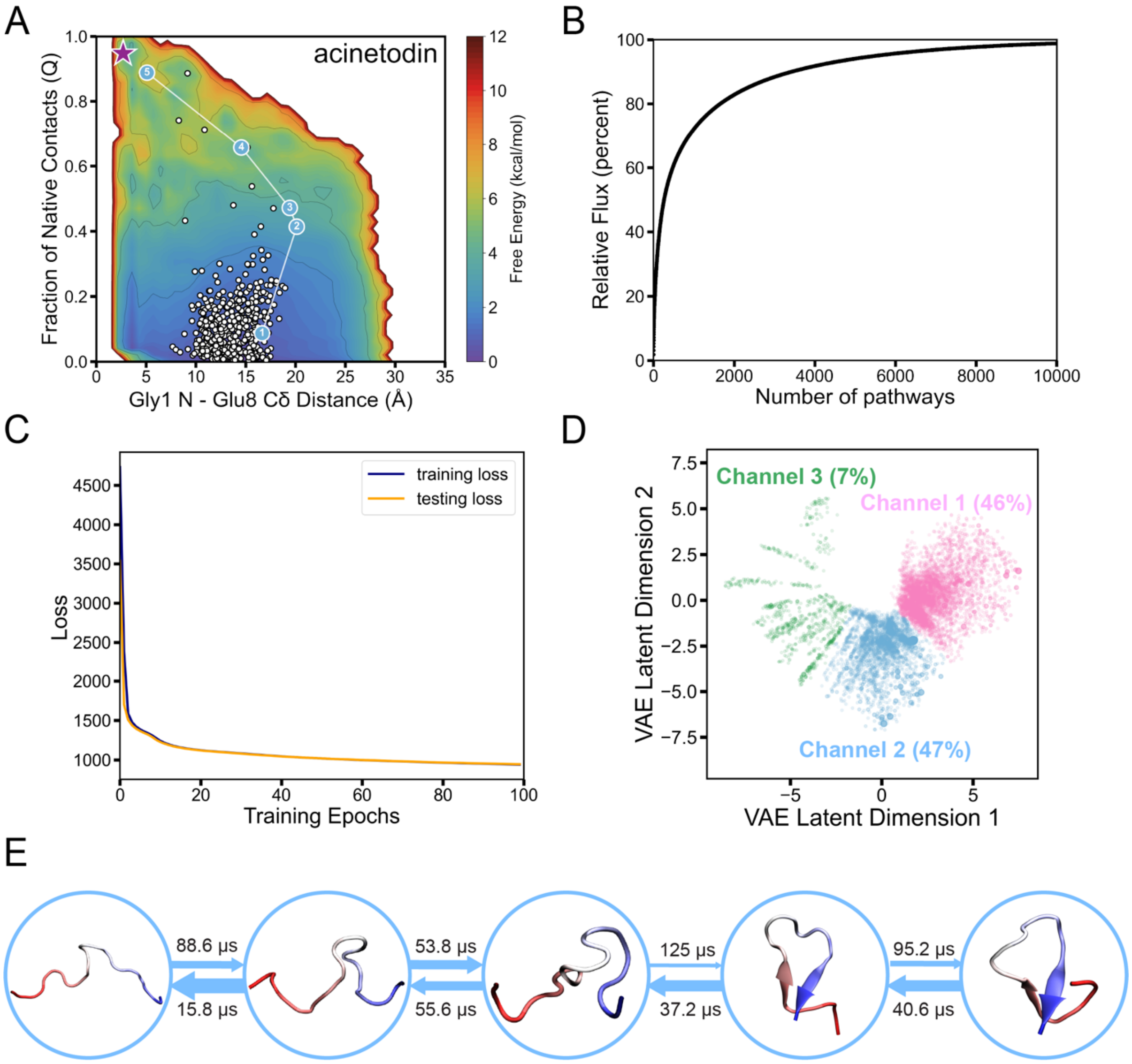

**Figure S13. Folding pathways of acinetodin.** (A) Distribution of microstate centers (white dots) on the TRAM-weighted free energy landscape, with the most representative folding pathway highlighted by its corresponding pathway channel color. The pre-folded conformation is marked with a purple star. (B) Cumulative flux as a function of the number of kinetic pathways. (C) Loss function vs training epochs of VAE. (D) Kinetic pathways and pathway channels in the latent space captured by the VAE-based LPC algorithm. Each point in the latent space corresponds to a single pathway, and its size is determined by its normalized flux value. Each pathway channel was color-labeled with its relative percentage of the total flux. (E) Structural details of the most representative folding pathway, with the mean first passage time (MFPT) calculated between every two adjacent states. The N-terminus of the lasso peptide is shown in red, and the C-terminus of the lasso peptide is shown in blue.

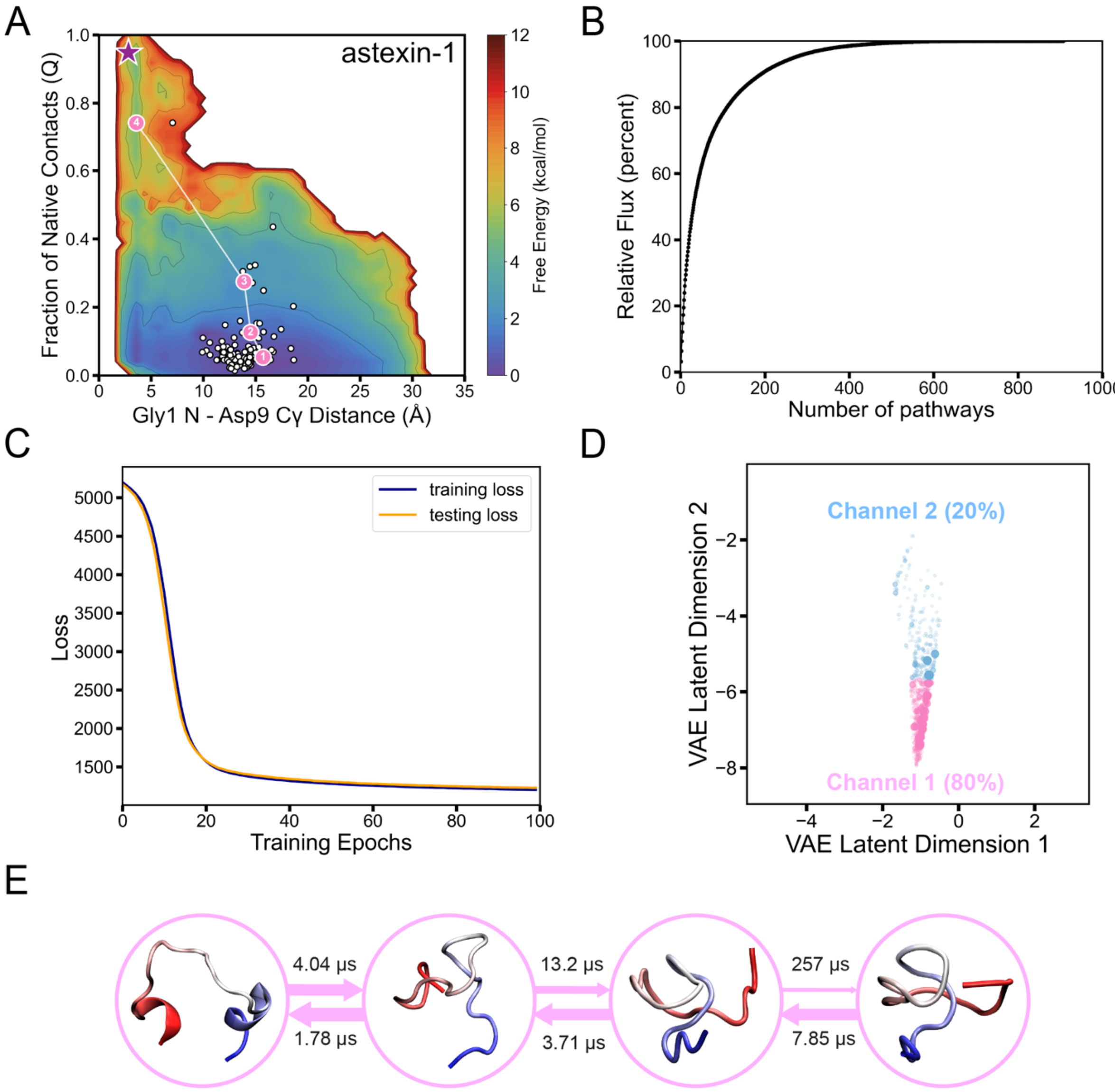

**Figure S14. Folding pathways of astexin-1.** (A) Distribution of microstate centers (white dots) on the TRAM-weighted free energy landscape, with the most representative folding pathway highlighted by its corresponding pathway channel color. The pre-folded conformation is marked with a purple star. (B) Cumulative flux as a function of the number of kinetic pathways. (C) Loss function vs training epochs of VAE. (D) Kinetic pathways and pathway channels in the latent space captured by the VAE-based LPC algorithm. Each point in the latent space corresponds to a single pathway, and its size is determined by its normalized flux value. Each pathway channel was color-labeled with its relative percentage of the total flux. (E) Structural details of the most representative folding pathway, with the MFPT calculated between every two adjacent states. The N-terminus of the lasso peptide is shown in red, and the C-terminus of the lasso peptide is shown in blue.

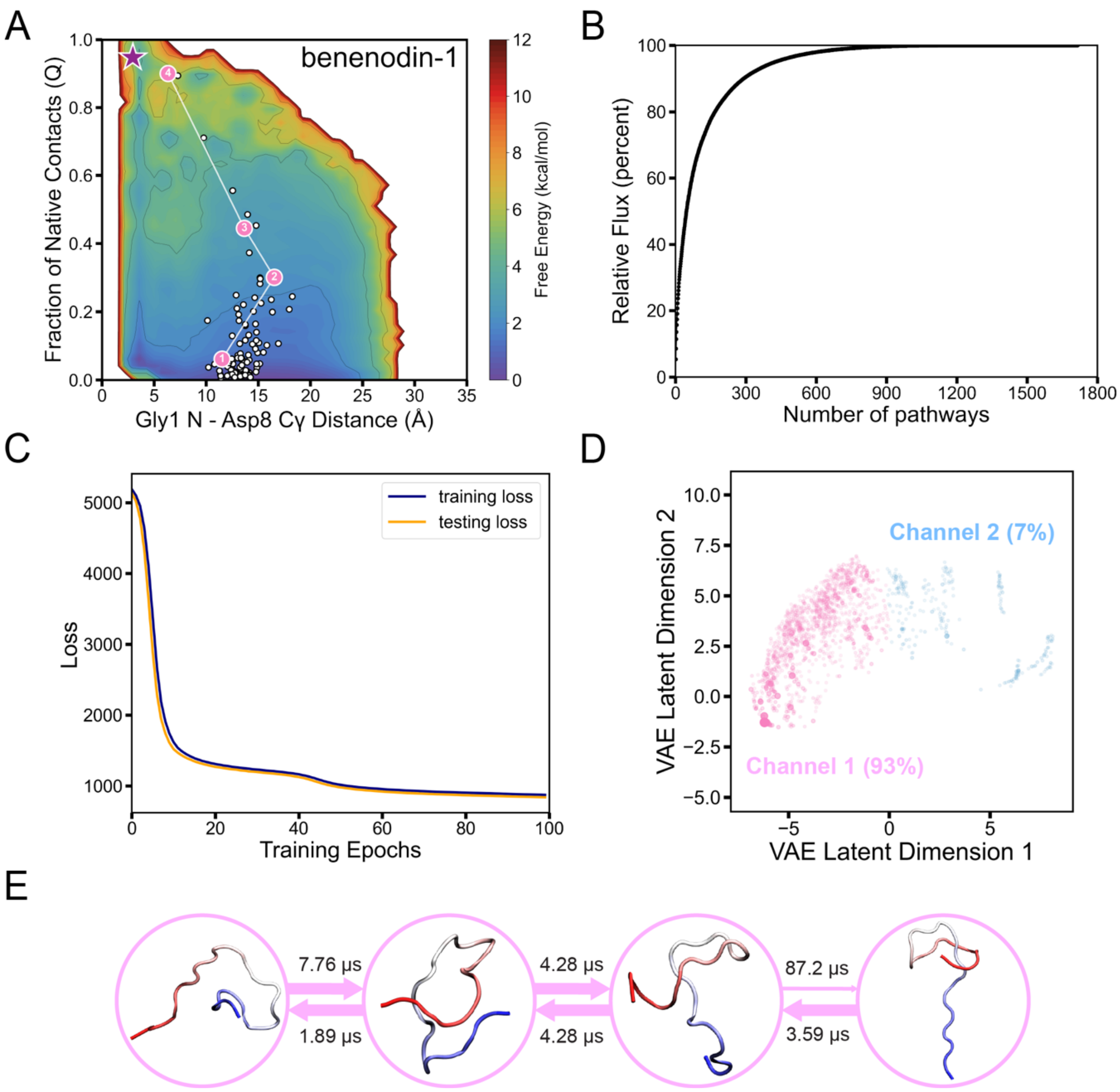

**Figure S15. Folding pathways of benenodin-1.** (A) Distribution of microstate centers (white dots) on the TRAM-weighted free energy landscape, with the most representative folding pathway highlighted by its corresponding pathway channel color. The pre-folded conformation is marked with a purple star. (B) Cumulative flux as a function of the number of kinetic pathways. (C) Loss function vs training epochs of VAE. (D) Kinetic pathways and pathway channels in the latent space captured by the VAE-based LPC algorithm. Each point in the latent space corresponds to a single pathway, and its size is determined by its normalized flux value. Each pathway channel was color-labeled with its relative percentage of the total flux. (E) Structural details of the most representative folding pathway, with the MFPT calculated between every two adjacent states. The N-terminus of the lasso peptide is shown in red, and the C-terminus of the lasso peptide is shown in blue.

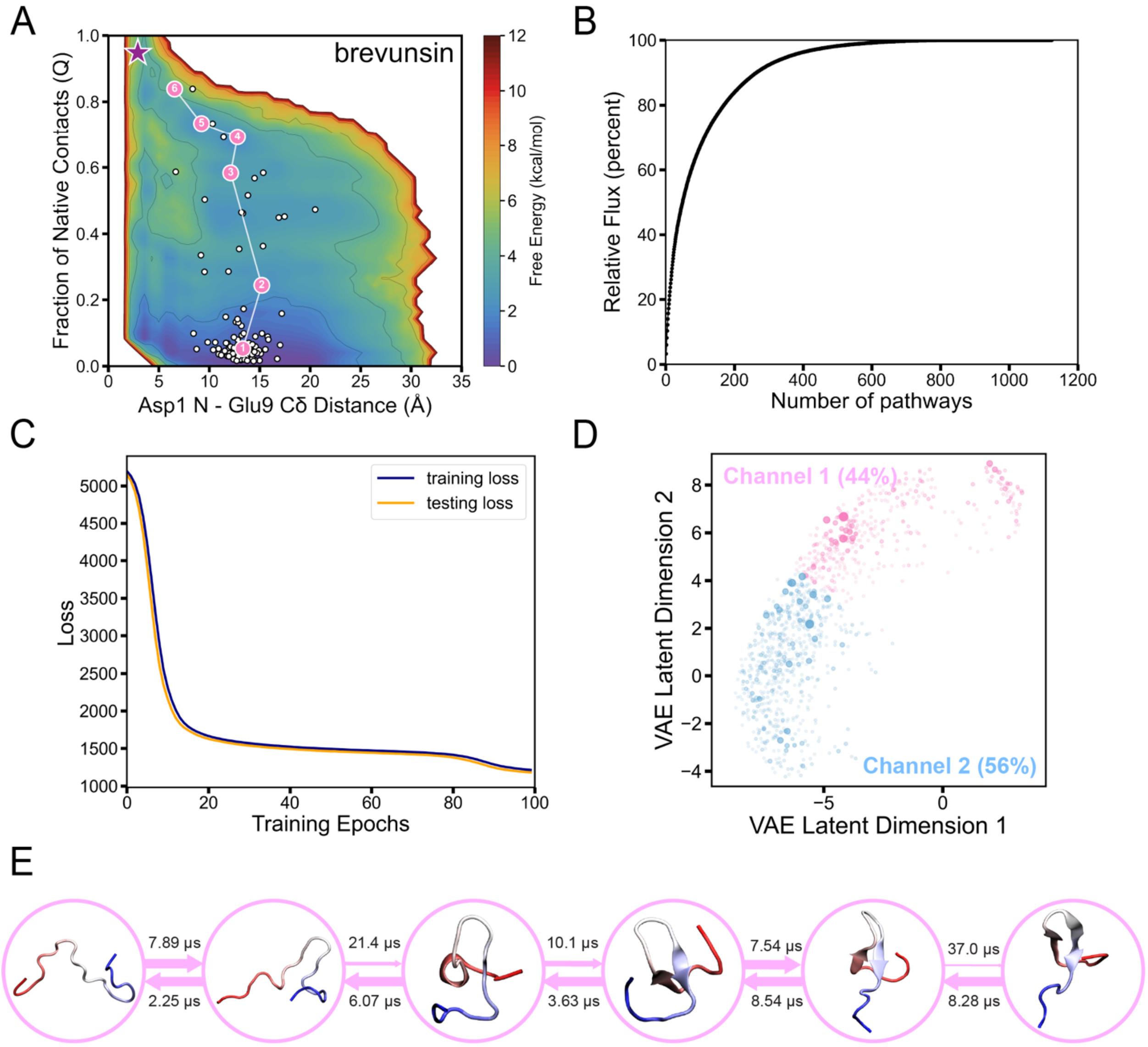

**Figure S16. Folding pathways of brevunsin.** (A) Distribution of microstate centers (white dots) on the TRAM-weighted free energy landscape, with the most representative folding pathway highlighted by its corresponding pathway channel color. The pre-folded conformation is marked with a purple star. (B) Cumulative flux as a function of the number of kinetic pathways. (C) Loss function vs training epochs of VAE. (D) Kinetic pathways and pathway channels in the latent space captured by the VAE-based LPC algorithm. Each point in the latent space corresponds to a single pathway, and its size is determined by its normalized flux value. Each pathway channel was color-labeled with its relative percentage of the total flux. (E) Structural details of the most representative folding pathway, with the MFPT calculated between every two adjacent states. The N-terminus of the lasso peptide is shown in red, and the C-terminus of the lasso peptide is shown in blue.

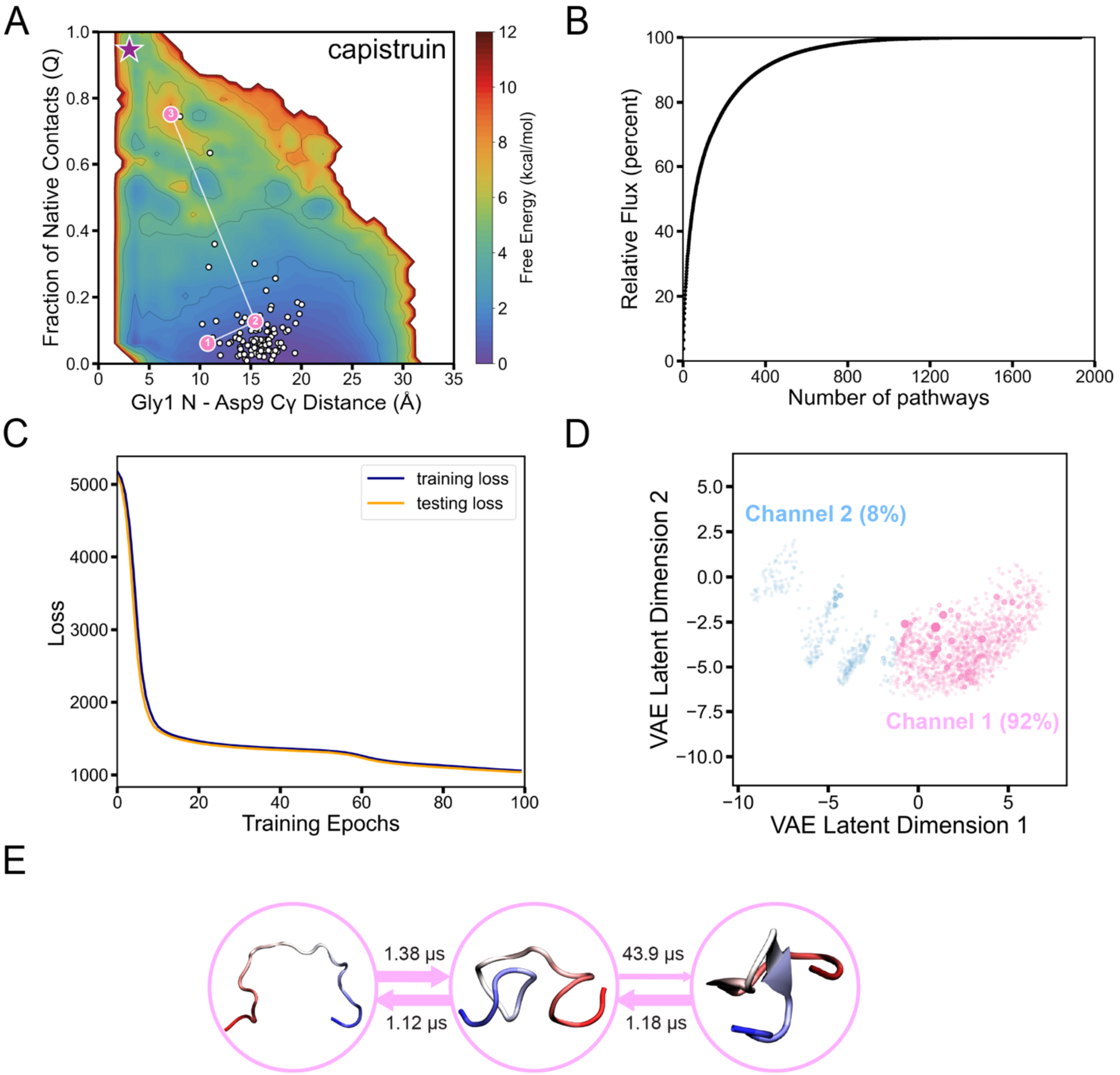

**Figure S17. Folding pathways of capistruin.** (A) Distribution of microstate centers (white dots) on the TRAM-weighted free energy landscape, with the most representative folding pathway highlighted by its corresponding pathway channel color. The pre-folded conformation is marked with a purple star. (B) Cumulative flux as a function of the number of kinetic pathways. (C) Loss function vs training epochs of VAE. (D) Kinetic pathways and pathway channels in the latent space captured by the VAE-based LPC algorithm. Each point in the latent space corresponds to a single pathway, and its size is determined by its normalized flux value. Each pathway channel was color-labeled with its relative percentage of the total flux. (E) Structural details of the most representative folding pathway, with the MFPT calculated between every two adjacent states. The N-terminus of the lasso peptide is shown in red, and the C-terminus of the lasso peptide is shown in blue.

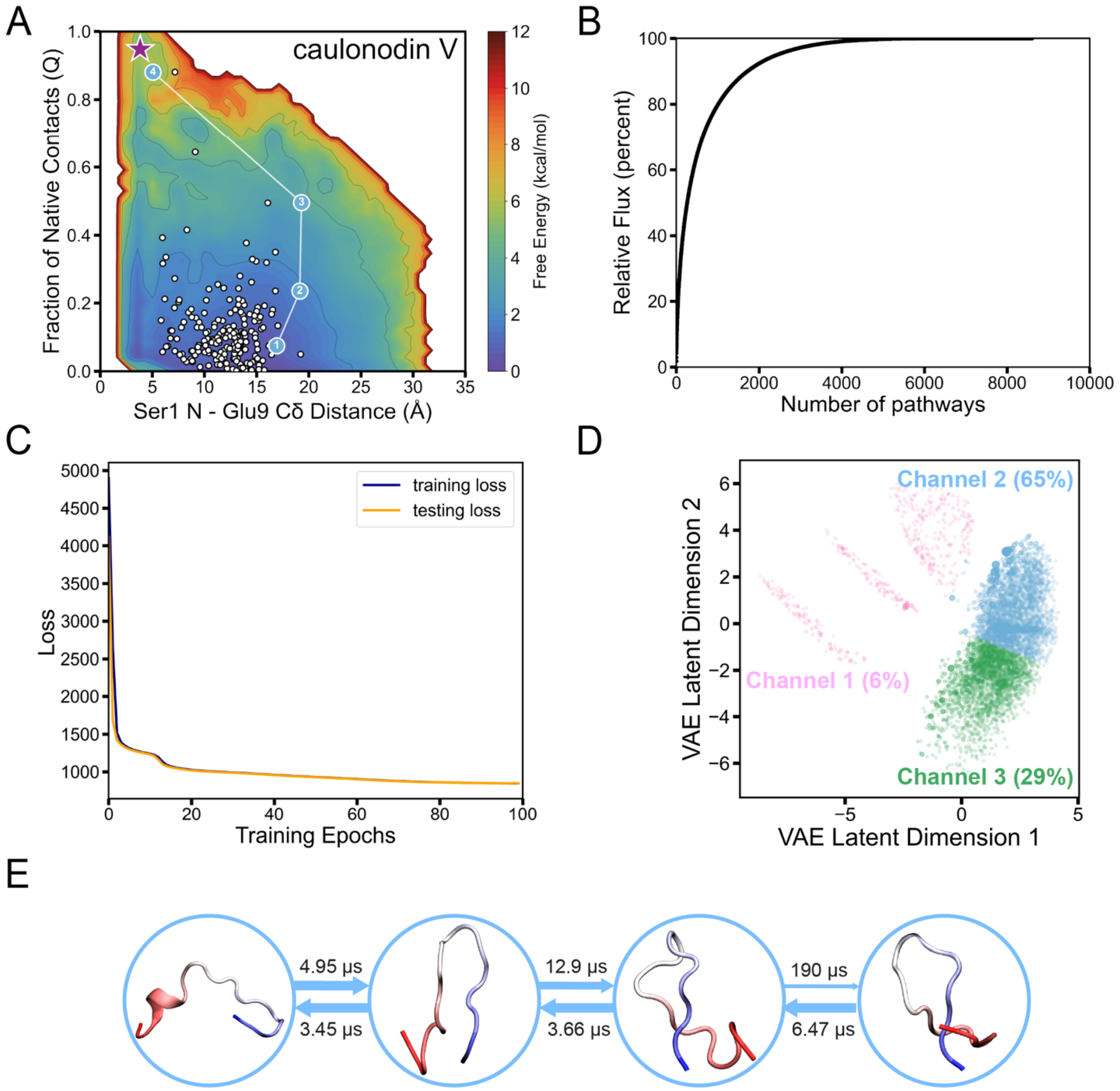

**Figure S18. Folding pathways of caulonodin V.** (A) Distribution of microstate centers (white dots) on the TRAM-weighted free energy landscape, with the most representative folding pathway highlighted by its corresponding pathway channel color. The pre-folded conformation is marked with a purple star. (B) Cumulative flux as a function of the number of kinetic pathways. (C) Loss function vs training epochs of VAE. (D) Kinetic pathways and pathway channels in the latent space captured by the VAE-based LPC algorithm. Each point in the latent space corresponds to a single pathway, and its size is determined by its normalized flux value. Each pathway channel was color-labeled with its relative percentage of the total flux. (E) Structural details of the most representative folding pathway, with the MFPT calculated between every two adjacent states. The N-terminus of the lasso peptide is shown in red, and the C-terminus of the lasso peptide is shown in blue.

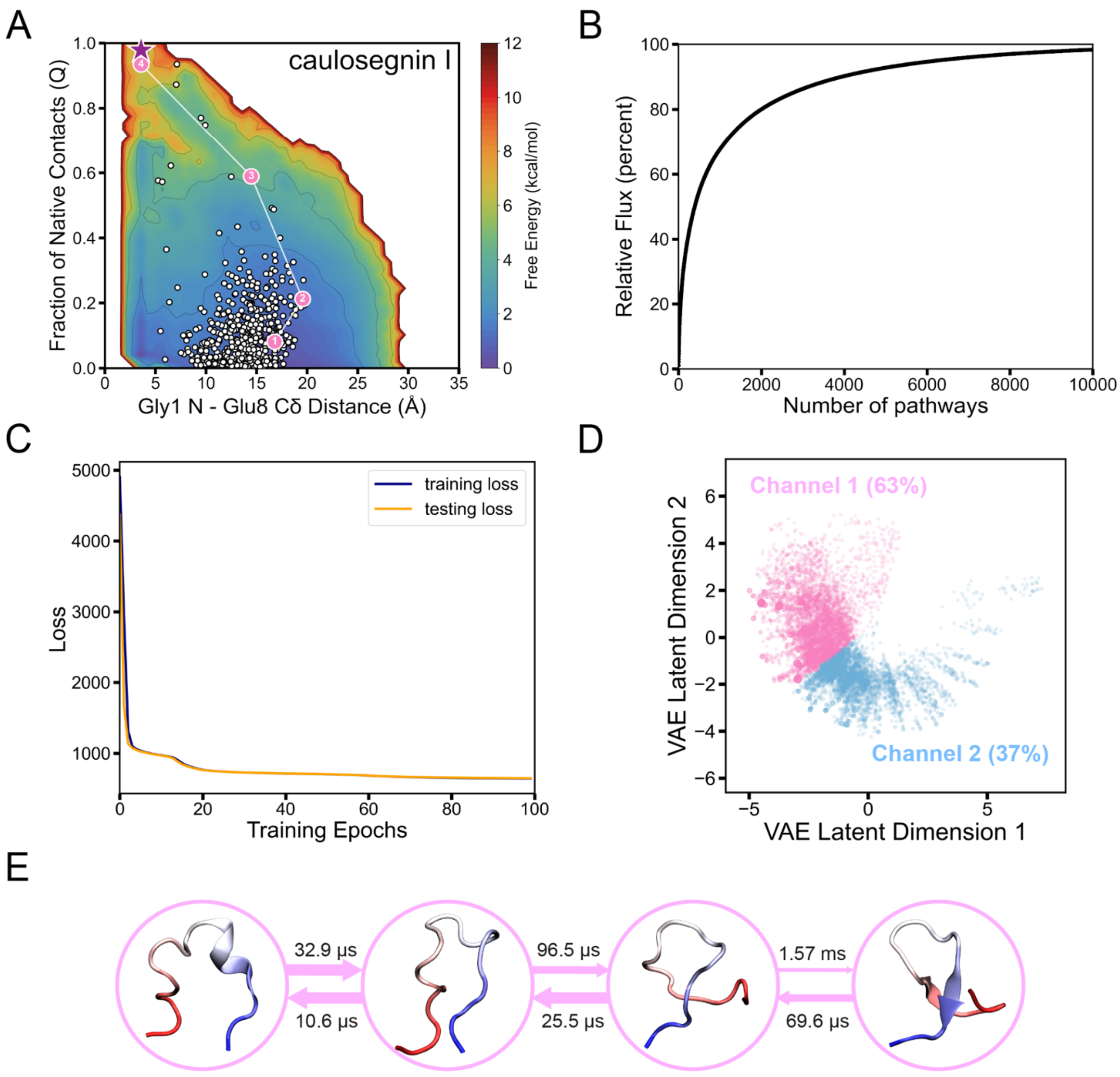

**Figure S19. Folding pathways of caulosegnin I.** (A) Distribution of microstate centers (white dots) on the TRAM-weighted free energy landscape, with the most representative folding pathway highlighted by its corresponding pathway channel color. The pre-folded conformation is marked with a purple star. (B) Cumulative flux as a function of the number of kinetic pathways. (C) Loss function vs training epochs of VAE. (D) Kinetic pathways and pathway channels in the latent space captured by the VAE-based LPC algorithm. Each point in the latent space corresponds to a single pathway, and its size is determined by its normalized flux value. Each pathway channel was color-labeled with its relative percentage of the total flux. (E) Structural details of the most representative folding pathway, with the MFPT calculated between every two adjacent states. The N-terminus of the lasso peptide is shown in red, and the C-terminus of the lasso peptide is shown in blue.

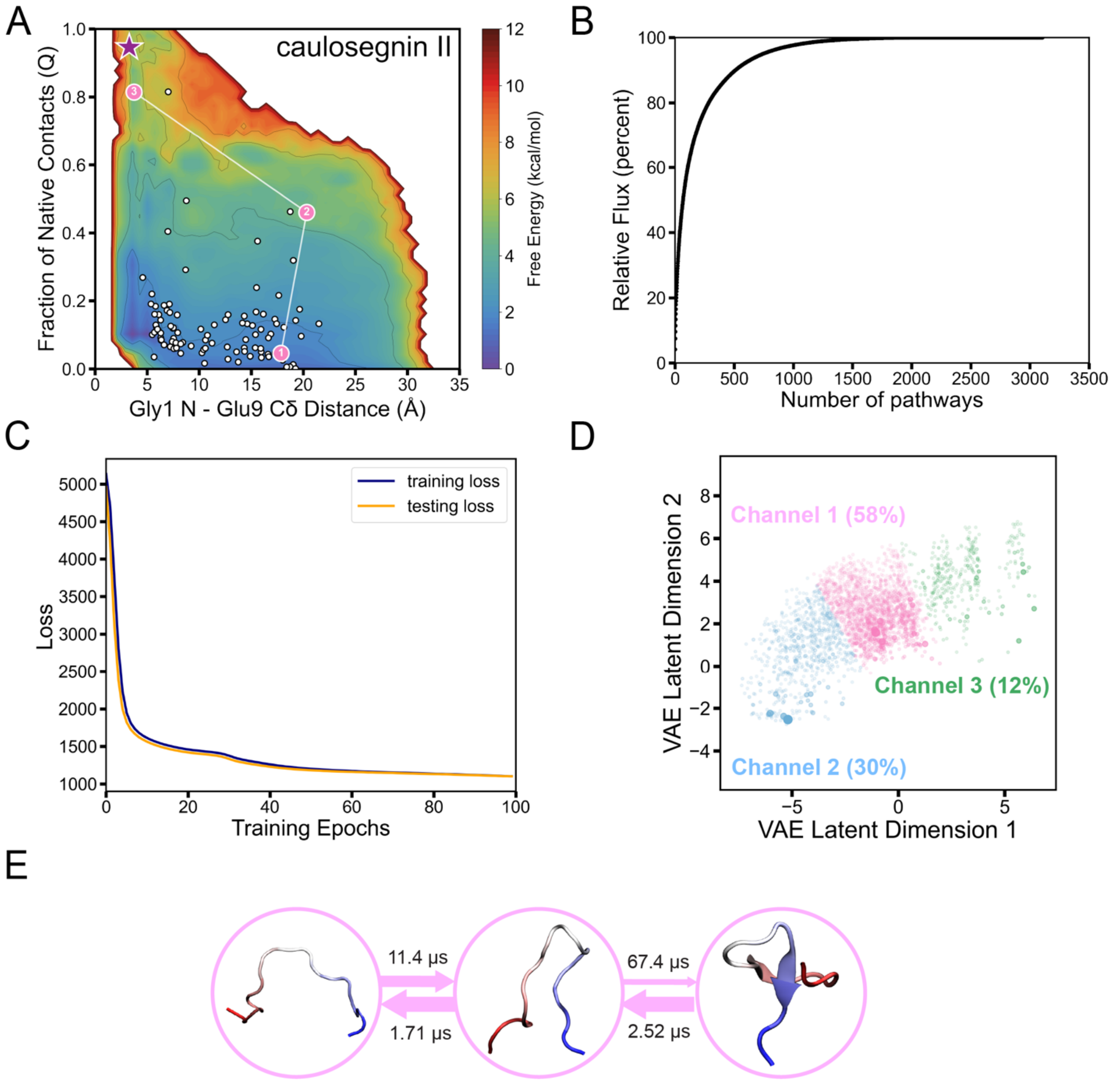

**Figure S20. Folding pathways of caulosegnin II.** (A) Distribution of microstate centers (white dots) on the TRAM-weighted free energy landscape, with the most representative folding pathway highlighted by its corresponding pathway channel color. The pre-folded conformation is marked with a purple star. (B) Cumulative flux as a function of the number of kinetic pathways. (C) Loss function vs training epochs of VAE. (D) Kinetic pathways and pathway channels in the latent space captured by the VAE-based LPC algorithm. Each point in the latent space corresponds to a single pathway, and its size is determined by its normalized flux value. Each pathway channel was color-labeled with its relative percentage of the total flux. (E) Structural details of the most representative folding pathway, with the MFPT calculated between every two adjacent states. The N-terminus of the lasso peptide is shown in red, and the C-terminus of the lasso peptide is shown in blue.

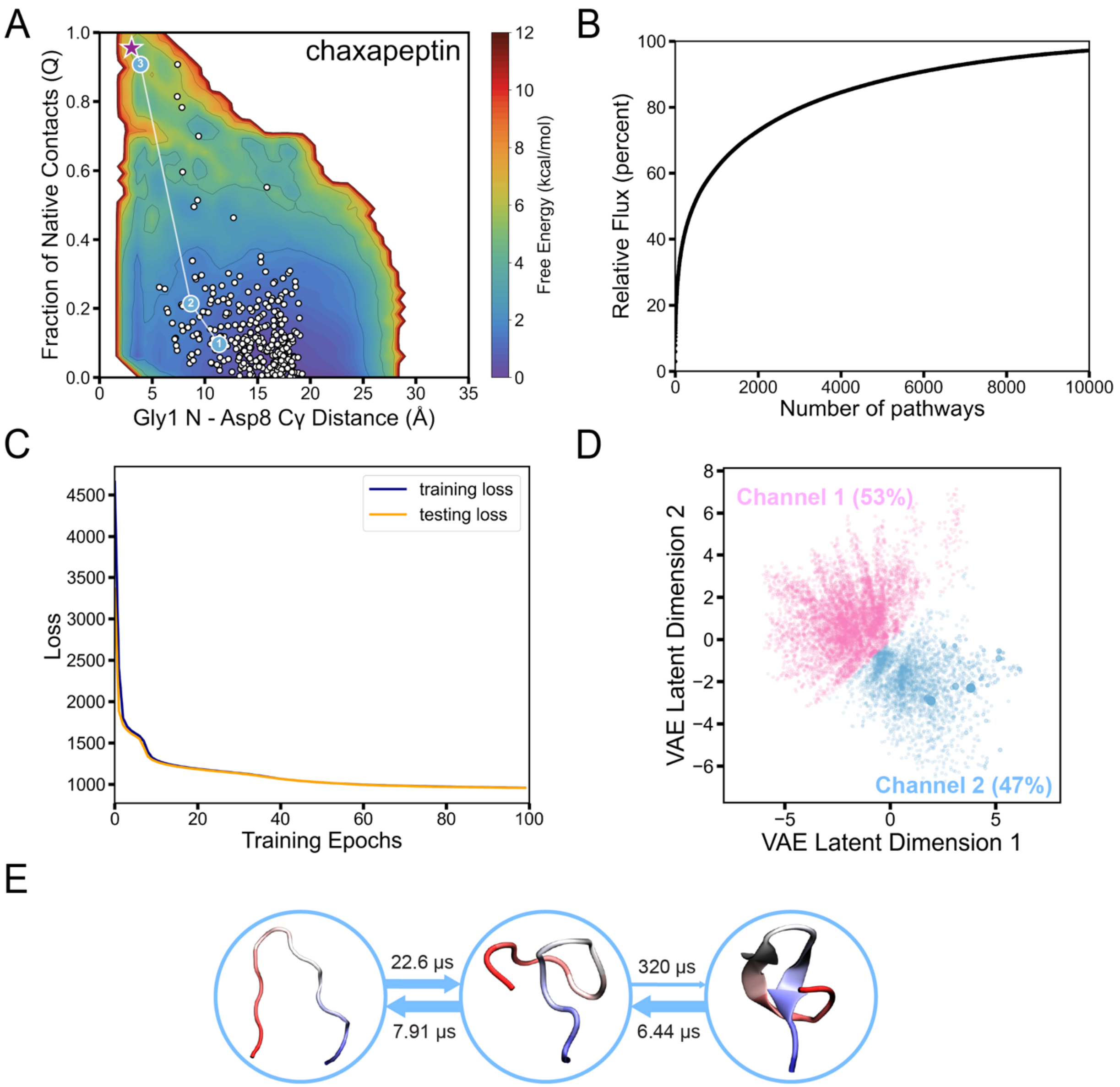

**Figure S21. Folding pathways of chaxapeptin.** (A) Distribution of microstate centers (white dots) on the TRAM-weighted free energy landscape, with the most representative folding pathway highlighted by its corresponding pathway channel color. The pre-folded conformation is marked with a purple star. (B) Cumulative flux as a function of the number of kinetic pathways. (C) Loss function vs training epochs of VAE. (D) Kinetic pathways and pathway channels in the latent space captured by the VAE-based LPC algorithm. Each point in the latent space corresponds to a single pathway, and its size is determined by its normalized flux value. Each pathway channel was color-labeled with its relative percentage of the total flux. (E) Structural details of the most representative folding pathway, with the MFPT calculated between every two adjacent states. The N-terminus of the lasso peptide is shown in red, and the C-terminus of the lasso peptide is shown in blue.

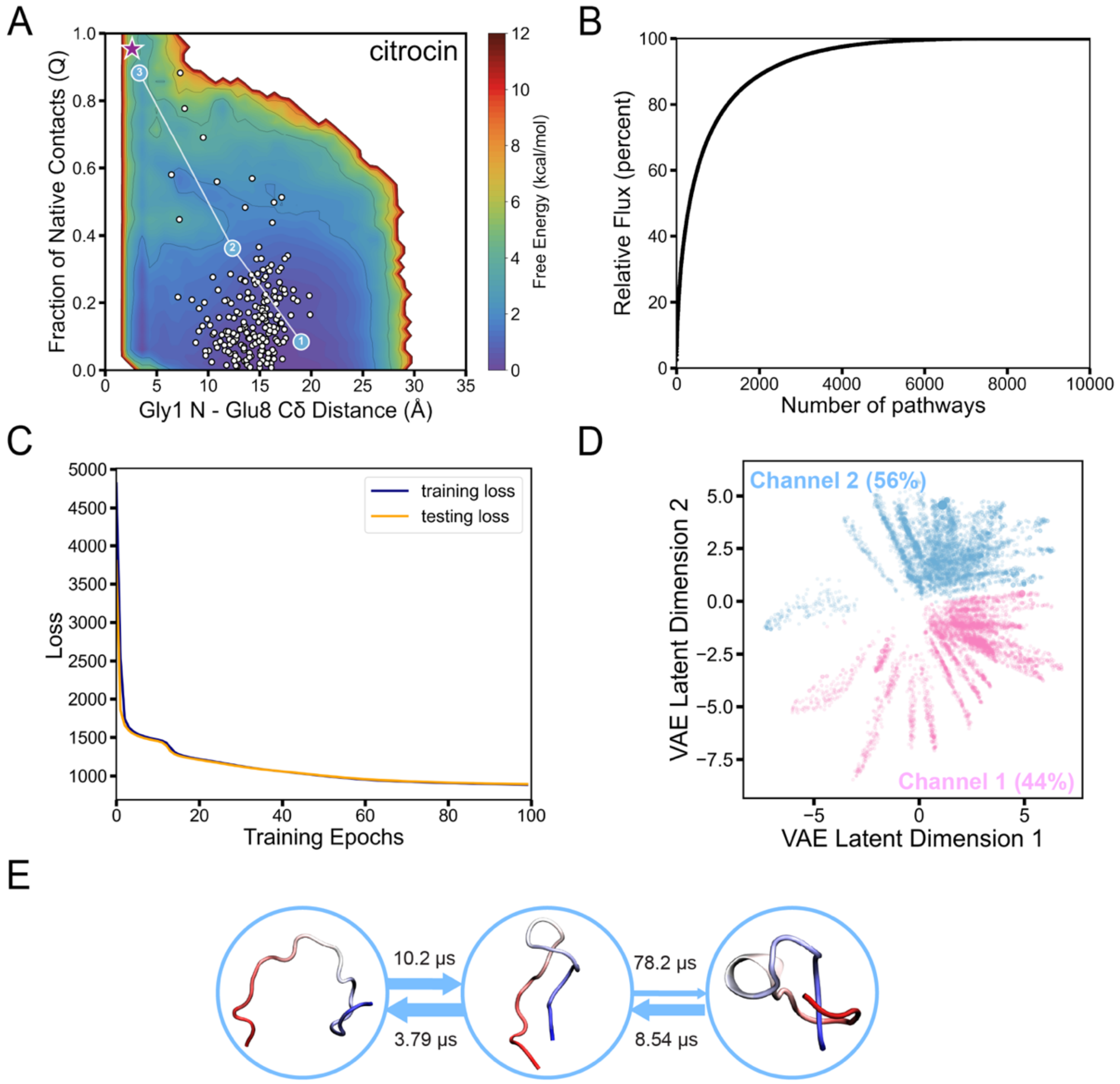

**Figure S22. Folding pathways of citrocin.** (A) Distribution of microstate centers (white dots) on the TRAM-weighted free energy landscape, with the most representative folding pathway highlighted by its corresponding pathway channel color. The pre-folded conformation is marked with a purple star. (B) Cumulative flux as a function of the number of kinetic pathways. (C) Loss function vs training epochs of VAE. (D) Kinetic pathways and pathway channels in the latent space captured by the VAE-based LPC algorithm. Each point in the latent space corresponds to a single pathway, and its size is determined by its normalized flux value. Each pathway channel was color-labeled with its relative percentage of the total flux. (E) Structural details of the most representative folding pathway, with the MFPT calculated between every two adjacent states. The N-terminus of the lasso peptide is shown in red, and the C-terminus of the lasso peptide is shown in blue.

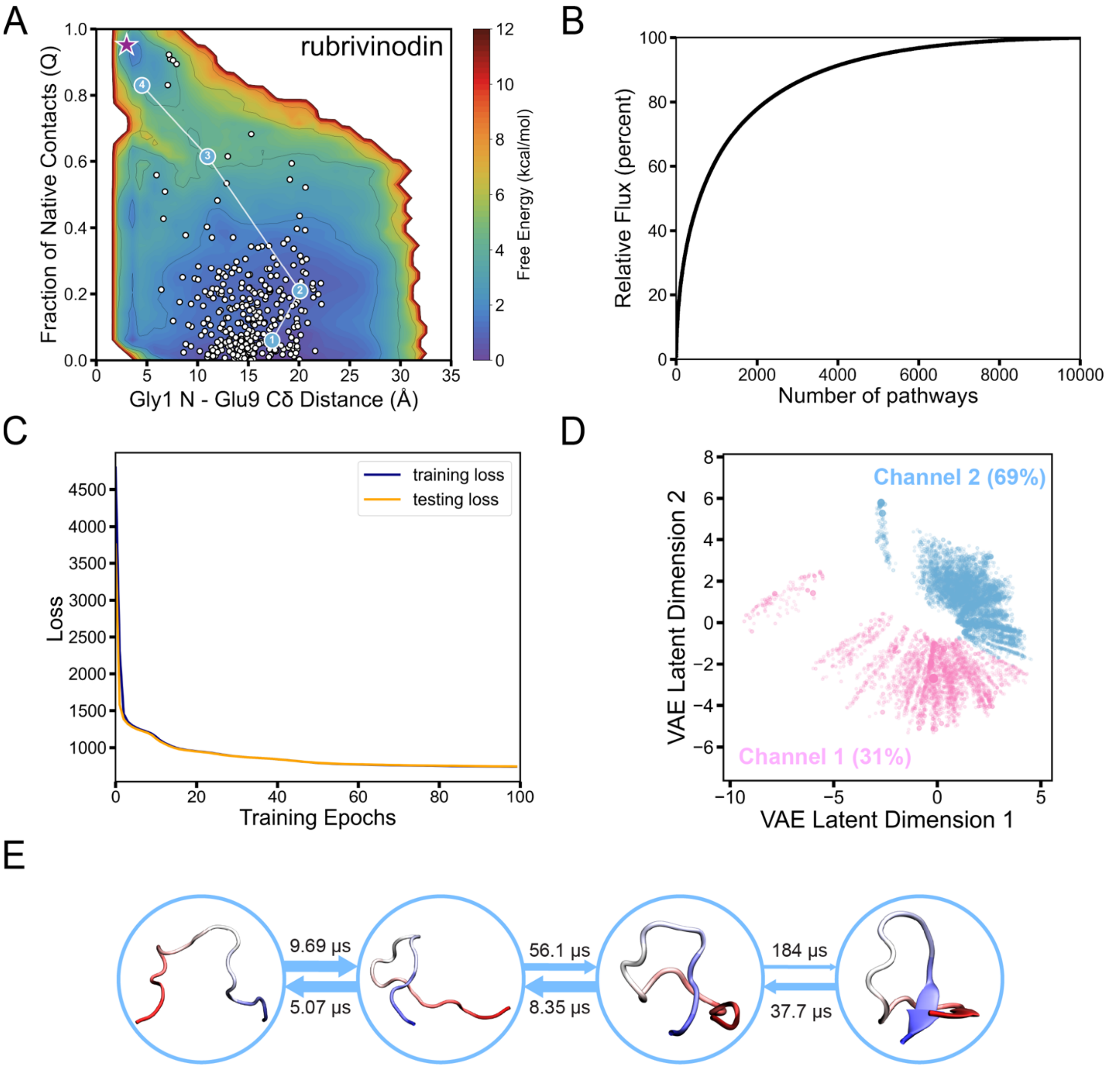

**Figure S23. Folding pathways of rubrivinodin.** (A) Distribution of microstate centers (white dots) on the TRAM-weighted free energy landscape, with the most representative folding pathway highlighted by its corresponding pathway channel color. The pre-folded conformation is marked with a purple star. (B) Cumulative flux as a function of the number of kinetic pathways. (C) Loss function vs training epochs of VAE. (D) Kinetic pathways and pathway channels in the latent space captured by the VAE-based LPC algorithm. Each point in the latent space corresponds to a single pathway, and its size is determined by its normalized flux value. Each pathway channel was color-labeled with its relative percentage of the total flux. (E) Structural details of the most representative folding pathway, with the MFPT calculated between every two adjacent states. The N-terminus of the lasso peptide is shown in red, and the C-terminus of the lasso peptide is shown in blue.

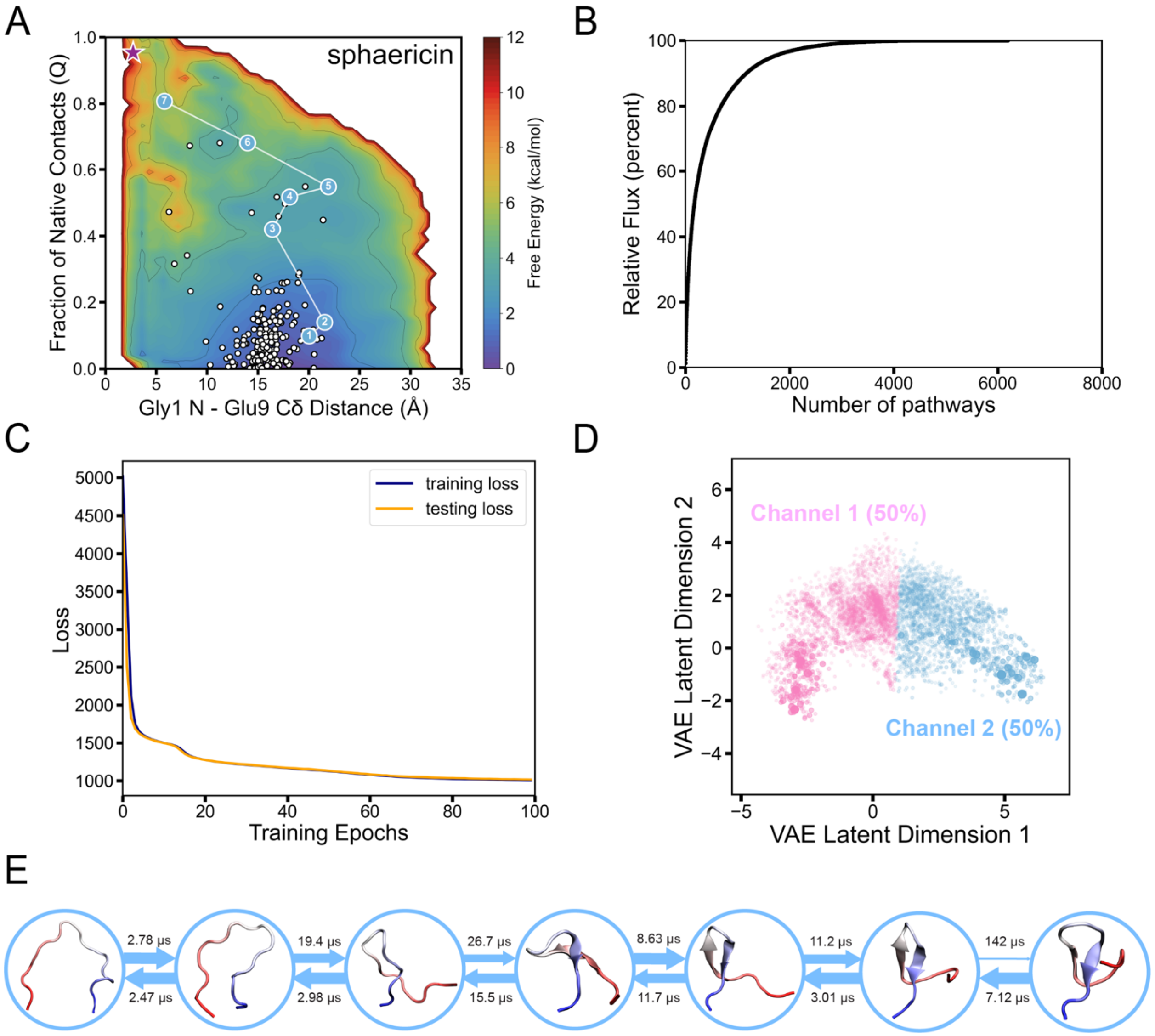

**Figure S24. Folding pathways of sphaericin.** (A) Distribution of microstate centers (white dots) on the TRAM-weighted free energy landscape, with the most representative folding pathway highlighted by its corresponding pathway channel color. The pre-folded conformation is marked with a purple star. (B) Cumulative flux as a function of the number of kinetic pathways. (C) Loss function vs training epochs of VAE. (D) Kinetic pathways and pathway channels in the latent space captured by the VAE-based LPC algorithm. Each point in the latent space corresponds to a single pathway, and its size is determined by its normalized flux value. Each pathway channel was color-labeled with its relative percentage of the total flux. (E) Structural details of the most representative folding pathway, with the MFPT calculated between every two adjacent states. The N-terminus of the lasso peptide is shown in red, and the C-terminus of the lasso peptide is shown in blue.

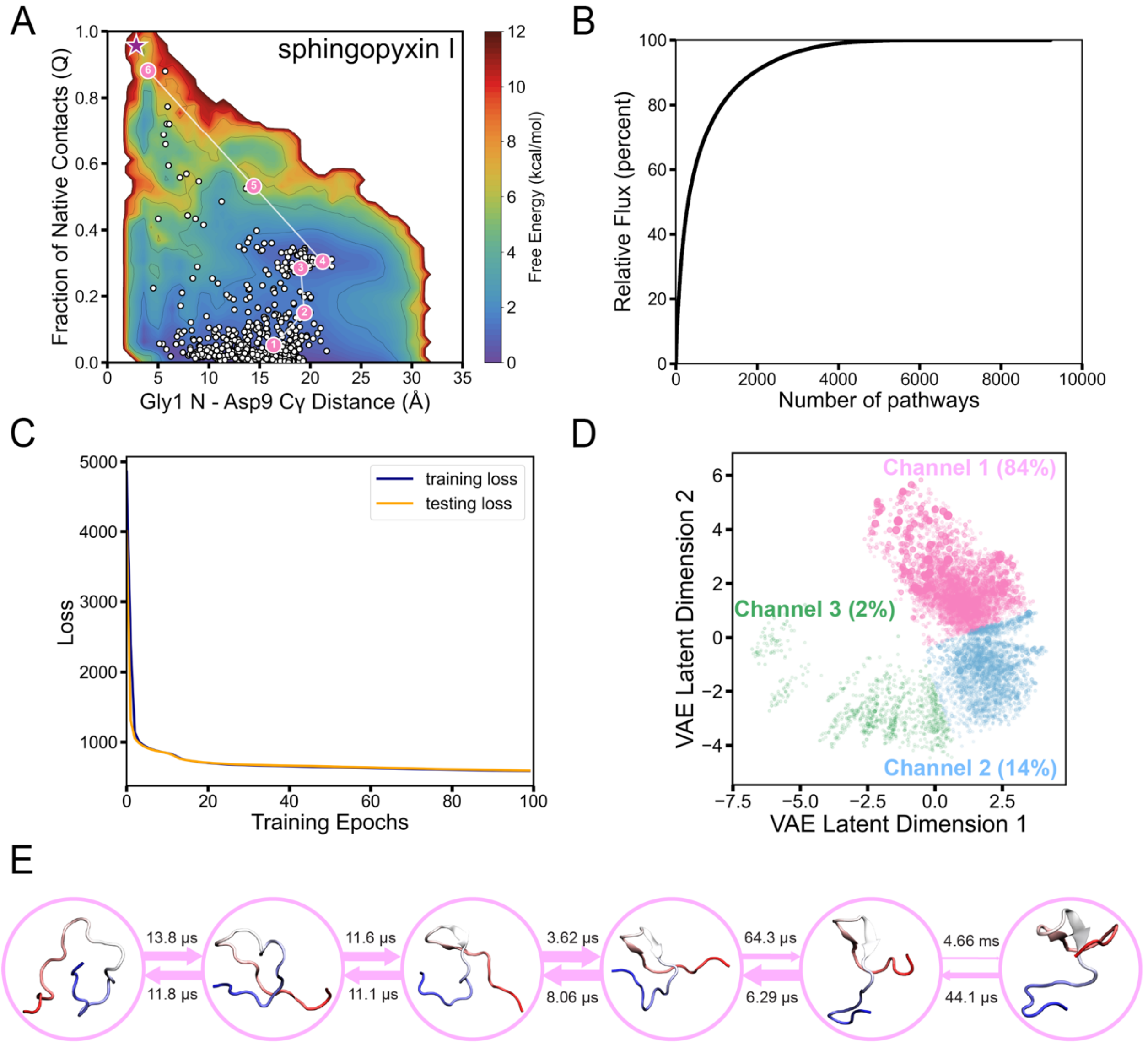

**Figure S25. Folding pathways of sphingopyxin-I.** (A) Distribution of microstate centers (white dots) on the TRAM-weighted free energy landscape, with the most representative folding pathway highlighted by its corresponding pathway channel color. The pre-folded conformation is marked with a purple star. (B) Cumulative flux as a function of the number of kinetic pathways. (C) Loss function vs training epochs of VAE. (D) Kinetic pathways and pathway channels in the latent space captured by the VAE-based LPC algorithm. Each point in the latent space corresponds to a single pathway, and its size is determined by its normalized flux value. Each pathway channel was color-labeled with its relative percentage of the total flux. (E) Structural details of the most representative folding pathway, with the MFPT calculated between every two adjacent states. The N-terminus of the lasso peptide is shown in red, and the C-terminus of the lasso peptide is shown in blue.

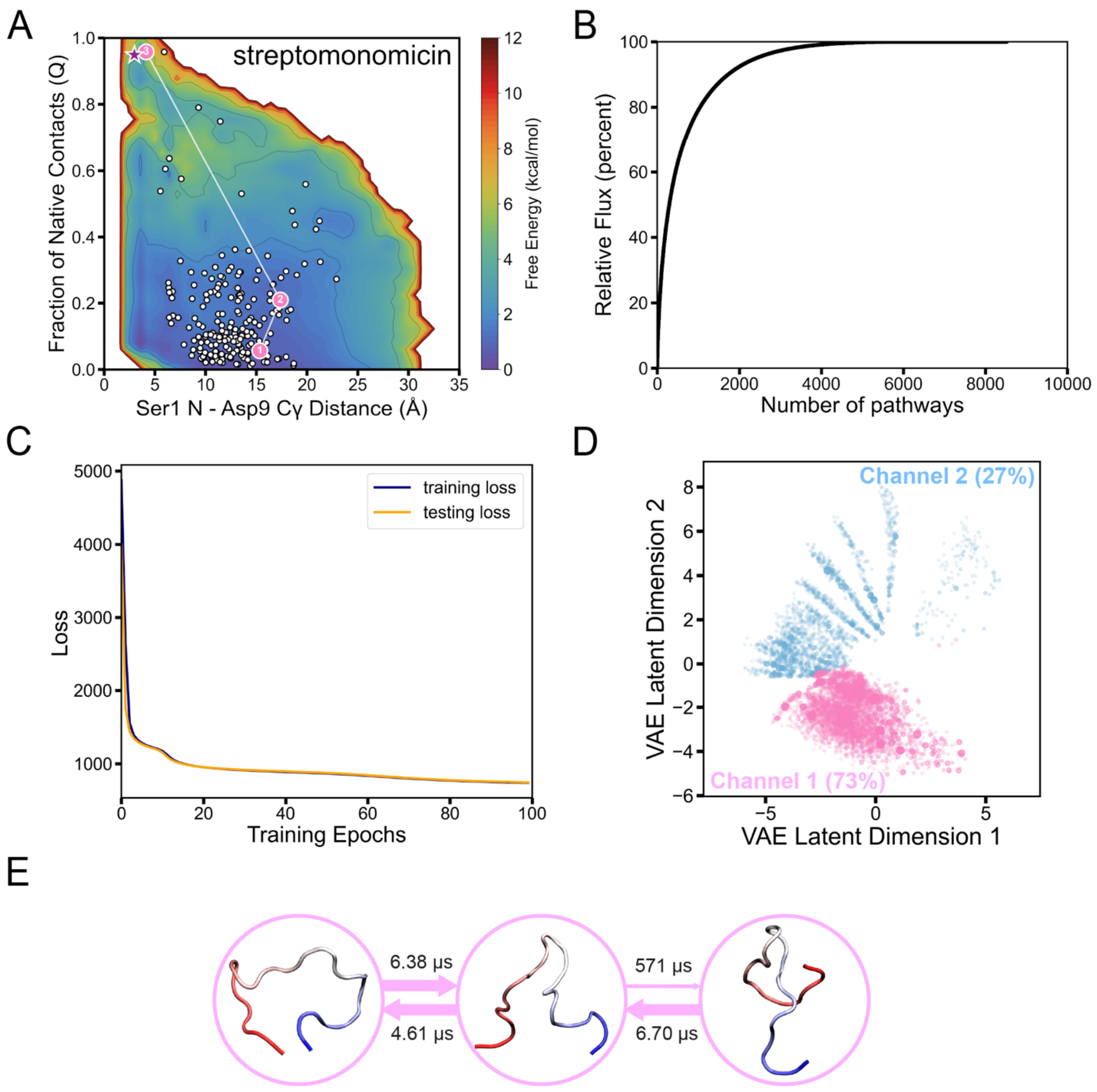

**Figure S26. Folding pathways of streptomonomicin.** (A) Distribution of microstate centers (white dots) on the TRAM-weighted free energy landscape, with the most representative folding pathway highlighted by its corresponding pathway channel color. The pre-folded conformation is marked with a purple star. (B) Cumulative flux as a function of the number of kinetic pathways. (C) Loss function vs training epochs of VAE. (D) Kinetic pathways and pathway channels in the latent space captured by the VAE-based LPC algorithm. Each point in the latent space corresponds to a single pathway, and its size is determined by its normalized flux value. Each pathway channel was color-labeled with its relative percentage of the total flux. (E) Structural details of the most representative folding pathway, with the MFPT calculated between every two adjacent states. The N-terminus of the lasso peptide is shown in red, and the C-terminus of the lasso peptide is shown in blue.

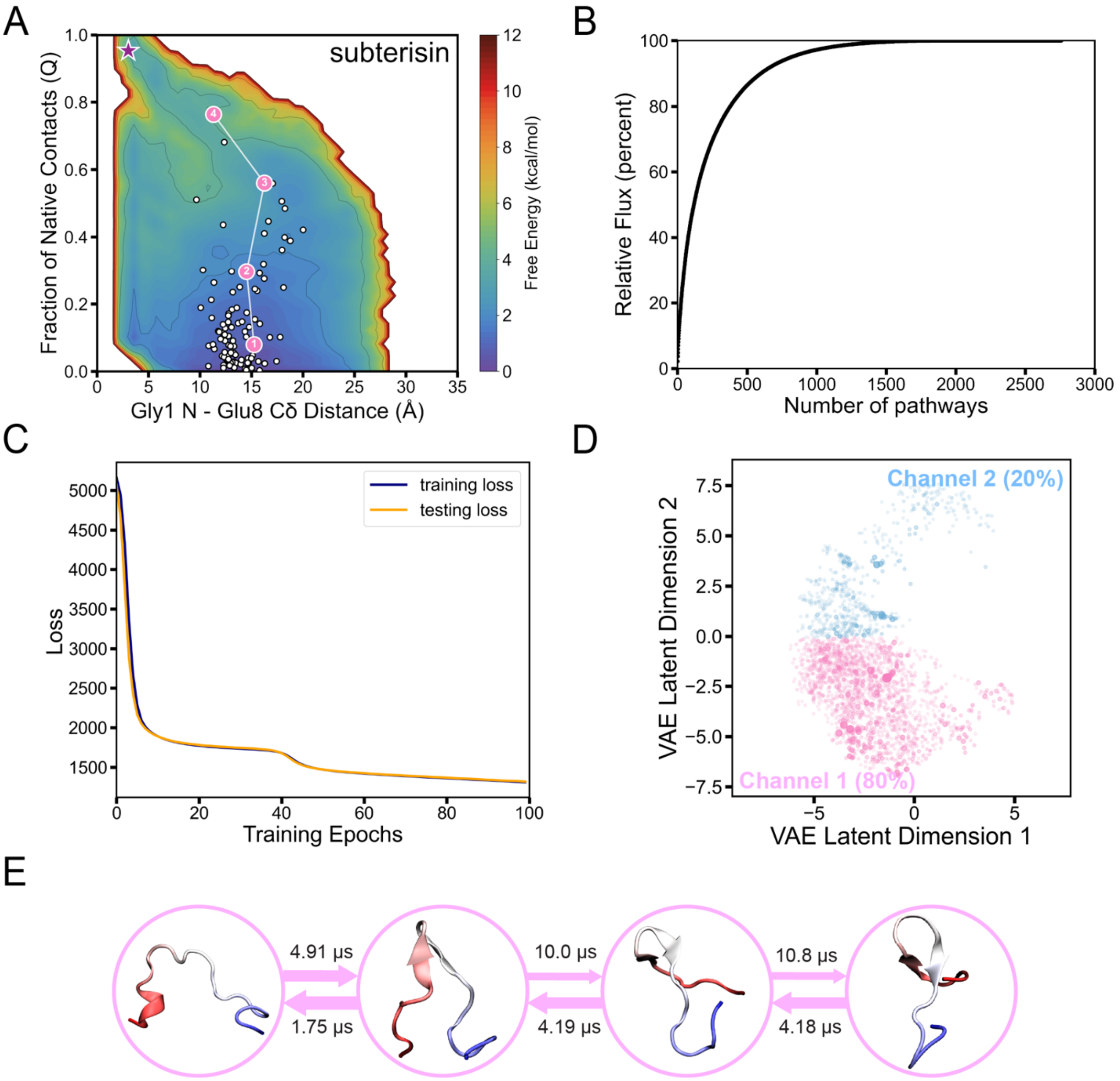

**Figure S27. Folding pathways of subterisin.** (A) Distribution of microstate centers (white dots) on the TRAM-weighted free energy landscape, with the most representative folding pathway highlighted by its corresponding pathway channel color. The pre-folded conformation is marked with a purple star. (B) Cumulative flux as a function of the number of kinetic pathways. (C) Loss function vs training epochs of VAE. (D) Kinetic pathways and pathway channels in the latent space captured by the VAE-based LPC algorithm. Each point in the latent space corresponds to a single pathway, and its size is determined by its normalized flux value. Each pathway channel was color-labeled with its relative percentage of the total flux. (E) Structural details of the most representative folding pathway, with the MFPT calculated between every two adjacent states. The N-terminus of the lasso peptide is shown in red, and the C-terminus of the lasso peptide is shown in blue.

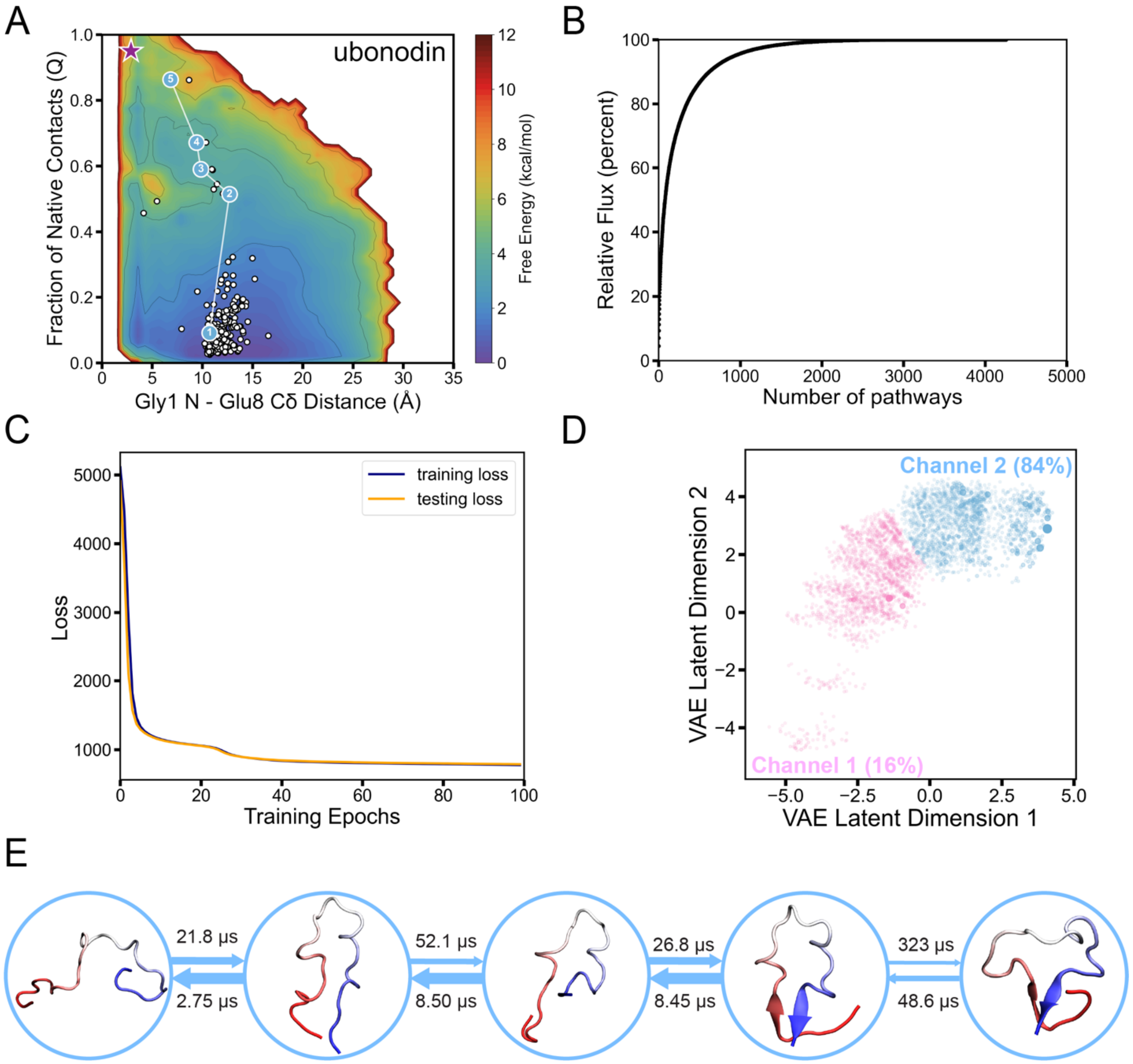

**Figure S28. Folding pathways of ubonodin.** (A) Distribution of microstate centers (white dots) on the TRAM-weighted free energy landscape, with the most representative folding pathway highlighted by its corresponding pathway channel color. The pre-folded conformation is marked with a purple star. (B) Cumulative flux as a function of the number of kinetic pathways. (C) Loss function vs training epochs of VAE. (D) Kinetic pathways and pathway channels in the latent space captured by the VAE-based LPC algorithm. Each point in the latent space corresponds to a single pathway, and its size is determined by its normalized flux value. Each pathway channel was color-labeled with its relative percentage of the total flux. (E) Structural details of the most representative folding pathway, with the MFPT calculated between every two adjacent states. The N-terminus of the lasso peptide is shown in red, and the C-terminus of the lasso peptide is shown in blue.

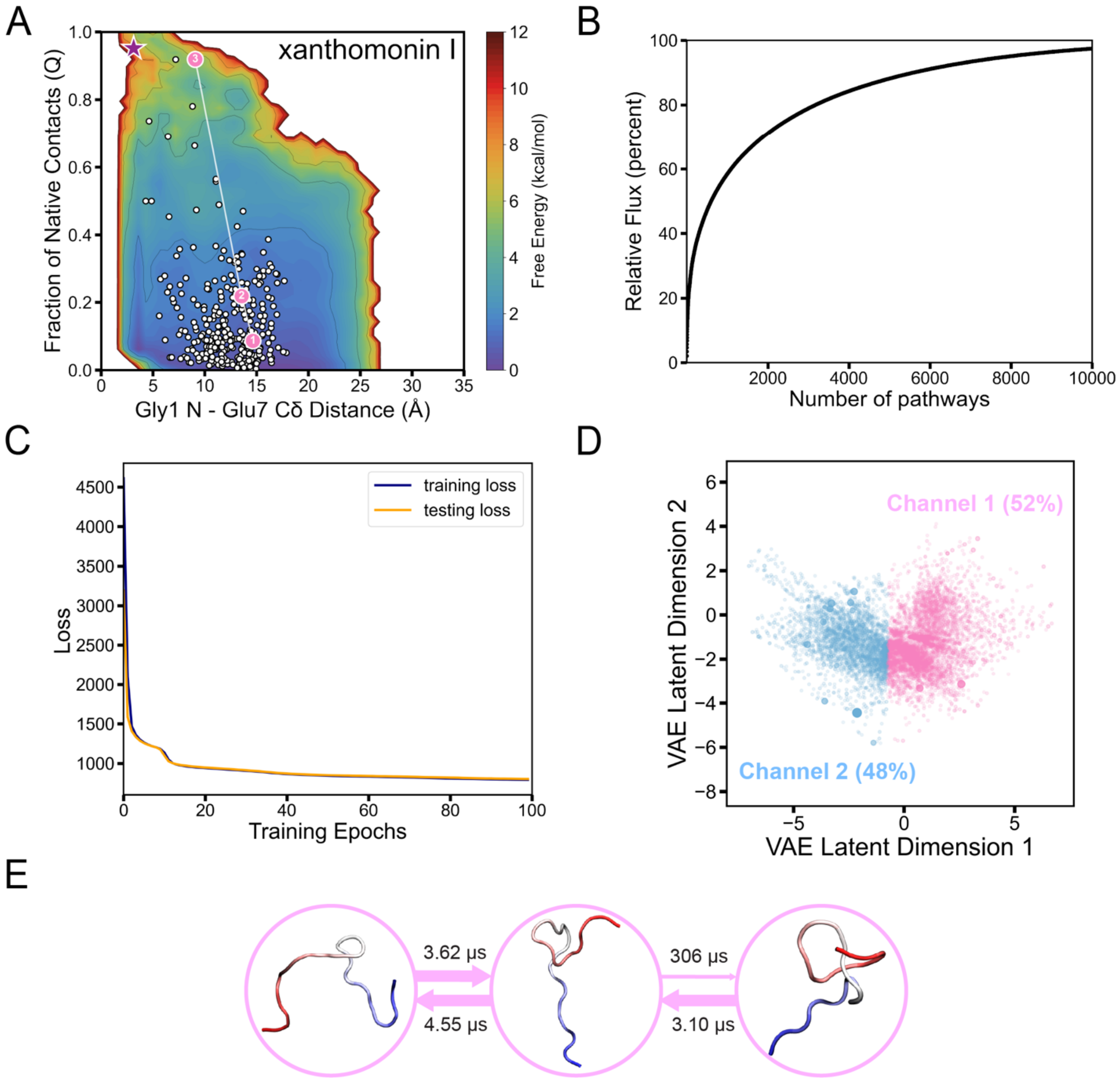

**Figure S29. Folding pathways of xanthomonin-I.** (A) Distribution of microstate centers (white dots) on the TRAM-weighted free energy landscape, with the most representative folding pathway highlighted by its corresponding pathway channel color. The pre-folded conformation is marked with a purple star. (B) Cumulative flux as a function of the number of kinetic pathways. (C) Loss function vs training epochs of VAE. (D) Kinetic pathways and pathway channels in the latent space captured by the VAE-based LPC algorithm. Each point in the latent space corresponds to a single pathway, and its size is determined by its normalized flux value. Each pathway channel was color-labeled with its relative percentage of the total flux. (E) Structural details of the most representative folding pathway, with the MFPT calculated between every two adjacent states. The N-terminus of the lasso peptide is shown in red, and the C-terminus of the lasso peptide is shown in blue.

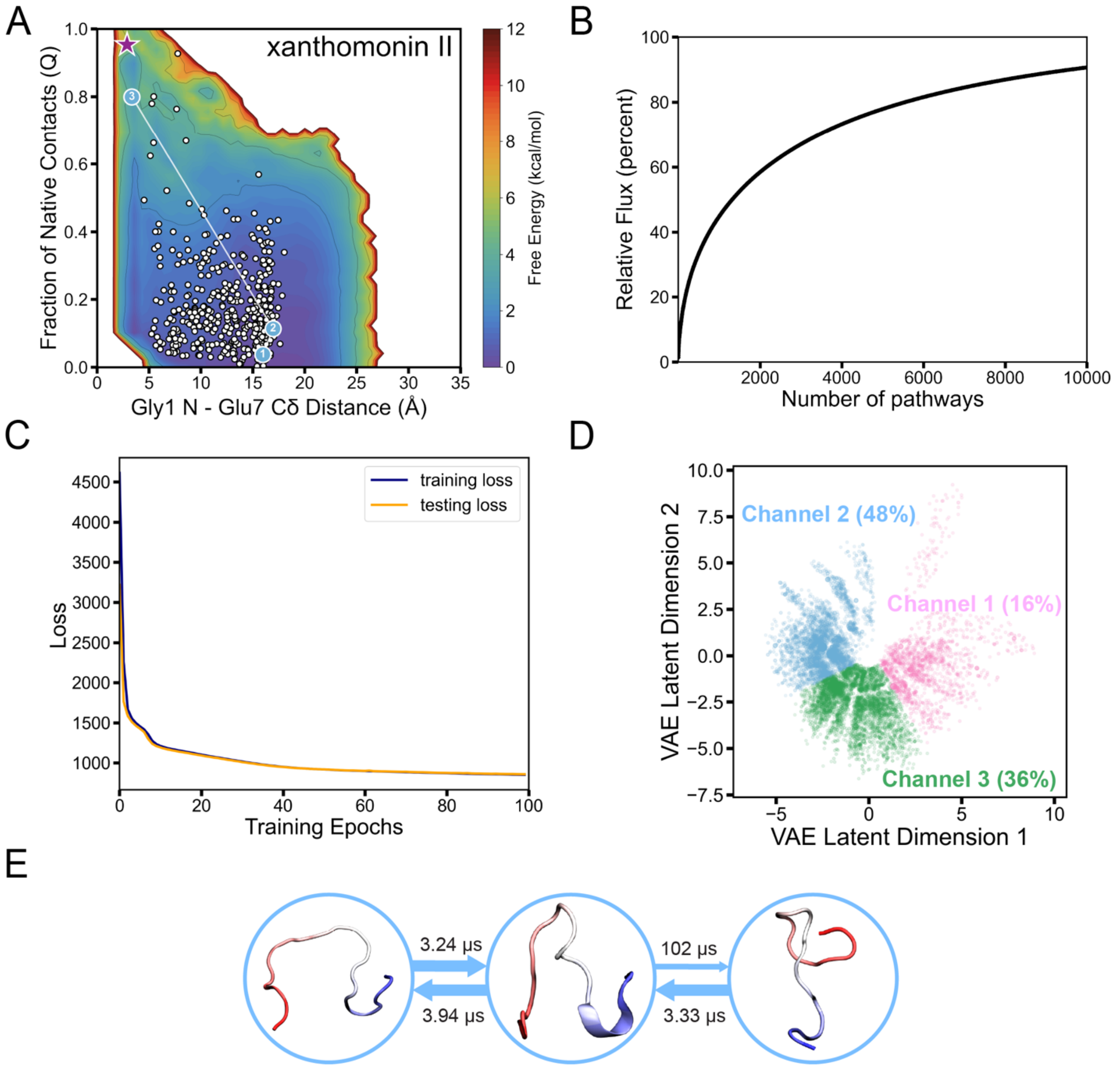

**Figure S30. Folding pathways of xanthomonin-II.** (A) Distribution of microstate centers (white dots) on the TRAM-weighted free energy landscape, with the most representative folding pathway highlighted by its corresponding pathway channel color. The pre-folded conformation is marked with a purple star. (B) Cumulative flux as a function of the number of kinetic pathways. (C) Loss function vs training epochs of VAE. (D) Kinetic pathways and pathway channels in the latent space captured by the VAE-based LPC algorithm. Each point in the latent space corresponds to a single pathway, and its size is determined by its normalized flux value. Each pathway channel was color-labeled with its relative percentage of the total flux. (E) Structural details of the most representative folding pathway, with the MFPT calculated between every two adjacent states. The N-terminus of the lasso peptide is shown in red, and the C-terminus of the lasso peptide is shown in blue.
